# Non-negative matrix factorization and deconvolution as dual simplex problem

**DOI:** 10.1101/2024.04.09.588652

**Authors:** Denis Kleverov, Ekaterina Aladyeva, Alexey Serdyukov, Maxim N. Artyomov

## Abstract

Non-negative matrix factorization (NMF) is one of the most powerful linear algebra tools, which has found application in various areas of data analysis, including computational biology. Despite numerous optimization methods devised for NMF, our comprehension of the inherent topological structure within factorizable matrices remains limited. In this work, we reveal the topological properties of the linear mixture data, which allow for a remarkable reduction in the dimensionality of the NMF problem and reformulation of the NMF problem as an optimization problem with only *K*(*K* −1)variables, with K representing the number of pure components, irrespective of the initial data matrix dimensionality. This is achieved by uncovering the dual simplex structure of the data, with complementary simplex structures existing in both the features’ and samples’ spaces and leveraging the Sinkhorn transformation to uncover the relationship between these simplexes. We validate this approach in the context of an unconstrained general mixed images scenario and achieve a significant improvement in decomposition accuracy. Furthermore, we successfully apply the proposed approach in the biological context of bulk RNA-seq gene expression data unmixing and single-cell RNA-seq data clustering.

## Introduction

Nonnegative matrix factorization (NMF) (D. D. Lee & Seung, 1999) is widespread in variety of computational biology problems, including genomics (Jin et al., 2024), transcriptomics (Elosua-Bayes et al., 2021; Rodriques et al., 2019), mass spectrometry (Nijs et al., 2021), and cytometry (Jiménez-Sánchez et al., 2020) etc. It is also well-known beyond biological sciences as a common technique for feature extraction (S. Lee & Pang, 2020) and data compression (Yuan & Oja, 2005). NMF seeks to represent a non-negative *N* ×*M* matrix *V* as a product of two non-negative matrices *W* and *H* (*V* = *W* ×*H*). While the data matrix *V* is typically high-dimensional (*N, M* ∼100-1000s), its rank (*K*) is often much smaller in biological settings (∼10s). The rank (*K*)of matrices *W* and *H* is usually thought of as the number of independent pure components/clusters/features comprising the dataset. In these terms, *W* is the *feature matrix*, representing the pure components of the data, and *H* is the *coefficients matrix*, representing the contributions of the pure components for each individual mixture.

One of the most common applications of the NMF is the linear unmixing/deconvolution problem, where matrix *H* is a proportion matrix, i.e. sum-to-one normalized (Miao & Qi, 2007). This constrained problem formulation is also widespread in modern computational biology –spanning the analysis of spectral cytometry data (McKinnon, 2018), spatial expression data analysis (Lundberg & Borner, 2019) and the dissection of composition of bulk transcriptional profiles (Gaujoux & Seoighe, 2013). Over the last decade, multiple computational approaches have been implemented to solve the deconvolution problem, especially in the context of gene expression, ranging from support vectors machines (Frishberg et al., 2019; Newman et al., 2015) to deep neural networks (Menden et al., 2020; Torroja & Sanchez-Cabo, 2019). While these methods often employ sophisticated computational techniques, no unified understanding of the underlying data structure for factorizable matrices exists at the moment. Indeed, even though NMF or deconvolution problems are seeking low-dimensional representation relative to dimensionality of the input data matrix, there is no explicit mathematical procedure that could rigorously leverage this notion in the factorization procedure.

In this work, we introduce an analytical framework that reveals dual/complementary simplexes within the features and samples spaces and then explicitly describe their relationship. We show that this can be achieved analytically by using projective formulation of the factorization/deconvolution problem for the Sinkhorn transformed non-negative matrix. By doing so, we first reduce NMF problem of finding *W* and *H* to finding two linked (*K* − 1) -dimensional simplexes. Furthermore, we then show that this problem is equivalent to finding only one (*K* − 1) - dimensional simplex with positivity constraints on it and its inverse, therefore resulting in formulation of the arbitrary NMF problem as optimization task with only *K*(*K* −1)independent variables.

The approach can be applied to both normalized and non-normalized non-negative matrices, and we illustrate its applicability by performing image unmixing, single-cell RNA-seq data clustering and complete deconvolution of the bulk transcriptional profiles. These findings advance our understanding of non-negative matrix factorization problem and provide unique insight into topological structures within the non-negative factorizable matrices. Practically, such dramatic reduction in dimensionality significantly simplifies computational task at hand and provides guidance for the best design for partial, signature-based, and complete deconvolution problems, as well as broader NMF problems such as image unmixing/decomposition and spatial data analysis.

### Complementary features and samples simplex structures

Let’s consider the non-negative matrix factorization problem *V* = *W* ×*H*. In general terms, non-negative factorizable matrix *V* can be viewed as a collection of *N* (e.g., number of samples) dots in an *M*(e.g., number of genes) dimensional space. We will call this space a *features space* since individual axes correspond to the values of the features (**Fig. 1a**). In turn, *V*^*T*^ can be viewed as collection of *M*dots (each representing individual features) in an *N*-dimensional space. We will call this space a *samples space* (**Fig. 1a**). In the most general case, when matrix *V* is neither row- nor column-normalized, matrices *V* and *V*^*T*^ represent unorganized collections of points in their corresponding spaces (**Fig. 1a**).

**Figure 1.**
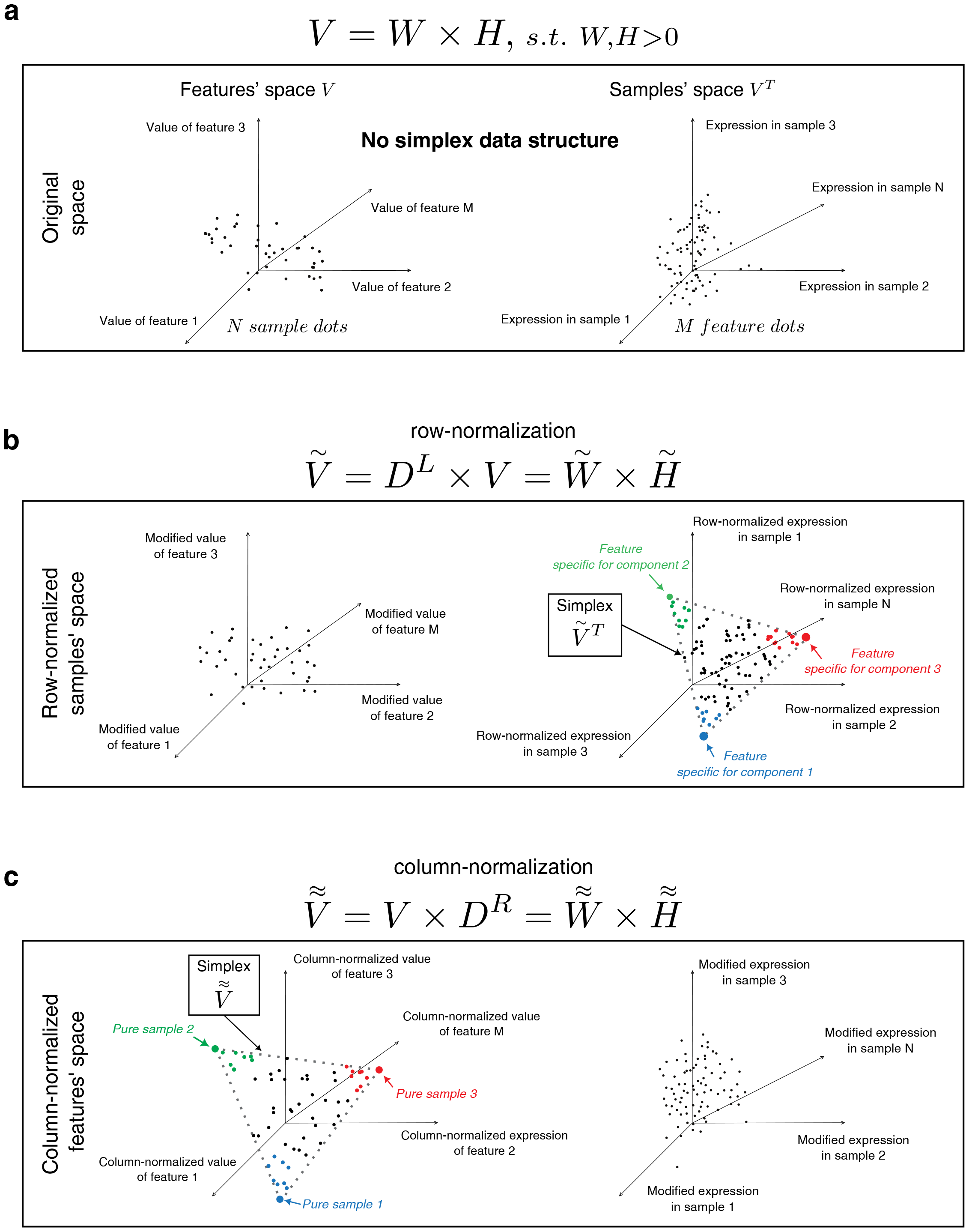
Existence of complementary features and samples simplex structures for normalized matrices. (a) Schematic representation of the data matrix in features and samples spaces. (b) Row normalization and transposition of original matrix aligns feature dots within a simplex in a samples space (c) Column normalization aligns sample dots within a simplex in features space

It has been observed that row normalization procedure applied to any positive and factorizable matrix leads to the emergence of a simplex structure in the space of its columns (Chan et al., 2008). Specifically, the row-normalized and transposed matrix 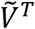 places all feature-dots within the (*K* −1)-dimensional simplex inside the *N*-dimensional space (**Fig. 1b**, Supplementary Note 1). In fact, the vertices of this simplex correspond to features specific to the pure samples, i.e. rows of matrix 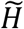 (**Fig. 1b**). More generally, the closer a feature-point is to a corner, the more specific it is for the given component (**Fig. 1b**). Accordingly, finding simplex corners in the samples space is equivalent to solving the deconvolution problem via finding component-specific feature sets (e.g., (Bioucas-Dias, 2009)). We and others demonstrated the application of this approach to the gene expression data deconvolution (Wang et al., 2016; Zaitsev et al., 2019). For example, in the case of a mixture of 3 types, this results in a 2-dimensional simplex (i.e., triangle) embedded in the multidimensional space of all the samples (Zaitsev et al., 2019).

On the other hand, column normalization of any positive and factorizable matrix reveals the simplex structure in the complementary samples space. Specifically, column normalization of the factorizable matrix *V* results in the matrix 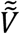 that describes a (*K* − 1) -dimensional simplex in an *M*-dimensional space (**Fig. 1c**, Supplementary Note 2 for proof). Indeed, 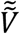 can be viewed as a collection of *N* dots (each representing an individual sample) in the *M*-dimensional space of features (**Fig. 1c**). Note that this space is the same as the *features space* in the non-transformed matrix since axes of this space correspond to the row-normalized values of each feature. However, following the transformations, the positions of the sample dots have changed, acquiring a new geometrical structure (**Fig. 1a, c**). In the context of gene expression deconvolution, the vertices of this simplex correspond to the transcriptional profiles of pure samples, i.e. columns of the matrix 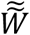 (Supplementary Note 2). Therefore, finding simplex corners in this space is also equivalent to solving the deconvolution problem.

Broadly speaking, the existence of these two simplex structures reflects two different approaches to solving the NMF problem: either by finding simplex corners in the feature space or in the samples space (**Fig. S1**). As we demonstrate below, there exists a connection between these simplexes which provides us with a unique and powerful approach to solving the NMF/deconvolution problem within a subspace of a significantly lower dimensionality.

### Sinkhorn transformation of data matrix links features and samples simplexes

First, let us remark that since column normalization leads to a simplex for any factorizable matrix (**Fig. 1**), the transposition and column normalization of the previously transformed matrix 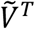 also results in a matrix, which forms a simplex in a *features space*. We can formalize this transformation using sequential multiplication by diagonal normalization matrices alternating between left and right multiplication (Eq. 1):

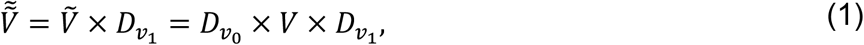

Where 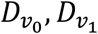 are diagonal matrices: 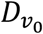 is a row-normalizing matrix for *V* and 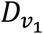 is a column normalizing matrix for the row normalized matrix 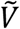.

Next, we note that subsequent row normalization applied to matrix 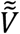 (Eq. 1) will result in a matrix 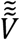 representing a simplex in a *samples space* which will be different from the previous simplex in the *samples space* obtained in the first step, i.e. 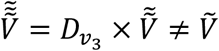. The next iteration will yield another simplex structure in the *features space* etc. In essence, the consecutive application of row normalization followed by column normalization conserves the simplex structures in their respective spaces. Such iterative normalization process is also known as *Sinkhorn transformation* of matrix (Sinkhorn, 1967, 1974) (**Fig. 2a**). Notably, according to the Sinkhorn-Knopp theorem (Sinkhorn, 1967), odd and even elements of this sequence will converge towards two related matrices 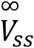 (the row-normalized matrix of the *samples space* simplex) and 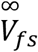 (the column-normalized matrix of the *features space* simplex):

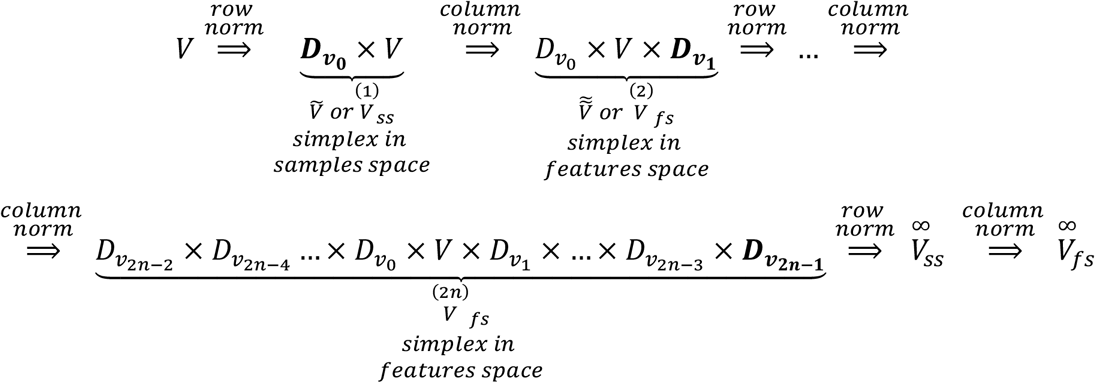

Furthermore, elements of these matrices will be proportional to each other (Sinkhorn, 1967). In the case of an initial *M*× *N* matrix *V*, the proportionality constant is *M*/*N* (**Fig. 2b**, Supplementary Note 3):

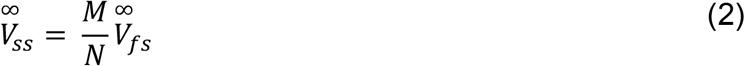

This convergence can be illustrated in a simulation scenario, where normalization is applied to the product of two randomly generated matrices *W* and *H* (*M*= 2000, *N* = 2000, *K* = 3) (**Fig. 2c left**). While single row or column normalization results in a single simplex structure within samples and features space respectively, the Sinkhorn procedure of as few as 3 iterations, reveals the simplex structures within both spaces simultaneously (**Fig. 2c right**).

**Figure 2.**
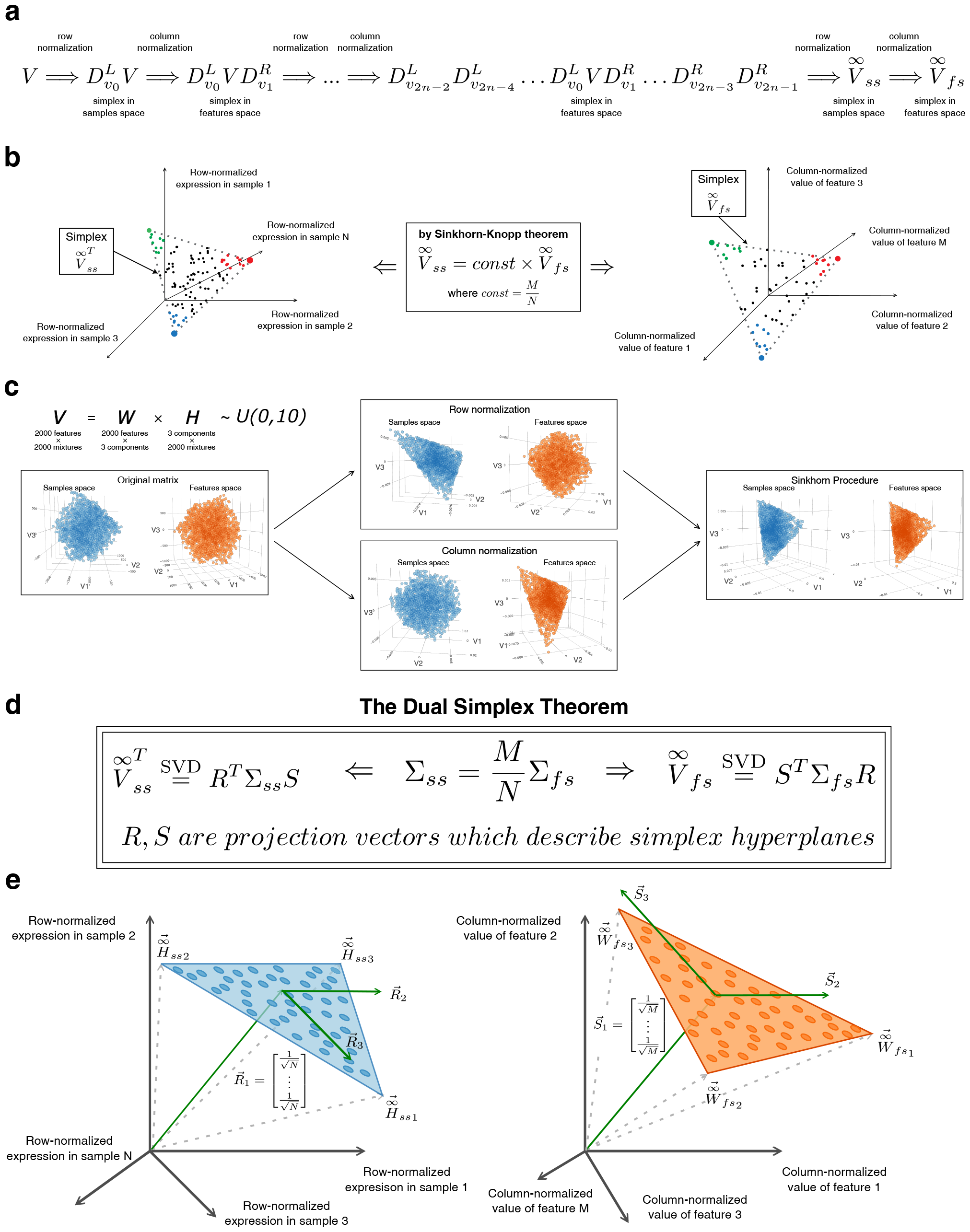
Sinkhorn transformation enables identification of the simplex hyperplane projection vectors. (a) Sinkhorn transformation is a process of iterative left and right multiplication by diagonal matrices (i.e. row and column normalization), producing two converging sequences of matrices, representing simplexes in features and samples spaces. (b) Converged matrices in features and samples spaces are proportionally related. (c) Example of Sinkhorn transformation applied to randomly generated factorizable matrix (K=3,M=1000,N=1000). While original spaces of features and samples does not contain simplex structures (left), column and row normalizations reveal simplex structure for one of the spaces (middle). Iterative normalizations applied to the matrix align points withing simplexes for both spaces (right). (d) The Dual Simplex theorem formulation. Singular vectors of Sinkhorn-transformed matrices provide projection vectors to hyperplanes in which samples and features simplexes are located. (e) Corollary of the Dual Simplex theorem: due to normalization constraint R_1 and S_1 have fixed form and represent the shift of the entire simplex hyperplane from the zero point.

### Sinkhorn transformation properties enable simultaneous identification of dual simplex hyperplanes

We will now consider how corresponding simplexes can be identified within 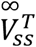 and 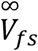. Locating the data simplex generally involves two fundamental steps: first, finding a lower dimensional hyperplane wherein the simplex is located, and then proceeding to subsequent identification of the simplex vertices within this hyperplane (**Fig. S1**). We find that Sinkhorn transformation enables a very elegant identification of the hyperplanes in which simplexes are confined.

To demonstrate this, we will first consider independent problems of hyperplanes identification for features and samples simplexes and then describe the relationship between them. Broadly, considering that the rank of the matrix *V* is *K*, we approach the task of hyperplane identification as a problem of searching a projection onto the corresponding *K*-dimensional subspace for either row-normalized and transposed matrix 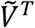 or the column-normalized matrix 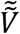. One can utilize a projective NMF framework searching for respective projection operators *ProjR* = *R*^*T*^*R* and *ProjS* = *S*^*T*^*S* (Yuan & Oja, 2005). Here, *R* is a (*K* × *N)* –matrix comprising orthogonal vectors that define the coordinate system within a projected subspace of the features space, and *S* is a (*K* × *M)* –matrix, defining the coordinate system within a projected subspace of the samples space (Eq. 3 a, b, **Fig. S2a**, Supplementary Note 4, 5).

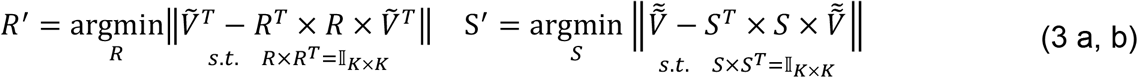

This *K*-dimensional optimization problem at hand is recognized to be equivalent to singular value decomposition and can be solved for any non-negative matrix *V* of rank *K* (Song & Ng, 2020).

However, the formulation presented in (Eq. 3 a, b) does not directly identify the hyperspace containing the (*K* − 1) -dimensional simplex, as singular vectors, in general, may not align precisely with this hyperspace (**Fig. S2**). Ideally, we aim to identify projection operator that would act on 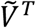 or 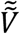 and yield a (*K* − 1) -dimensional simplex. This decrease in dimensionality stems from an additional sum-to-one constraint which is a consequence of normalization procedure. In principle, one can identify such a (*K* − 1) -dimensional structure embedded in a *K*-dimensional space by initially getting a centered data matrix 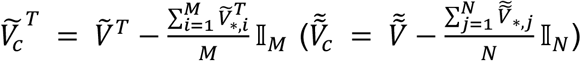 and then proceeding with finding of the (*K* − 1) -dimensional singular value decomposition (SVD) of the centered matrix (Chan et al., 2008) (**Fig. S2b**).

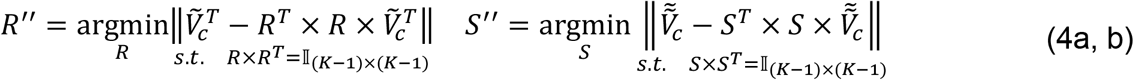

Note that the solution vectors for (Eq. 4 a, b) taken together with centering vectors 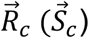 form two sets of vectors 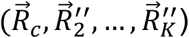 and 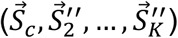 that can describe the hyperplane in the space of initial features and samples. However, these two sets of vectors are not necessarily mutually orthogonal. For instance 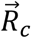 is typically not orthogonal to the hyperspace formed by 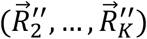 (**Fig. S2b**). Therefore, operators formed by 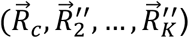 and 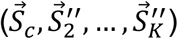 do not form projection operators acting on 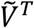 and 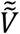 respectively.

Appropriate projection vectors can be obtained through an orthogonalization procedure applied to 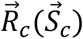, or equivalently, by introducing the orthogonalization constraint into the optimization problem (Eq. 5 a, b, Supplementary Note 6, 7):

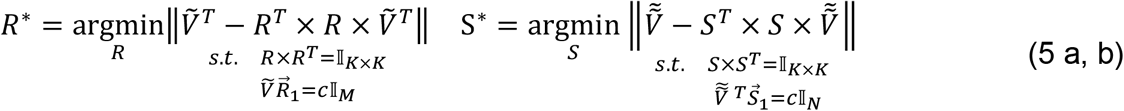

or in case of Sinkhorn transformed matrices (Eq. 6a, b, Supplementary Note 8):

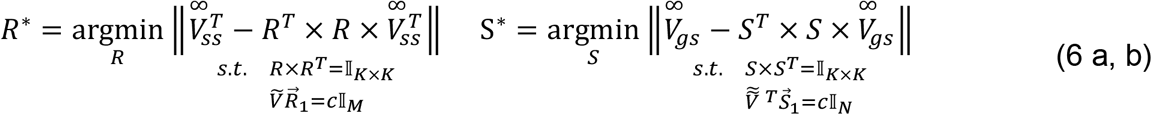

In the context of this optimization problem (Eq. 6 a, b), the special properties of Sinkhorn transformed matrices enable a very elegant identification of the hyperplanes in which simplexes are confined. Specifically, the singular value decomposition of the matrix 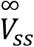 (or 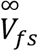) yields the sought-after projection vectors *R* and *S* for the constrained projection optimization problem (**Fig. 2d**, Supplementary Note 9 for proof). This represents the key theoretical result of our work, which is formulated as **the Dual Simplex Theorem**:

**Theorem**. *For Sinkhorn transformed matrices* 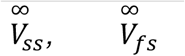 *(where* 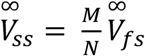)*obtained from the non-negative matrix V, solution vectors to optimization problems (eq. 6 a, b) can be found as follows:*

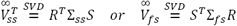

- *R*^*T*^ *could be found as left singular matrix of* 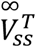 *and right singular vectors of* 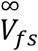
- *S*^*T*^ *could be found as right singular matrix of* 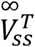 *and left singular vectors of* 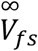

The simple consequence of this Theorem and (Eq. 2) is that singular values are linearly connected for matrices 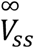 and 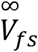, so that 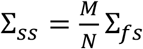. In short, the Sinkhorn transformation and subsequent SVD representation offer a clear path to identifying each of the individual hyperplanes where simplexes are located.

### Geometrical implications of the main theorem

Furthermore, from the main theorem and the matrix row/column normalization property, it follows that the pair of the top singular vectors 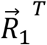 and 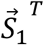 has a well-defined form and singular values. Specifically, vector 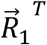 is 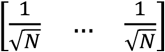 and corresponds to the singular value 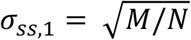 (**Fig. 2e**, see Supplementary Note 9 for proof). We demonstrate that the (*K* − 1) -dimensional simplex of samples space lies in the hyperplane that is strictly perpendicular to the vector 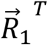, i.e. diagonal in the *N*-dimensional space, and intersects this diagonal at a specific point 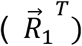. Similarly, the corresponding (*K* −1)-dimensional hyperplane of features space simplex is strictly perpendicular to the *M*-dimensional diagonal 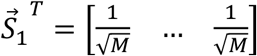, which is a singular vector corresponding to the singular value 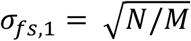. These vectors (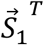 and 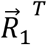 ) are present in the singular value decomposition for any Sinkhorn-transformed data matrix and are a consequence of sum-to-one scaling iterations. Geometrically, these vectors represent the shift of the simplex hyperplane away from the center of coordinates along the diagonal (Supplementary Note 9 for proof). The remaining (*K* −1)vectors of *R* (and *S*) define the subspace of the (*K* − 1) -dimensional simplex itself. For instance, in the case of 3 cell types (*K* = 3), a 2-dimensional simplex (a triangle) is located within a subspace defined by vectors 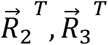 in the samples’ space and vectors 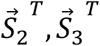 in the features’ space (**Fig. 2e**).

Understanding the structure of *R* and *S* also allows interpretable low-dimensional representations of the coefficient (*H*) and pure components (*W*) matrices as corresponding simlpexes. This can be done by introducing *K* × *K* simplex matrices 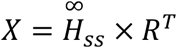 and 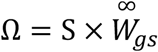 (**Fig. 3a**). Generally speaking, the rows of *X* and the columns of *Ω* define *K* points in *K*-dimensional space, which are located at the simplex vertices (**Fig. 3a**). In fact, as a consequence of the sum-to-one scaling operations and the corresponding form of the 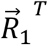 and 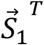 vectors, matrices *X* and *Ω* will also be partially known. For example, for 3 cell types, out of 9 elements (more generally, *K* × *K*), 3 elements of the matrix *X* (*Ω*)will be fixed (more generally, *K* elements) (**Fig. 3a**, Supplementary Note 9 for general case proof). The first column of matrix *X* will be a column-vector of 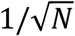 and first row of matrix *Ω* will be row-vector of 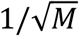 (**Fig. 3a**, Supplementary Note 9). Since rows of matrix *X* represent coordinates of the features’ simplex in *K*-dimensional space, the remaining *K* − *1* components of each row-vector of *X* will define simplex location in the (*K* − *1*) - dimensional space (**Fig. 3a**). Similarly, column-vectors of *Ω* define (*K* −1)-dimensional coordinates of simplex in the samples’ space (**Fig. 3a**).

**Figure 3.**
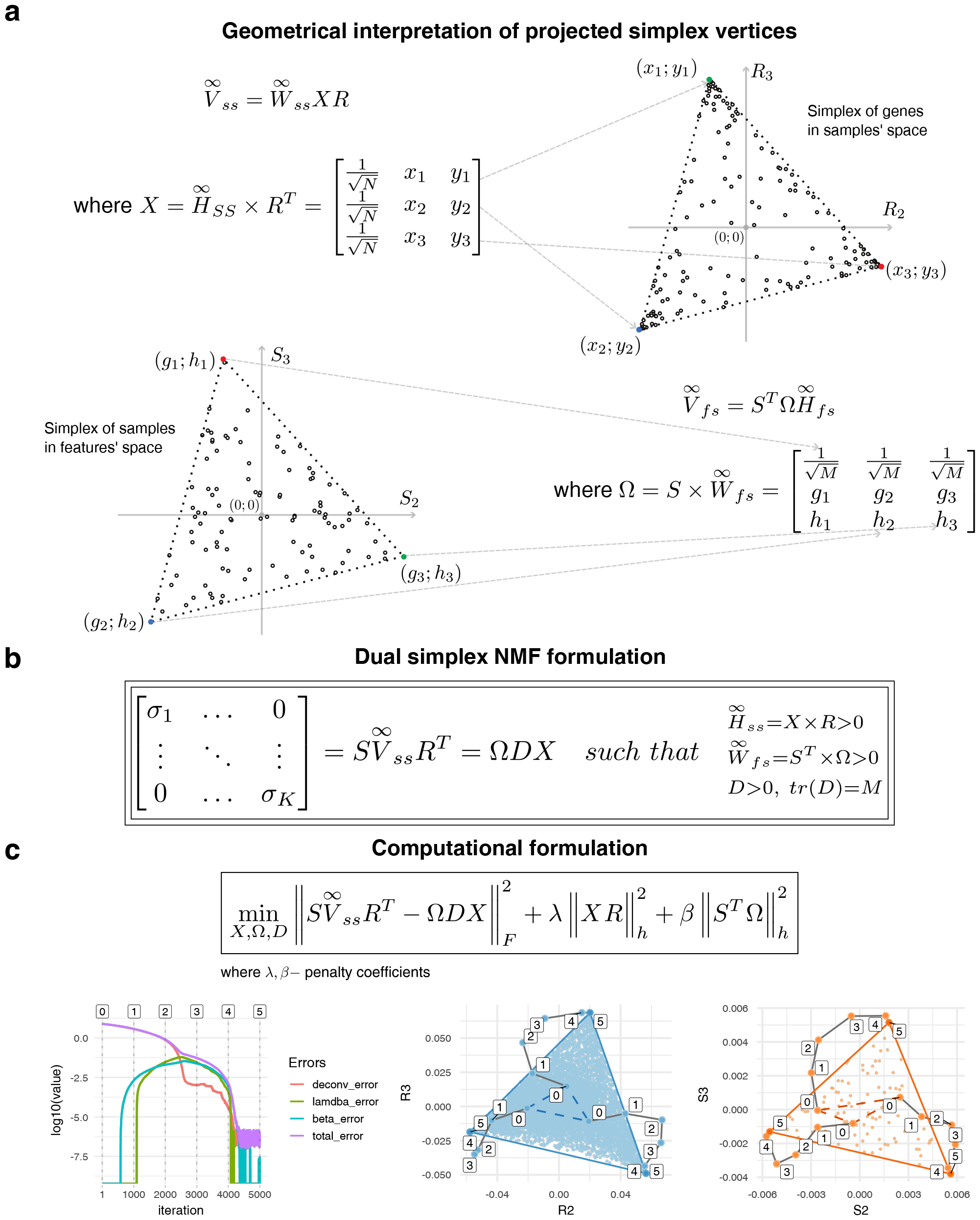
Interpretation of the simplex coordinates. Unified Dual Simplex problem formulation. (a) Geometrical interpretation of the projected data points: due to proportional relation between simplexes and constraints enforced for vectors R and S, the first coordinate of each projected point represents only shift of the hyperplane away from the center of coordinates, while the entire simplex structure is described by only (K-1) remaining coordinates. Search for Simplex vertices is therefore a search of a coordinates for geometrical simplex in a (K-1) -dimensional space (e.g., triangle in case when K=3). (b) Dual Simplex problem formulation is K-dimensional matrix equation equivalent to initial NMF problem. Left part of this equation is a precomputed singular values matrix while the right part contains three unknown variables representing coordinates of simplexes in both features and samples spaces as well as diagonal normalization matrix D which connects these spaces. (c) Gradient descent optimization formulation of the Dual Simplex problem. The cost function comprises the main deconvolution term as well as two additional terms (with coefficients A and ?) to address the positivity constraints. The optimization trajectory and endpoints in features and samples spaces for simulated gene expression data is presented on the bottom.

One can view deconvolution problem as search for simplex *X* in the samples space or search for simplex *Ω* in features space, which would be defined by two different optimization formulations (Supplementary Notes 4-7). However, both simplexes exist simultaneously within the Sinkhorn-transformed data. Below, we show that special properties of Sinkhorn transformation can be leveraged to explicitly reveal the relationship between the simplexes *X* and *Ω*.

### Unified Features’ and Samples’ dual simplex optimization problem

Given that left and right singular vectors are not randomly ordered but correspond to each other’s variability components, *the dual simplex theorem* indicates that there is a relationship between hyperspaces of features and samples simplexes. In fact, the Sinkhorn transformation enables a unified optimization problem formulation for two simplexes simultaneously.

Since matrices 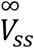 and 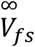 are proportional, we can merge two projection formulations into a single one (Eq. 7):

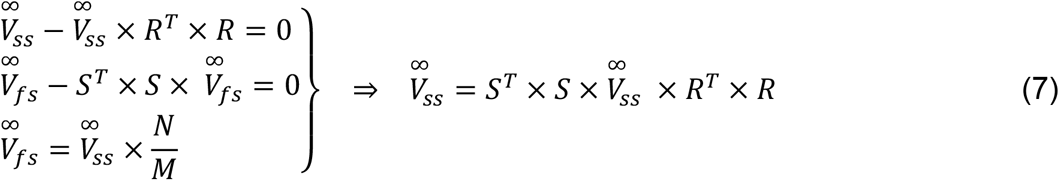

This then could be represented in terms of factor matrices 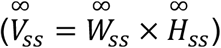 and their projection coordinates (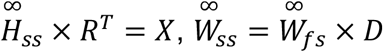 and 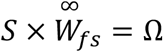), where *D* is a diagonal non-negative matrix that connects matrices 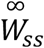 and 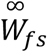 with property that *tr*(*D*)*=M* (Supplementary Note 8):

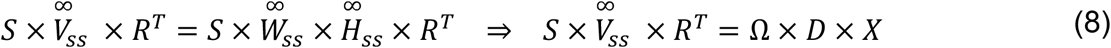

Equation 8 holds true for all exactly factorizable non-negative matrices and can serve as the basis for optimization problems in noisy and approximation scenarios. In such case, the main goal of the optimization is to minimize the difference between left and right part of this equation, thereby obtaining the approximate simplex coordinates (*X, Ω*) with highest possible precision. As a result, we describe the following common optimization formulation for the simultaneous search of samples’ and features’ simplexes (Eq. 9, **Fig. 3b**, Supplementary Note 8):

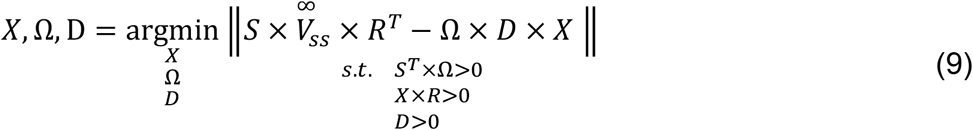

Note, that 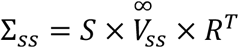 is a *K* × *K* diagonal matrix of singular values which is precomputed based on the Sinkhorn transformed expression matrix (see Main Theorem 1). Therefore, the deconvolution optimization problem (Eq. 9) becomes a problem of simultaneous optimization of two simplexes (*Ω* and *X*) in a low (*K* − 1) -dimensional subspaces, along with diagonal matrix *D*, which represents linear scaling between two simplexes (**Fig 3b**).

This problem can be tackled using a gradient descent numerical approach (**Fig. 3c**, Methods, Supplementary Note 10). Using gene expression simulated data with added noise as a simple illustrative study case ((Zaitsev et al., 2019), Methods), we find our computation procedure stably converging, producing valid results for the location of simplex corners (**Fig. 3c**).

### Geometrical interpretation of the relation between simplexes

In fact, the optimization problem described by (**Fig. 3**, Eq. 8) can be generalized to eliminate the explicit search for matrix *D*, and instead, only search for simplex corners of two modified simplexes 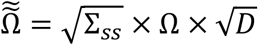 and 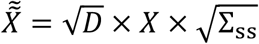, with only constraints on positivity and proportionality of 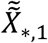 and 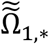 (Eq. 10, **Fig. 4a**, Supplementary Note 10).

**Figure 4.**
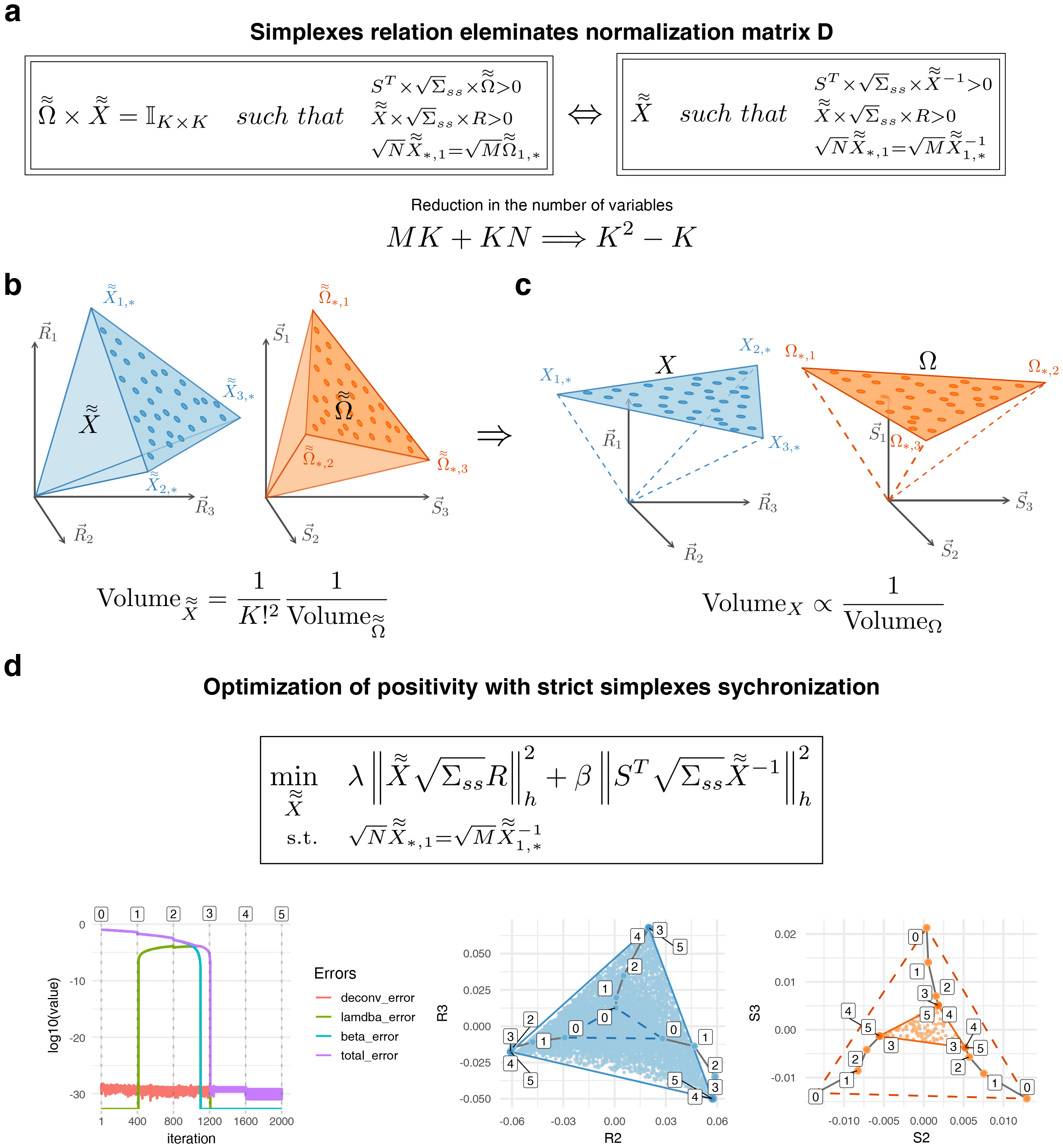
Minimal formulations of the Dual Simplex problem. (a) Dual simplex problem is equivalent to problem of finding single simplex with constraint on its inverse. This formulation demonstrates the dramatic reduction in the number of optimized variables achieved by Dual Simplex approach. (c) Geometrical interpretation of simplexes in features and samples spaces highlights inverse proportionality of their volumes. (d) Gradient descent computational formulation of the minimal formulation of the Dual Simplex problem. Here, only positivity terms (with coefficients 11 and ?) are optimized as deconvolution term is strictly enforced by the introduction of inverse. Below the optimization result for simulated gene expression data is presented.

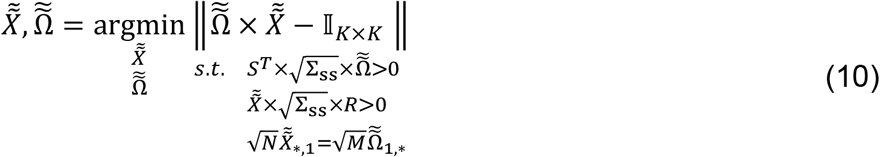

Strictly speaking, this means that we are only searching for a single simplex (either 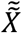 or 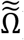) and imposing specific positivity constraints on it and its inverse (**Fig. 4a**):

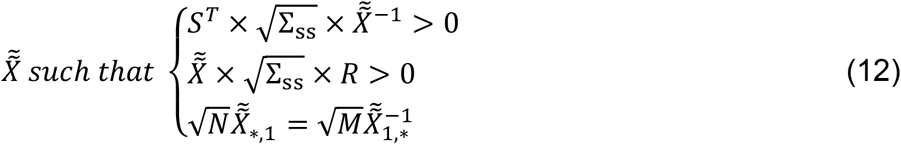

This approach allows for a significant reduction in the search space. For example, if you have 500 samples (*N*) and 10,000 features (*M*), but you are dealing with only 3 main components that make up all mixtures, the overall complexity of the deconvolution optimization task can be significantly reduced. Instead of searching for 31,500 values (30000 elements of 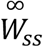 and 1500 elements of 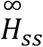), you would only need to search for 6 values, which correspond to the unknown elements of *X* (in general case *K*(*K* − *1*) variables). This substantial reduction in complexity is a notable advantage of the proposed approach.

Furthermore, this formulation reveals the geometrical meaning of the relation between simplex structures. The fact that the matrix of simplex corners 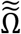 is the inverse of the matrix 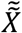 implies that the volumes of the corresponding hypercones defined by each simplex are inversely proportional (**Fig. 4b**, Supplementary Note 10). Moreover, this naturally propagates to a proportionality between the volumes of the original simplex coordinates *X* and *Ω* (**Fig. 4c**, Supplementary Note 10). This suggests that the problem of deconvolution of a positive matrix can be reduced to a problem of finding two related minimal volume simplex structures in both samples and features spaces. We further illustrate the convergence of this formulation as well as geometrical relation between solution simplex volumes (areas in *2D* case) using a gradient descent procedure applied to simulation data (**Fig. 4d**).

Once appropriate solutions for the corresponding optimization problems (**Figs. 3c, 4d**) in Sinkhorn transformed space are found, one can get original pure components feature profiles (*W*)and coefficient/proportions matrices (*H*)by applying reverse Sinkhorn scaling procedure to matrices 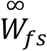 and 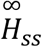 (Supplementary Note 11 for details).

Taken together, proposed framework (**Figs. 1-4**) describes a projective dual simplex formulation of the general non-negative matrix factorization problem. As such, it could be applied to any non-negative matrix to perform its non-negative factorization, or to special case of decomposition of the linearly mixed signals. Next, we will provide several examples of the practical applicability of this approach and discuss some special case implications of this approach for problems such as image decomposition, the single-cell RNA-seq data analysis, and deconvolution of gene expression data.

### Application of Dual Simplex approach to the NMF problem

To illustrate the applicability of our approach, we first applied our method to factorize the dataset, containing computationally mixed images (here, *K* = *4*) with either no noise or random proportional noise added (**Fig. 5a**). Note that the images were mixed with random positive coefficients *α*_*1*_, *…, α*_*4*_ that were not required to sum to one. The resulting mixed matrices were Sinkhorn-transformed, and the corresponding projections were visualized (**Fig. 5b, c**). The green lines on Figs 5b and 5c represent the true simplex coordinates for either the pictures’ or pixels’ space. It can be observed that in our image deconvolution scenario, even though the number of features (pixels) is high (*M*= *16*3*84*), pixel intensities in general are not specific to a particular picture, as all pixel points are relatively far away from the true corners of a simplex structure (**Fig. 5b, left**). In case when number of mixed samples is significantly smaller than number of pixels (*N* = *100 << M* = *16*3*84*), the simplex of mixed images in the space of pixels will have only 100 points, which is also relatively far from the simplex corners (**Fig. 5b, right**). Furthermore, the introduction of noise significantly distorts the structure of the data, making it even less related to the original simplexes for both pictures and pixels spaces (**Fig. 5c**).

**Fig 5.**
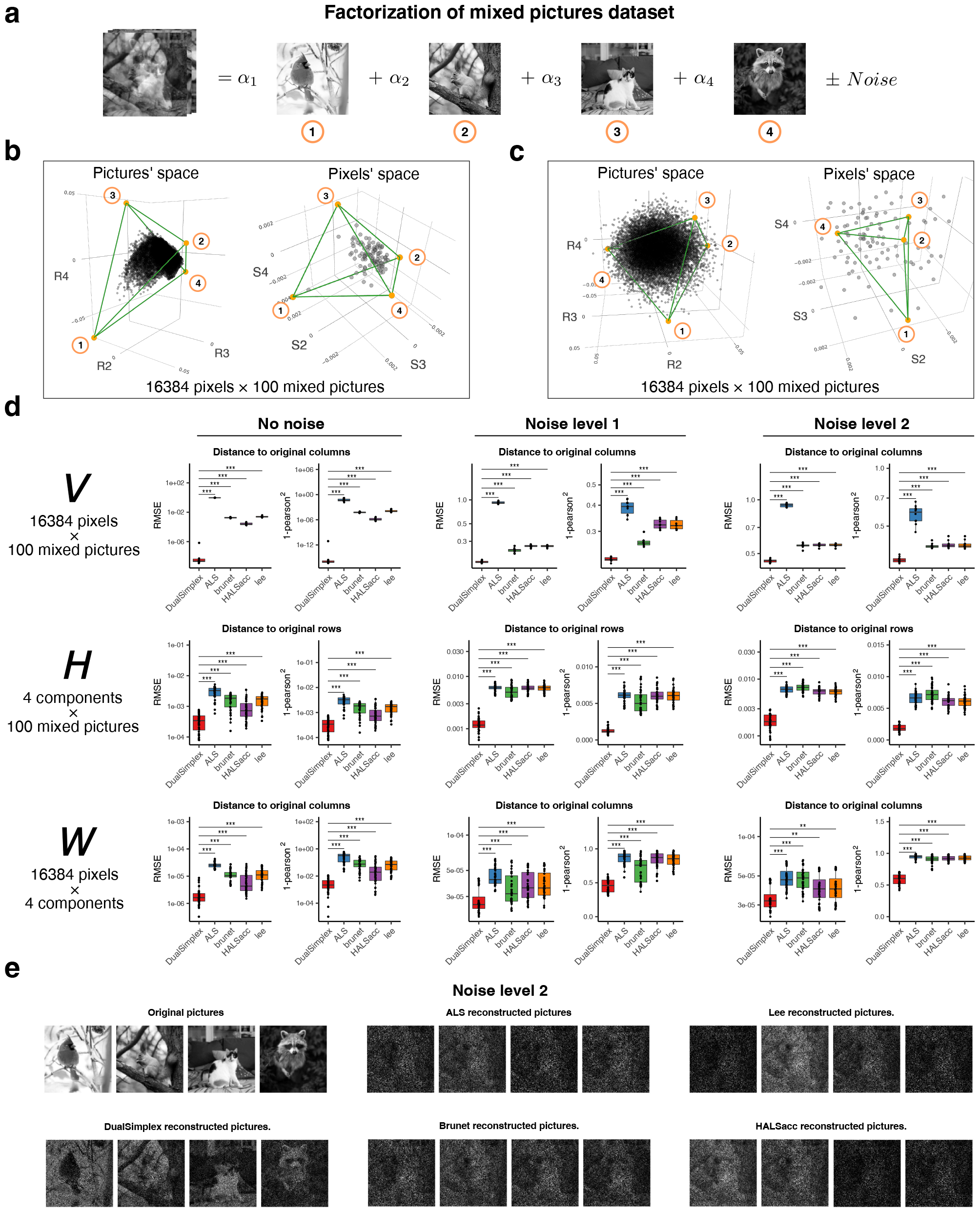
Application of the dual simplex approach to image data decomposition. (a) Each mixed image is obtained as a sum of the K=4 main picture components with nonnegative non-constrained coefficients. Noisy scenario involves introduction of the proportional noise with predefined standard deviation representing the noise level. (b) Visualization of the individual pixel points (left) and picture points (right) in respective projection space. Green lines represent true underlying simplex coordinates obtained by projecting rows of H and columns of W to respective space. For no noise scenario all the points are located within the geometrical simplex.(c) In case of noise present (noise_level = 2 in this case) the simplex structure becomes distorted and individual points can be located outside the original geometrical structure. (d) Reconstruction quality metrics measured for matrices V (row 1), H (row 2) and W (row 3) for different noise levels. Quality metrics are defined as median RMSE and 1-pearson^2 values measured between rows of predicted matrix H and true matrix H as well as columns of predicted matrices W and V in comparison to columns of the true matrices W and V. (e) Reconstructed images (columns of matrix W) visualized for all five methods for the high level of the noise (noise_level = 2).

In direct comparison of the accuracy of *V, H* and *W* reconstruction, our method (DualSimplex) outperforms commonly used NMF algorithms (Brunet et al., 2004; DeBruine et al., 2021; Gillis & Glineur, 2012; D. D. Lee & Seung, 2000), yielding significantly more accurate matrices *W* and *H* for all noise levels, both in terms of RMSE error and Pearson correlation coefficient of rows/columns (**Fig. 5d**). We observe that since the number of features, describing the simplex in a pictures space is high (*M*= *16*3*84*), all NMF methods can reconstruct *H* with high accuracy for all scenarios. However, the most striking difference in performance becomes apparent for the reconstruction of the *W* matrix of the pure components, for which the utilization of the complementary simplex structure helps significantly. The major improvement delivered by the Dual Simplex approach is apparent from the comparisons based on both RMSE or correlation-based metrics, as well as directly by looking at the reconstructed images (**Fig. 5e**). Therefore, we conclude that the Dual Simplex approach is more robust to noise values and allows for partial reconstruction of the original pure picture components even in situations where other methods fail to do so. Additionally, we tested it with an increased number of mixed pictures in a dataset, where we observed the same pattern (**Fig. S3a, b**).

Furthermore, to validate our approach in a most general and unconstrained matrix factorization task, we considered completely random positive matrices *W* and *H* (*M*= *1000, N* = *800*). For this scenario the Dual Simplex approach again demonstrated improved accuracy of reconstruction for both clean and noisy data matrices for both *W* and *H* (**Fig. S3c**). We also observed the same pattern that matrix W is reconstructed significantly better by DualSimplex approach compared to other methods, especially in the presence of noise.

### Single cell clustering analysis

One of the main areas where factorization of the multidimensional datasets may be required is gene expression data analysis, especially in the context of the single cell RNA-seq data. From the prospective of the linear mixture problem, single-cell RNA-seq data represent the data on thousands of samples where each sample is typically a pure cell type (with the exclusion of doublets and other artifacts). Conceptually, non-negative factorization of such dataset should yield clustering of the underlying data (DeBruine et al., 2021). This task in essence can be regarded as a deconvolution problem with only pure cell types/components present in experiment.

To demonstrate the ability of our method to solve this task we applied the method to a GSE103322 single cell dataset of head and neck squamous cell carcinoma (sc-HNSCC), containing 5902 cells extracted from 18 cancer patients. This dataset contains 2215 malignant as well as 3363 non-malignant cells of primary oral cavity tumors as well as metastatic lymph nodes. **Fig. 6a** reproduces t-SNE plot from original paper (Puram et al., 2017) with annotation provided by authors.

**Figure 6.**
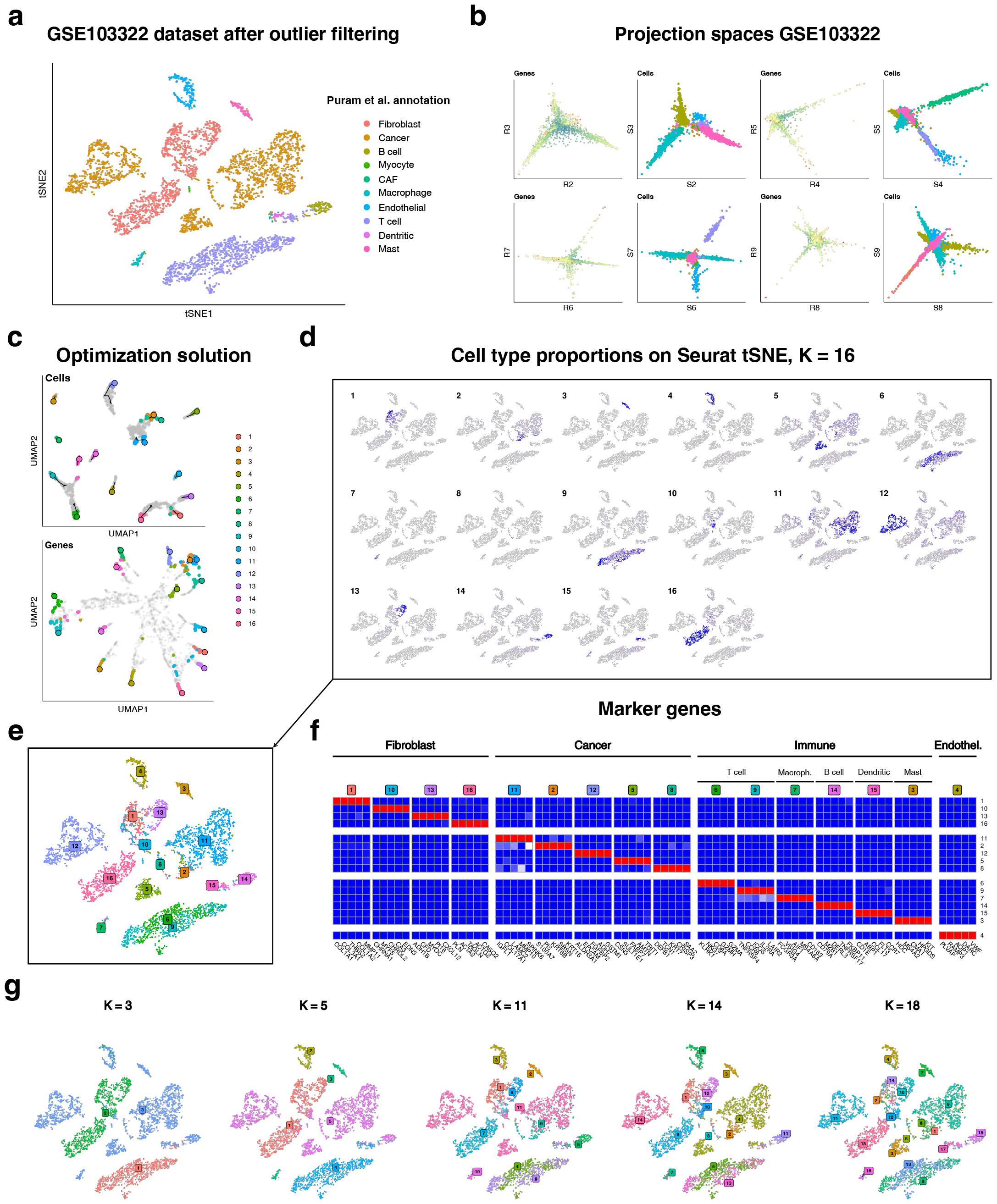
Single cell clustering with dual simplex approach. (a) A tSNE plot of the single cell HNSC (GSE103322) dataset with main clusters annotated by Puram et al. (b) First six dimensions for both genes (i.e. features) and cells (i.e. samples) projection subspaces (K=16) upon Sinkhorn transformation applied to filtered expression matrix. Cell populations are clearly separable within this subspace. (c) Dual simplex optimization procedure result with result simplex locations, optimization history and marker genes for the simplex genes. (d) Original tSNE cell points in a space of genes colored by proportion values (matrix H) for each of the K=16 main components. (e) Cell cluster labels assigned as component name with maximum proportion value for each individual point. (f) The heatmap of expression profiles for subsets of marker genes specific to main components. (g) Clustering result with different number of main components (K) selected.

For the analysis, we kept 3000 Most variable genes following the standard routine within the Seurat package (Hao et al., 2020), filtered out outliers and then Sinkhorn transformed resulting matrix (Methods). The SVD vectors of Sinkhorn transformed single cell matrix can be used to visualize different projections of the genes’ 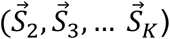 and cells’ 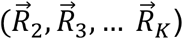 spaces (**Fig. 6b**). Cell subgroups/types are readily visually separable in these projections indicating that this representation can be used for clustering of different cell types.

We found it to be practical to utilize UMAP transformation (McInnes et al., 2018) of the projected points for visualization as it tends preserve cell-type-specific genes (those located far from the initial zero point) in the corners of the figure (**Fig. 6c**). Upon optimization procedure we observe solution points converged to corners of the genes simplex also represent different parts of island of the cells’ simplex (**Fig. 6c**). Moreover, resulting proportions matrix *H* provides us cell cluster locations, which can be visualized using original t-SNE plot of the data (**Fig. 6d, e**). At the same time columns of *W* represent transcriptional profiles of the main components. We show that these transcriptional profiles correspond to main subsets of cells reported within this dataset (**Fig. 6f**).

The Dual Simplex approach provides intuitive procedure and direct control over the clustering process. Indeed, one of the aspects of the commonly used single cell clustering pipelines is the need to build the neighborhood graph to understand the spatial distribution of the points within the space of main variational components, followed by clustering based on the distance thresholds. Such an approach results in unpredictable number of clusters. Our approach on other hand performs simultaneous alignment and clustering of the cells and genes providing a natural way to perform clustering of the data with explicitly controllable number of clusters (*K*) (**Fig 6g**).

Practically speaking, this DualSimplex based clustering requires optimization process which can take time and might be sensitive to parameter selection in some cases. However, this can be circumvented using simplified approximate procedure rooted in the fact that simplex corners are well separated in the case when mixture consists of only pure samples (e.g. scRNA-seq data). Utilizing simplest simplex corner finding algorithms such as VCA (Nascimento & Dias, 2005) provides set of points which are very close to the actual solution (**Fig. S4**). Subsequently, applying the K-means assignment procedure using these corner points as the cluster centroids provides a shortcut to obtain reasonably good clusters without any optimization procedure (**Fig. S4**). The results of such robust clustering are comparable to the results of the fully optimized DualSimplex approach as well as state-of-the-art single-cell clustering pipeline (Hao et al., 2020) (**Fig. S4**).

### Bulk gene expression deconvolution

#### Complete deconvolution

To demonstrate how the proposed complete deconvolution approach dissects real-world datasets of mixed samples, we first applied it to known benchmarks GSE19830 and GSE11058 (Abbas et al., 2009; Shen-Orr et al., 2010). These datasets represent mixtures of 3 and 4 pure components, which yield underlying simplexes as triangle and tetrahedron-like structures (**Fig. 7a, b**). Search in the genes and samples spaces yield accurate corners of the corresponding simplexes which map to the genes specific to pure cell types and accurately recover proportions of pure components (**Fig. 7a, b**).

**Figure 7.**
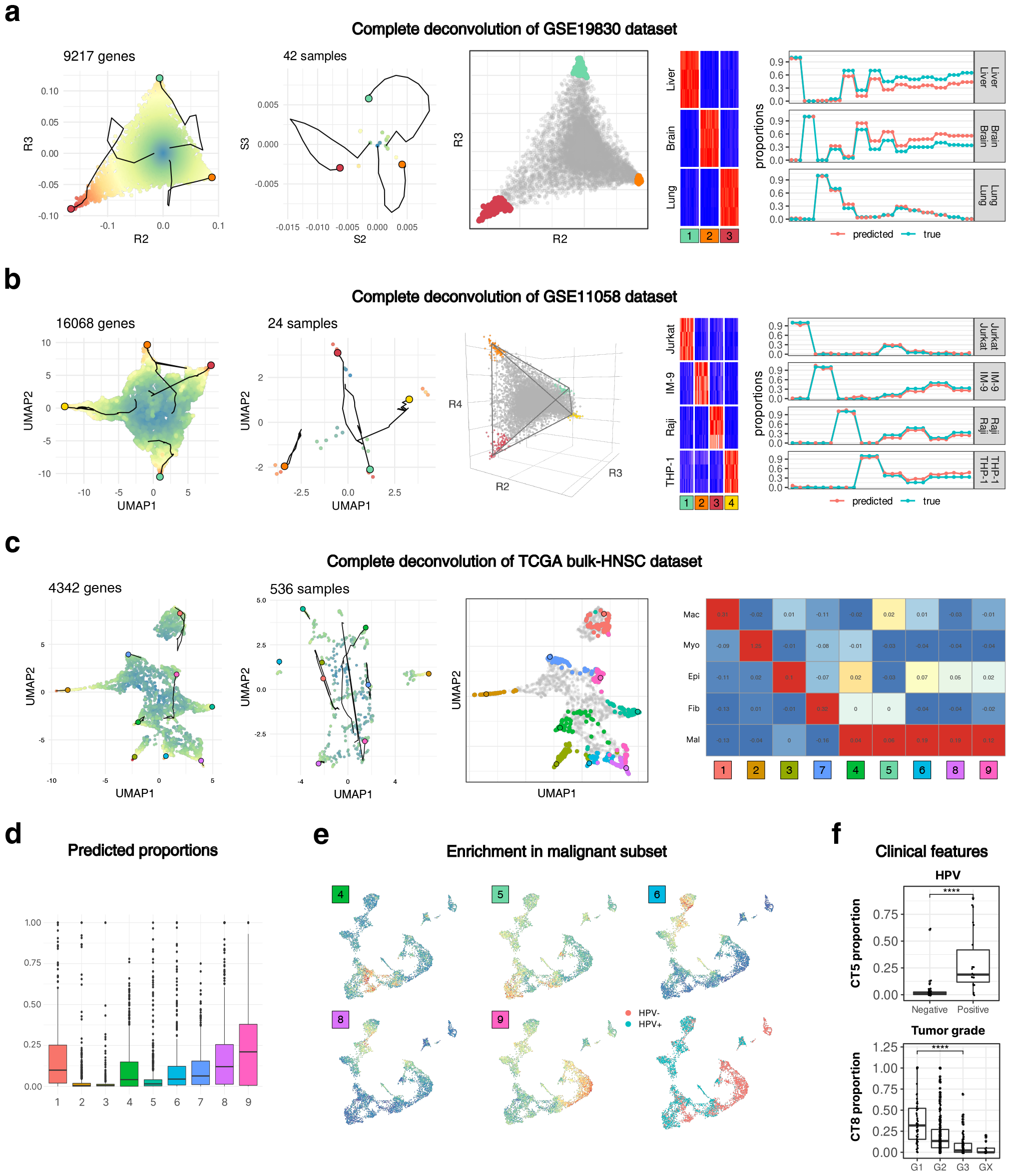
Complete deconvolution of bulk-RNA datasets with dual simplex approach. (a) Complete deconvolution of GSE19830 with result simplex location and optimization history for both genes and samples spaces (left), extracted marker genes highlighted on a simplex of genes (middle) and heatmap of gene expression from pure lung, brain and liver samples (heatmap). Predicted proportions values for each sample compared to true proportions extracted from the data (right). (b) Complete deconvolution of GSE11058 with result simplex location and optimization history for both genes and samples spaces (left), obtained marker genes highlighted on a simplex of genes (middle). Marker genes transcriptional profile compared to signature genes extracted from pure samples (heatmap). Predicted proportions values for each sample compared to true proportions extracted from the data (right). (c) Complete deconvolution of TCGA bulk-HNSC dataset with result simplex location and optimization history for both genes and samples spaces (left), obtained marker genes highlighted on a simplex of genes (middle). Enrichment of predicted cell types’ marker genes in a major annotation cell subsets extracted from the external GSE181919 dataset (right). (d) Predicted proportions distribution for each of the main components. (e) Enrichment of predicted malignant cell subsets on a UMAP of malignant cells. (f) Correlation of predicted proportions with selected clinical features.

For a more complex and realistic scenario, we applied the dual simplex approach to a TCGA bulk RNA-seq dataset profiling *M*= *548* samples of the Head and Neck Squamous Cell Carcinoma (bulk-HNSCC) cancers characterized by the global gene expression profiles. For the preprocessing of this type of data, we find it beneficial to keep only reasonably well-expressed genes and filtering out noisy and housekeeping genes, which introduce distortion to singular value decomposition (see Methods for details).

We found our computational approach stably converging for the *K* = *9* cell type specific points perfectly aligning with corners of a simplexes (**Fig. 7c, S5c**). To validate the results of the deconvolution we assessed the enrichment profile of the predicted marker genes in the GSE181919 single cell dataset. Our analysis shows that each predicted cell type is biologically relevant to a specific subset of the single cell data, revealing 4 major non-malignant cell types (including macrophages, myocytes, ciliated epithelial cells, and fibroblasts) as well as 5 different cancer subtypes (**Fig. 7c, d**). To understand if these cancer subtypes represent real subpopulations or mathematical artifacts, we zoomed into malignant cells and found that five identified cancer subtypes are enriched across different malignant cell subclusters (**Fig. 7e**). To further confirm the biological relevance of the predicted signatures, we measured the enrichment of 100 markers per predicted cell type in msigdb (Liberzon et al., 2011). The enriched pathways for each cell type generally correspond to single cell-based assumptions (**Fig. S5a, b**). Of interest, we found that cancer subtype 5 is significantly more abundant in the samples of HPV-positive patients and that cancer subtype 8 was inversely correlated with tumor grade (**Fig. 7f**). These experiments illustrate the effectiveness of our computational approach in solving the deconvolution problem for noisy simulated and high-dimensionality real-world datasets.

#### Signature-based partial deconvolution

Important scenario for the deconvolution problem is dissecting the mixtures when only key marker genes of the pure components are available (e.g., spatial transcriptomics, scRNA-seq based bulk deconvolution etc). Generally knowing marker genes for a cell type is equivalent to knowing coordinates of the simplex corners in a sample space (i.e., matrix *W*). The framework described in this paper provides straightforward procedure for partial deconvolution using Dual Simplex approach: the gene space simplex is defined by reduced markers expression matrix, leading to proportion matrix *H* (Methods). This reduces the problem to conventional partial deconvolution problem with known proportions and complete profiles of pure components, which can be defined via, for example, Non-Negative Least Squares (NNLS) procedure. We illustrate the signature-based partial deconvolution using GSE11058 data (Abbas et al., 2009) of 4 mixed cell types and single-cell derived computational mixing approach by Sutton et al (Sutton et al., 2022) (**Fig. S6**). As can be seen, implementation of the dual simplex approach behaves robustly and accurately recovers proportions and gene expression profiles of pure components (**Fig. S6**).

## Discussion

Our work builds on the previously described idea that the problem of linear and sum-to-one constrained matrix unmixing is in essence a problem of a simplex identification (Boardman, 1993). Various approaches for finding such simplexes were proposed for general hyperspectral images (Boardman, 1993; Nascimento & Dias, 2005; Winter et al., 1999). With introduction of the non-negative matrix factorization problem (D. D. Lee & Seung, 1999), the linear unmixing was recognized to be a sum-to-one constrained NMF problem (Tao et al., 2007). Furthermore, it was noted that general NMF problem can be transformed to linear unmixing/deconvolution problem via matrix normalization (Chan et al., 2008). In biological settings, we, and others, explored simplex-based methods in the context of the gene expression data deconvolution (Wang et al., 2016; Zaitsev et al., 2019). Importantly, the simplex structures examined thus far in these, and other applications were designed for a collection of points forming the feature vectors (which correspond to the rows of the original matrix, such as genes). The key feature of our approach stems from the observation that the input data simultaneously contain two simplexes: (1) simplex that is revealed upon row-normalization, i.e. simplex in the samples’ space, and (2) simplex that is revealed upon column-normalization of the matrix, i.e. simplex in the features’ space. Conceptually, this observation reflects the equivalence of the features and samples in the linear unmixing problem.

Recognizing these properties, we developed novel analytical framework for non-negative matrix factorization grounded in topological properties of the problem. Using projective formulation (Yuan & Oja, 2005) of the Sinkhorn-transformed NMF problem, we explicitly connect two simplexes. This approach yields unified dual simplex optimization formulation of the NMF/deconvolution problem (Eq. 8, **Fig. 3**). Our approach reveals that NMF problem can be effectively reduced to optimization of K(K-1) independent variables, K being number of cell types, clusters or independent components in general.

The Dual Simplex approach provides unique power in terms of dimensionality reduction and robustness relative to disbalance between number of features and samples, as optimization requires the best fit for both simplexes. Indeed, our results from unconstrained nonnegative factorization of image datasets demonstrate that simulations optimization for both simplexes makes our algorithm more robust to noise and provides significantly more accurate decomposition matrices for the general NMF problem compared to other methods, even though the optimization itself is conducted in space of much lower dimensions (**Fig. 5, S3**).

Moreover, our approach provides a unified framework for solving different flavors of the sum-to-one constrained version of the NMF problem, i.e. deconvolution problem, which is commonly utilized in a computational biology due to the wealth of the information from RNA-sequencing data. These flavors include complete and signature-based deconvolution of bulk-RNA sequencing data (**Fig. 7, Fig. S5, S6**), Furthermore, one can easily expand this approach to address *incomplete deconvolution* problem, i.e. the situation when some of the cell types in the mixture are known while others are not. In this problem, one can freeze the solution in a simplex corner for already known markers (e.g., using signature-based approach described), and then perform optimization, searching for rest of the corners in a low-dimensional space, utilizing the geometrical properties of the described simplex structure.

Besides that, we demonstrate that our method can efficiently dissect the cluster structure of the data, e.g. single cell RNA-seq data (**Fig. 6, S4**). From one perspective this problem can be regarded to as an extreme case of deconvolution with only pure components present in the data. Therefore, in the genes’ space, sample-dots are concentrated in the vertices of the simplex with almost no points in between. In these settings, deconvolution of the scRNA-seq data would be equivalent to clustering of the cells in the space of genes. On the other hand, one can think about obtained pure components in terms archetypes constituting the expression profile of a single point as described by Alon and colleagues for bulk and single-cell gene expression data (Korem et al., 2015). This idea was then developed using different computational and numerical approaches such as kernel analysis etc (Persad et al., 2023; van Dijk et al., 2018). Our work effectively provides rigorous mathematical basis and procedure for implementation of the archetypical (i.e. simplex) structures within the gene expression data.

More broadly, the ability to solve the NMF problem suggests the applicability of our approach to a wider range of tasks such as those that can be formulated as NMF problems, including spatial transcriptomics analysis (Rodriques et al., 2019), mass spectrometry analysis (Nijs et al., 2021), and spectral cytometry analysis (Jiménez-Sánchez et al., 2020) as well applications beyond computational biology.

## Acknowledgments

Authors would like to thank computational community at Washington University in St. Louis and ITMO University for useful suggestions and helpful comments, in particular Alexey Sergushichev and Konstantin Zaitsev. D.K. also would like thank Ekaterina Vysotckaia for moral support during manuscript preparation.

## Methods

### Data and Implementation

The entire framework is implemented using R language with a C++ backend and Armadillo library for linear algebra calculations. (Sanderson & Curtin, 2018). It is available online in the form of source code repository (https://github.com/artyomovlab/dualsimplex). Scripts to reproduce all the figures as well as corresponding data are stored in a separate repository (https://github.com/artyomovlab/dualsimplex_paper).

### Simulation data generation

For general matrix unmixing problems *V* = *W* ×*H*, matrices *H* and/or *W* were sampled from a random uniform distribution *W,H∼U*(*0, 10)* . The additive proportional noise was then applied using formula *V*_*noisy*_ = *V* * (*1 + N*(*0, σ) )*, where *σ* represents the chosen level of noise deviation. Negative elements of such matrix were set to zero.

For gene expression scenarios, when generating basis vectors (columns of the matrix ***W***), expression values *x* were chosen using formula *x* = *2*^*X*^ where *X* is a random variable sampled uniformly from two normal distributions, specifically 𝒩(*4, 0*.*75*)and 𝒩(*10, 1*.*5)*, with a *2/*3 ratio. This type of expression generation was designed with the assumption of a bimodal nature of the expression data. The first mode represents the expression of lowly or non-expressed genes, while the second mode represents well-expressed genes. Conversely, the proportions for a given cell type (rows of the matrix ***H***) were sampled as random points within a geometrical simplex structure to enforce the sum-to-one constraint. In the case of noisy data scenario, the noise values (*y*) were generated using formula *y* = *2*^*σY*^ where *Y* follows standard normal distribution (𝒩(*0*, 1) ) and *σ* represents the chosen level of noise deviation.

### Gene expression data preprocessing

To apply our method to a real expression dataset, we recommend a three-step procedure for data cleaning, which enhances the accuracy of identification for the hyperplanes of the simplexes (i.e., vectors *R* and *S*).

#### Step 1: Data prefiltering

We suggest prefiltering of highly correlated genes, such as ribosomal and mitochondrial genes. Additionally, it is beneficial to remove housekeeping genes since such genes are not related to any cell type specifically but can introduce bias during singular value decomposition. For improved biological interpretation we also remove non-coding genes, although it is not necessary from the algorithmic perspective. Furthermore, since an exact definition of housekeeping genes is lacking, we found it practical to not only remove genes derived from previously designated housekeeping genes lists but also to eliminate those with expression profiles that highly correlate with these genes. To achieve this, we employed a straightforward KNN-based approach (Ripley, 1996), considering the proximity between values of genes in the projected space as this proximity reflects the similarity in their expression patterns across samples. Complete lists of genes specific for each of the filtered subsets could be found in (Supplementary Table 1).

#### Step 2. Filtering based on variability

Since our approach leverages linear components of variability in gene expression to dissect cell type composition, it is beneficial to remove the genes that are lowly varying. In order to achieve this, we perform median absolute deviation (MAD) filtering, keeping genes that have MAD greater than selected threshold. The precise number of genes filtered was chosen based on each dataset specifically. The distribution of MAD in the data typically forms a distinctive bimodal distribution, where one group of genes contains lowly variated genes and technical artifacts, whereas the second group of genes contains highly variated genes. The threshold for the filtering was chosen for the first group to be filtered out.

#### Step 3: Data denoising/Outlier removal

Another way to improve the projection optimization solution is to denoise the data by removing outlier genes and samples, since SVD-based approaches are generally sensitive to outliers. Here we suggest using the approach of removing genes/samples, which are too far from the projection hyperplane in terms of geometrical distance. Details of approach are found in Supplementary Note 12.

Our approach implements an iterative filtering process where step 3 can be repeated, allowing for gradual strengthening of filtering conditions. The improvement is generally evaluated via 2-dimensional projection plots to major singular vectors, as well as plane and zero distance plots, with the goal of observing reasonably homogenous group of dots without distant outliers (see Supplementary Note 12 for details).

### Optimization procedure

Minimization problem (eq 8) was solved using gradient descent optimization procedure. The main implementation uses cost function (**Fig. 3c**) and is in essence a block-wise gradient descent procedure operating in a projection space with respect to 3 matrix variables (*X, Ω, D*)with the sequence of updates *X → D → Ω → D*. For the variables *X* and *Ω* algorithm computes partial derivatives ∇*F*_*X*_ and ∇*F*_*Ω*_ which then are used in update rules 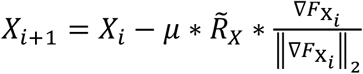 and 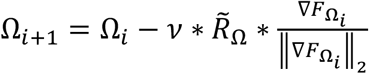. Here *µ* * 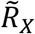 and *𝒱* * 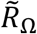 are scaling constants for step magnitude based on the average norm of projected vectors in their respective spaces. New values for the variable *D*, calculated analytically via NNLS:

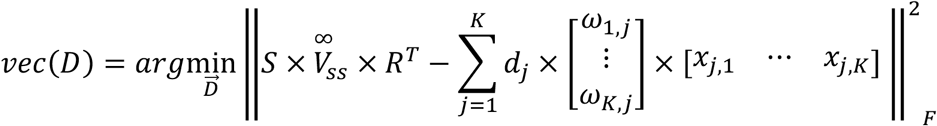

### Initialization strategies

To select initial values for matrices *X* and *Ω*, we employed three different strategies: *random centered, select_x* and *select_omega*. The first strategy involves selecting corner points that are centered around zero for a set of randomly generated coordinates.

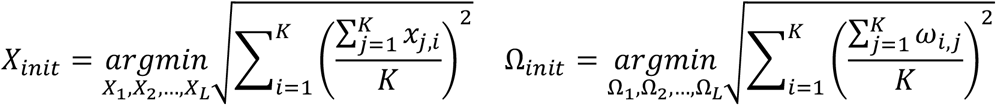

The matrix *D* for this scenario is obtained as a solution for non-negative least squares (NNLS) problem:

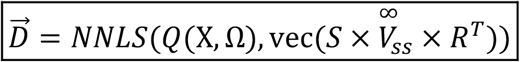

The select_x and select_omega initialization methods involve geometrical initialization using the VCA algrorithm (Nascimento & Dias, 2005) for one of the initial simplexes (e.g. *X*), followed by analytical imputation of another one (e.g. *Ω*). This scenario is especially effective in cases, when one of the simplexes is described better due differences in number of points (e.g. *N* ≫ *M*) or the quality of these points (e.g., for single cell data we have only pure component points for *Ω*, which already describe the simplex).

We start if geometrical initialization of first simplex:

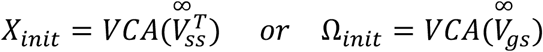

After this we get the 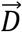 based on the known values using Sinkhorn transformed matrices properties, which we have proven (Supplementary Note 10).

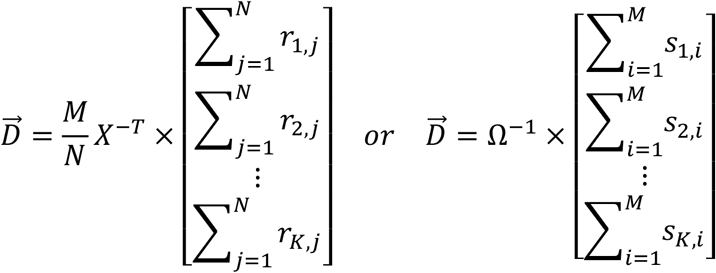

And finally obtain missing coordinates for second simplex.

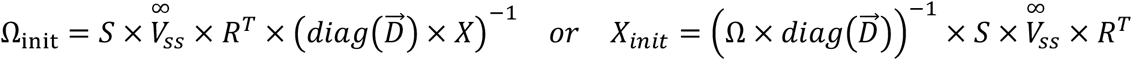

### Non-negative matrix factorization

For NMF problem statement, we consider *K* = *4* different flattened pictures of the same size (*M*= *16*3*84 pixels*), which are utilized as columns of the original components’ matrix *W* (*16*3*84 pixels* × *4 pictures)* . The input matrix *V* comprises the chosen number *N* mixed pictures (for example *N* = *100* for **Fig. 5**), where each column *i* was generated as a sum of main component pictures with different coefficients:

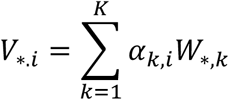

Here, coefficients *α*_*k,i*_ constitute the columns of the coefficients matrix *H*. These are sampled uniformly from the interval *[0,1]* and do not necessarily sum to one.

The dual simplex procedure was applied to the result matrix *V* without additional filtering steps. The optimization procedure was randomly initialized involved a sequence optimization steps with gradual decrease of the learning rate (12 iterations starting from (*µ* = *0*.*1, 𝒱* = *0*.*1*) and ending with (*µ* = *10*^−*12*^, *𝒱* = *10*^−*12*^)). During each optimization step negativity penalty coefficients were increased multiple times using the formulas *α* = *µ*^*2*^ * *100*^*i*^ and *α* = *𝒱*^*2*^ * *100*^*i*^ where *i* is a current iteration of optimization step (we performed 3 iterations per step). This design was chosen to achieve comparable values and gradual convergence for all cost function terms.

Additionally, for each scenario four different NMF algorithms Lee (D. D. Lee & Seung, 2000), Brunet (Brunet et al., 2004), HALSacc (Gillis & Glineur, 2012) and ALS (DeBruine et al., 2021), were tested using NMF R package (Gaujoux & Seoighe, 2010).

### Comparison of different NMF methods

Since nonnegative matrix factorization does not have any special constraints for *W* and *H*, these matrices could only be reconstructed up to the scaling factors of the columns of *W* and rows of *H*. This produces incomparable matrices *W* and *H* for different methods. To make them we scale columns of *W* and rows of *H*, representing all solutions in terms of column normalized matrix *W*_*colnorm*_, row normalized matrix *H*_*rownorm*_ and the scaling matrix *D*.

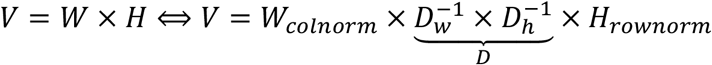

The accuracy of such unified NMF solutions were the estimated using two metrics the RMSE error (*rmse*) and transformed Pearson correlation between matrix values (*pearson)* . Normalized columns of the solution basis matrix *W′*_*colnorm*_ were compared to normalized columns of original matrix *W*_*colnorm*_, and vice versa, normalized rows of solution matrix *H*^*′*^ were compared to original normalized rows of the matrix *H*_*rownorm*_. For matrix *V* the same functions were measured for the columns of the matrix.

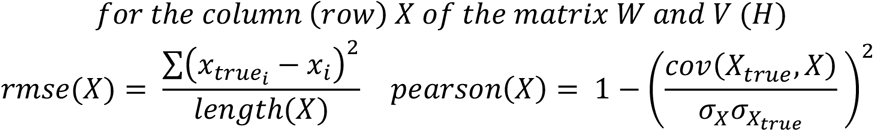

### Single cell RNAseq HNCS dataset clustering

The GSE103322 dataset of head and neck squamous cell carcinoma extracted from the GEO database (Clough & Barrett, 2016) and linearized (*V* = *2*^*V*^). Top 3000 most variable genes were selected using the Seurat FindVariableFeatures method (Hao et al., 2020). For outlier filtering, we applied consequent distance cutoffs, reaching thresholds of *zero_distance* = *0*.*05, plane_distance* = *0*.*15* for genes, and *zero_distance* = *0*.*16* for samples. This resulted in a dataset of 2517 genes and 5727 cells.

We then performed nonnegative factorization of this dataset via the Dual Simplex approach with the number of pure components *K* ranging from 3 to 21. The gradient descent was initialized using the select_x method, and 2 consecutive steps of optimization were performed: the first step consisted of 650 iterations with parameters (*λ* = *0*.*01, β* = *0*.*01, µ* = *0*.*001, 𝒱* = *0*.1),and the second step consisted of 3000 iterations with parameters (*λ* = *0*.*001, β* = *0*.*001, µ* = *0*.*001, 𝒱* = *0*.*00*1) ). Clusters were then assigned to cells as the row index for the maximum value of the column representing this cell in the matrix *H*.

### TCGA bulk-HNCS dataset deconvolution

The dataset, along with clinical data, was retrieved from GDC Data Portal, using the project_id of TCGA-HNSC.

The dataset was filtered according to proposed procedure. First, we applied a MAD cutoff by the threshold of *log_mad* = *0*.*1*. After that, we calculated an initial projection of the data points onto simplex hyperplanes (*K* = *20*) and removed HK genes with their closest neighbors in genes hyperplane, using the threshold of *KNNd* = *0*.*015*. For outlier filtering, we employed consequent distance cutoffs, reaching thresholds of *zero_distance* = *0*.*2, plane_distance* = *0*.*15* for genes and *zero_distance* = *0*.*1, plane_distance* = *0*.*045* for samples (See Supplementary Table 2 for more details). This resulted in a dataset with *M*= *4*3*26* genes and *N* = *5*3*5* samples. The information about removed and retained genes is available in Supplementary table 1.

Housekeeping and non-coding gene annotations were retrieved from publicly available sources (Eisenberg & Levanon, 2013; Hounkpe et al., 2021) and Ensembl (Martin et al., 2023) respectively.

After this initial filtering, the elbow point in SVD plot began to manifest, establishing a range of approximately 7 to 12 potential cell types for analysis. For cell type numbers within this range, hyperplane projections were calculated. To determine the final number of cell types, the deconvolution procedure was executed across this range of 7–12 potential cell types. Several criteria were used to identify the optimal number of cell types: the alignment between the number of corners on the genes UMAP and the number of points in the solution; the distinctiveness and specificity of the predicted cell type signatures; the biological relevance of the identified cell types. Using these criteria, it was established (subjectively) that the most suitable number of cell types for demonstration purposes is 9. It is noteworthy that the deconvolution can be extended to accommodate a larger number of cell types, unveiling smaller and less distinct cell subpopulations. However, such subpopulations, although genuine, were not as suitable for demonstration with single cell reference.

After that, a deconvolution, using optimization, was performed. The gradient descent was initialized using the select_x method, and run for 9000 steps, with *hinge_H* = *0*.*01, hinge_W* = *0*.*01*, and a learning rate of 0.001.

### TCGA bulk-HNCS deconvolution results analysis

Marker genes enrichment in single cell data was calculated by taking *n* = *100* top marker genes per cell type and calculating module scores for them using the AddModuleScore function from Seurat 4.0.0 (Hao et al., 2020). Aggregated heatmaps show the average expression across all cells within each analyzed single cell cluster.

Pathway analysis was performed by taking the same marker genes and calculating p-values for their enrichment in reference gene sets with clusterProfiler 3.18.1 (Yu et al., 2012). MSigDB gene sets from the H, C2:CGP, GO:BP and GO:CC categories were retrieved using msigdbr 7.2.1 (Dolgalev, 2022) and used as reference for clusterProfiler.

For clinical associations analysis (Fig. 6G), sample annotation was obtained from GDC (Grossman et al., 2016). Potentially relevant features were selected manually, numerical features were converted to binned categorical values. Subsequently, Kruskal-Wallis tests were conducted for each of the selected features, considering each cell type. The objective of the experiment was to assess whether samples, separated based on specific feature values, demonstrate statistically significant differences in predicted cell type proportions.

### GSE181919 single cell data preprocessing

The whole dataset without additional filtering as well as the malignant subset, were separately scaled and normalized using standard Seurat v. 4.0.0 procedure, integrated using harmony 1.2.0 (Korsunsky et al., 2019) with parameter *N_PC* = *20*, and mapped onto UMAP coordinates, in which form they are presented in Fig. 6.

### Signature-based partial deconvolution

The algorithm for signature-based deconvolution (**Fig. S6**) accepts the same gene expression matrix *V* as the complete algorithm. It also requires a list of marker gene names for each of the cell types present in the mixture.

The expression matrix and signature genes for a deconvolution benchmark GSE11058 dataset were obtained using CellMix package (Gaujoux & Seoighe, 2013). The complete expression matrix comprises 24 samples of 4 mixed cell types with expression values for 7862 genes. For each cell type, 20 marker genes were selected as input for the algorithm resulting in a reduced expression matrix *V*_*mark*_ (*80 genes* × *24 samples*). Solution for each cell type was set as a mean location for the respective marker genes in the projected space. This initialization produced a signature proportion matrix *H*_*mark*_ (*4 cell types* × *24 samples*). This matrix was then used to infer the basis matrix *W* (*7862 genes* × *4 cell types*)through conventional NNLS procedure using the original matrix *V*:

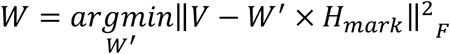

The same procedure was applied to deconvolve a single-cell-based simulation dataset (Sutton et al., 2022) consisting of *K* = *7* cell types mixed in *N* = *100* samples. For this data, three different marker sets were extracted from (Sutton et al., 2022), specifically VL (Velmeshev et al., 2019), NG (Nagy et al., 2020),CA (Hodge et al., 2019)

This approach can be particularly useful when applied to the datasets where the simplex structure is distorted due to technical or biological noise, making precise detection of the simplex plane through SVD and simplex corners through optimization a challenging task. In these situations, the removal of a significant portion of the genes effectively eliminates unwanted variations associated with those genes, while still retaining the essential corners of the gene simplex.

## Figure legends

**Fig. 1. Existence of complementary features and samples simplex structures for normalized matrices**.

a. Schematic representation of the data matrix in features and samples spaces

b. Row normalization and transposition of original matrix aligns feature dots within a simplex in a samples space

c. Column normalization aligns sample dots within a simplex in features space

**Fig. 2. Sinkhorn transformation enables identification of the simplex hyperplane projection vectors**.

a. Sinkhorn transformation is a process of iterative left and right multiplication by diagonal matrices (i.e. row and column normalization), producing two converging sequences of matrices, representing simplexes in features and samples spaces.

b. Converged matrices in features and samples spaces are proportionally related.

c. Example of Sinkhorn transformation applied to randomly generated factorizable matrix (*K* = 3,*M* = *1000, N* = *1000*). While original spaces of features and samples does not contain simplex structures (left), column and row normalizations reveal simplex structure for one of the spaces (middle). Iterative normalizations applied to the matrix align points withing simplexes for both spaces (right).

d. The Dual Simplex theorem formulation. Singular vectors of Sinkhorn-transformed matrices provide projection vectors to hyperplanes in which samples and features simplexes are located.

e. Corollary of the Dual Simplex theorem: due to normalization constraint *R*_*1*_ and *S*_*1*_ have fixed form and represent the shift of the entire simplex hyperplane from the zero point.

**Fig. 3. Interpretation of the simplex coordinates. Unified Dual Simplex problem formulation**.

a. Geometrical interpretation of the projected data points: due to proportional relation between simplexes and constraints enforced for vectors *R* and *S*, the first coordinate of each projected point represents only shift of the hyperplane away from the center of coordinates, while the entire simplex structure is described by only (*K* −1)remaining coordinates. Search for Simplex vertices is therefore a search of a coordinates for geometrical simplex in a (*K* −1)-dimensional space (e.g., triangle in case when *K* = 3).

b. Dual Simplex problem formulation is K-dimensional matrix equation equivalent to initial NMF problem. Left part of this equation is a precomputed singular values matrix while the

c. right part contains three unknown variables representing coordinates of simplexes in both features and samples spaces as well as diagonal normalization matrix D which connects these spaces.

d. Gradient descent optimization formulation of the Dual Simplex problem. The cost function comprises the main deconvolution term as well as two additional terms (with coefficients λ and β) to address the positivity constraints. The optimization trajectory and endpoints in features and samples spaces for simulated gene expression data is presented on the bottom.

**Fig. 4. Minimal formulations of the Dual Simplex problem**.

a. Dual simplex problem is equivalent to problem of finding single simplex with constraint on its inverse. This formulation demonstrates the dramatic reduction in the number of optimized variables achieved by Dual Simplex approach.

b. Geometrical interpretation of simplexes in features and samples spaces highlights inverse proportionality of their volumes.

c. Gradient descent computational formulation of the minimal formulation of the Dual Simplex problem. Here, only positivity terms (with coefficients λ and β) are optimized as deconvolution term is strictly enforced by the introduction of inverse. Below the optimization result for simulated gene expression data is presented.

**Fig. 5. Application of the dual simplex approach to image data decomposition**.

a. Each mixed image is obtained as a sum of the *K* = *4* main picture components with nonnegative non-constrained coefficients. Noisy scenario involves introduction of the proportional noise with predefined standard deviation representing the noise level.

b. Visualization of the individual pixel points (left) and picture points (right) in respective projection space. Green lines represent true underlying simplex coordinates obtained by projecting rows of *H* and columns of *W* to respective space. For no noise scenario all the points are located within the geometrical simplex.

c. In case of noise present (*noise_level* = *2* in this case) the simplex structure becomes distorted and individual points can be located outside the original geometrical structure.

d. Reconstruction quality metrics measured for matrices *V* (row 1), *H* (row 2) and *W* (row 3) for different noise levels. Quality metrics are defined as median *RMSE* and *1* − *pearson*^*2*^ values measured between rows of predicted matrix *H* and true matrix *H* as well as columns of predicted matrices *W* and *V* in comparison to columns of the true matrices *W* and *V*.

e. Reconstructed images (columns of matrix *W*) visualized for all five methods for the high level of the noise (*noise_level* = *2*).

**Fig. 6. Single cell clustering with dual simplex approach**

a. A tSNE plot of the TCGA single cell HNSC (GSE103322) dataset with main clusters annotated by Puram et al.

b. First six dimensions for both genes (i.e. features) and cells (i.e. samples) projection subspaces (*K* = *16*)upon Sinkhorn transformation applied to filtered expression matrix. Cell populations are clearly separable within this subspace.

c. Dual simplex optimization procedure result with result simplex locations, optimization history and marker genes for the simplex genes.

d. Original tSNE cell points in a space of genes colored by proportion values (matrix *H*) for each of the *K* = *16* main components.

e. Cell cluster labels assigned as component name with maximum proportion value for each individual point.

f. The heatmap of expression profiles for subsets of marker genes specific to main components.

g. Clustering result with different number of main components (*K*)selected.

**Fig. 7. Complete deconvolution of bulk-RNA datasets with dual simplex approach**.

a. Complete deconvolution of GSE19830 dataset with result simplex location and optimization history for both genes and samples spaces (left), extracted marker genes highlighted on a simplex of genes (middle) or heatmap of gene expression from pure lung, brain and liver samples (heatmap). Predicted proportions values for each sample compared to true proportions extracted from the data (right).

b. Complete deconvolution of GSE11058 with result simplex location and optimization history for both genes and samples spaces (left), obtained marker genes highlighted on a simplex of genes (middle). Marker genes transcriptional profile compared to signature genes extracted from pure samples (heatmap). Predicted proportions values for each sample compared to true proportions extracted from the data (right).

c. Complete deconvolution of TCGA bulk-HNSC dataset with result simplex location and optimization history for both genes and samples spaces (left), obtained marker genes highlighted on a simplex of genes (middle). Enrichment of predicted cell types’ marker genes in a major annotation cell subsets extracted from the external GSE181919 dataset (right)

d. Predicted proportions distribution for each of the main components.

e. Enrichment of predicted malignant cell subsets on a UMAP of malignant cells

f. Correlation of predicted proportions with selected clinical features.

**Fig. S1. Projective NMF approach**

a. Representation of the original data matrix both in space of samples and features. Factorization can be solved with two unrelated procedures.

b. First approach is a projection to a hyperplane of genes in a space of samples for row normalized data matrix, followed by identification of vertices of the simplex within this hyperplane, which results in a matrix 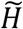

c. First approach is a projection to a hyperplane of samples in a space of samples fir column normalized matrix, followed by identification of vertices of the simplex within this hyperplane, which results in a matrix 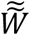

**Fig. S2. Visualization of projection optimization solutions**.

a. SVD vectors delivering minimum to unconstrained optimization problem are not necessarily located within the simplex hyperplane.

b. Centering of the dataset helps to identify two unrelated projection vectors sets located withing the hyperplane, but not necessarily forming orthogonal coordinate systems.

c. Additional orthogonality constraint enforces first projection vectors to be orthogonal to hyperplane. This solution provides orthogonal coordinate systems to represent either of the features and samples simplexes in (*K* − *1*)-dimensional space.

**Fig. S3. Additional NMF experiments**

a. Factorization of the image dataset with increased number of mixtures (*N* = 3*00*) with no noise present. 3D representation of projection points demonstrates predicted simplex geometrical shape for pictures (red) and pixels (blue) spaces along with true simplex shape (green).

b. Factorization of the image dataset with increased number of mixtures (*N* = 3*00*) with *noise_level* = *6*.

c. Factorization result for both clean and noisy randomly generated data matrices *W,H∼U*(*0,50*)containing *M*= *1000* features and *N* = *800* mixtures.

**Fig. S4. Proportion based clustering with dual simplex approach**.

The comparison of cell clusters obtained by dual simplex approach optimization procedure (*DualSimplex column*), geometrical simplex initialization procedure using VCA algorithm (Nascimento & Dias, 2005) applied to Sinkhorn transformed matrix (*VCA column*) and Seurat (Hao et al., 2020) pipeline (*Seurat column*) for different number of clusters (*K*).

**Fig. S5. Further analysis of the bulk-HNCS dataset**.

a. Enrichment of top-100 marker genes for each obtained cell type in pathways extracted from the Molecular Signatures Database (Liberzon et al., 2011).

b. Heatmap of expression values for selected genes for each individual cell type.

c. Assessment of the stability of the convergence for the dataset. Randomized initialization points converge to the same solution points of the simplex.

**Fig. S6. Signature-based deconvolution with dual simplex approach**

a. Signature based deconvolution of computationally mixed brain dataset by Sutton et al (Sutton et al., 2022) with 3 different marker gene sets VL (Velmeshev et al., 2019), NG (Nagy et al., 2020),CA (Hodge et al., 2019). Enrichment of marker genes in (Velmeshev et al., 2019) demonstrated at the top of the figure. These marker genes aligned with the corners of simplex structure of genes which is demonstrated on UMAP of genes (left). Dual simplex performance is comparable to DSA and CIBERSORT methods (Newman et al., 2015; Zhong et al., 2013) (right).

b. Signature based deconvolution of the benchmark GSE11058 dataset with markers extracted from the pure samples (left). Dual simplex approach is able to reconstruct the cell type proportions and gene expression profiles with a low RMSE error.

---

**Supplementary Figure 1.**
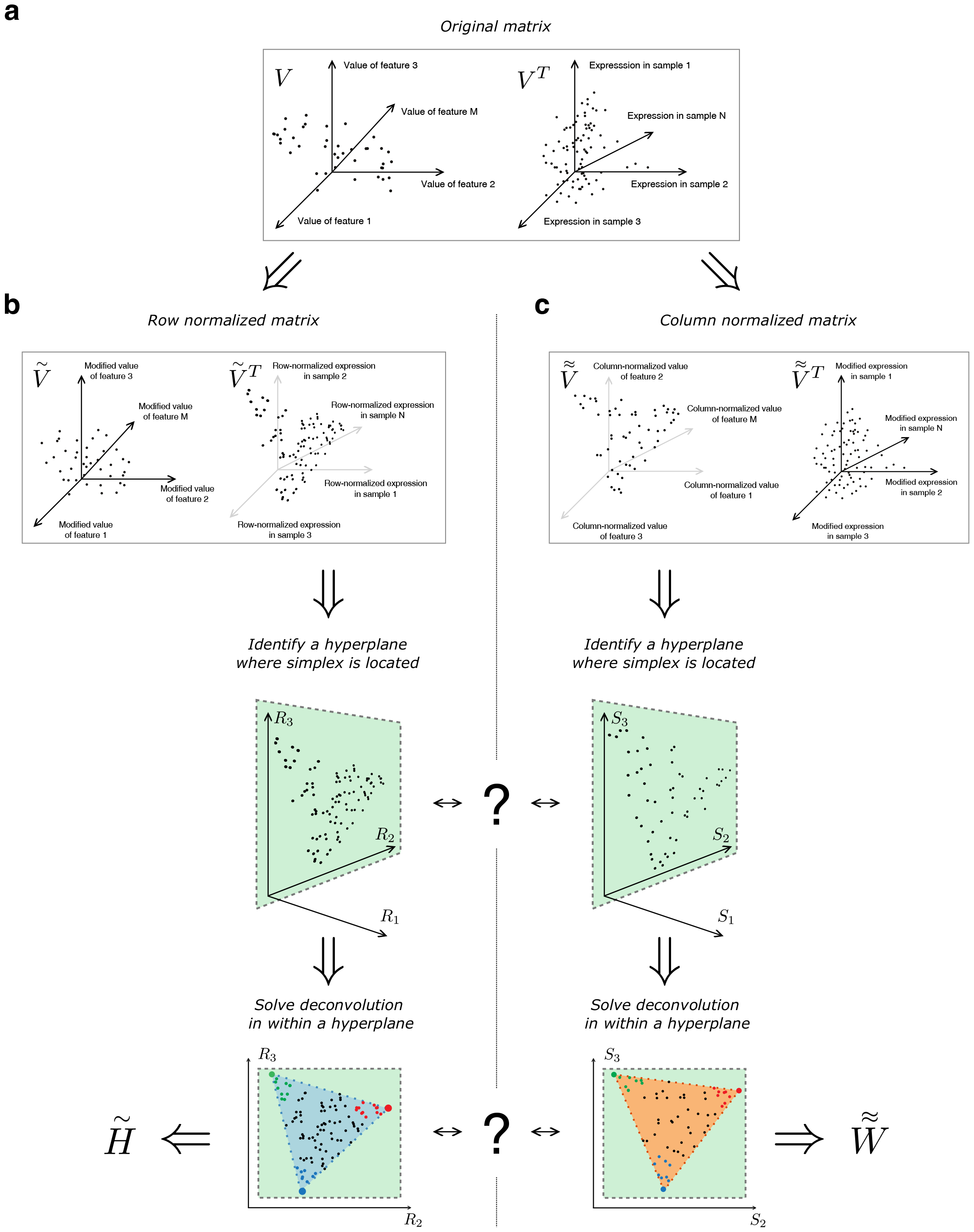
Projective NMF approach. (a) Representation of the original data matrix both in space of samples and features. Factorization can be solved with two unrelated procedures. (b) First approach is a projection to a hyperplane of genes in a space of samples for row normalized data matrix, followed by identification of vertices of the simplex within this hyperplane, which results in a coefficients matrix . (c) First approach is a projection to a hyperplane of samples in a space of samples fir column normalized matrix, followed by identification of vertices of the simplex within this hyperplane, which results in a basis matrix.

**Supplementary Figure 2.**
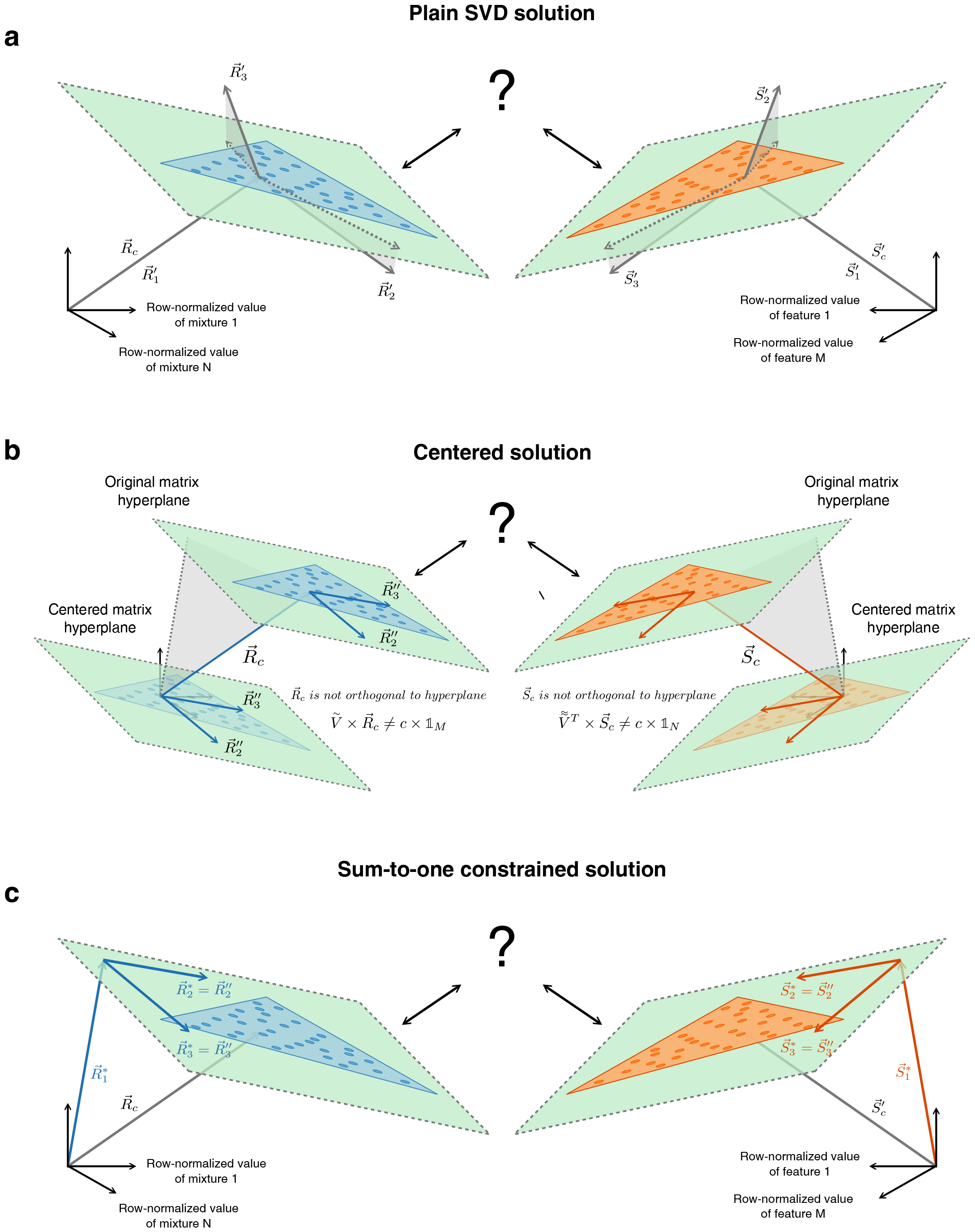
Visualization of projection optimization solutions. (a) SVD vectors delivering minimum to unconstrained optimization problem are not necessarily located within the simplex hyperplane. (b) Centering of the dataset helps to identify two unrelated projection vectors sets located withing the hyperplane, but not necessarily forming orthogonal coordinate systems. (c) Additional orthogonality constraint enforces first projection vectors to be orthogonal to hyperplane. This solution provides orthogonal coordinate systems to represent either of the features and samples simplexes in (K-1)-dimensional space.

**Supplementary Figure 3.**
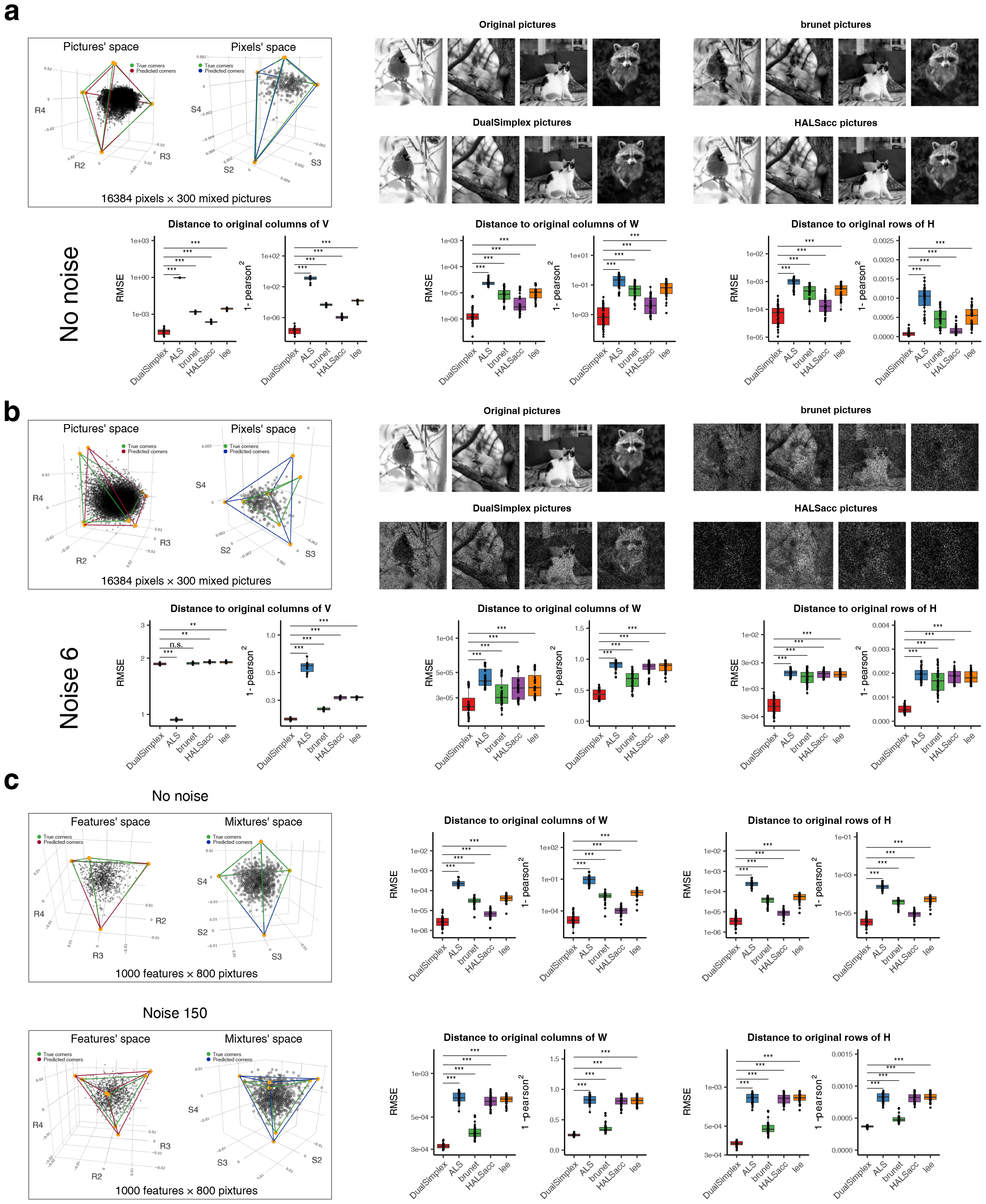
Additional NMF experiments. (a) Factorization of the image dataset with increased number of mixtures (N=300) with no noise present. 3D representation of projection points demonstrates predicted simplex geometrical shape for pictures (red) and pixels (blue) spaces along with true simplex shape (green). (b) Factorization of the image dataset with increased number of mixtures (N=300) with noise_level=6. (c) Factorization result for both clean and noisy randomly generated data matrices W,H∼U(0,50) containing M=1000 features and N=800 mixtures.

**Supplementary Figure 4.**
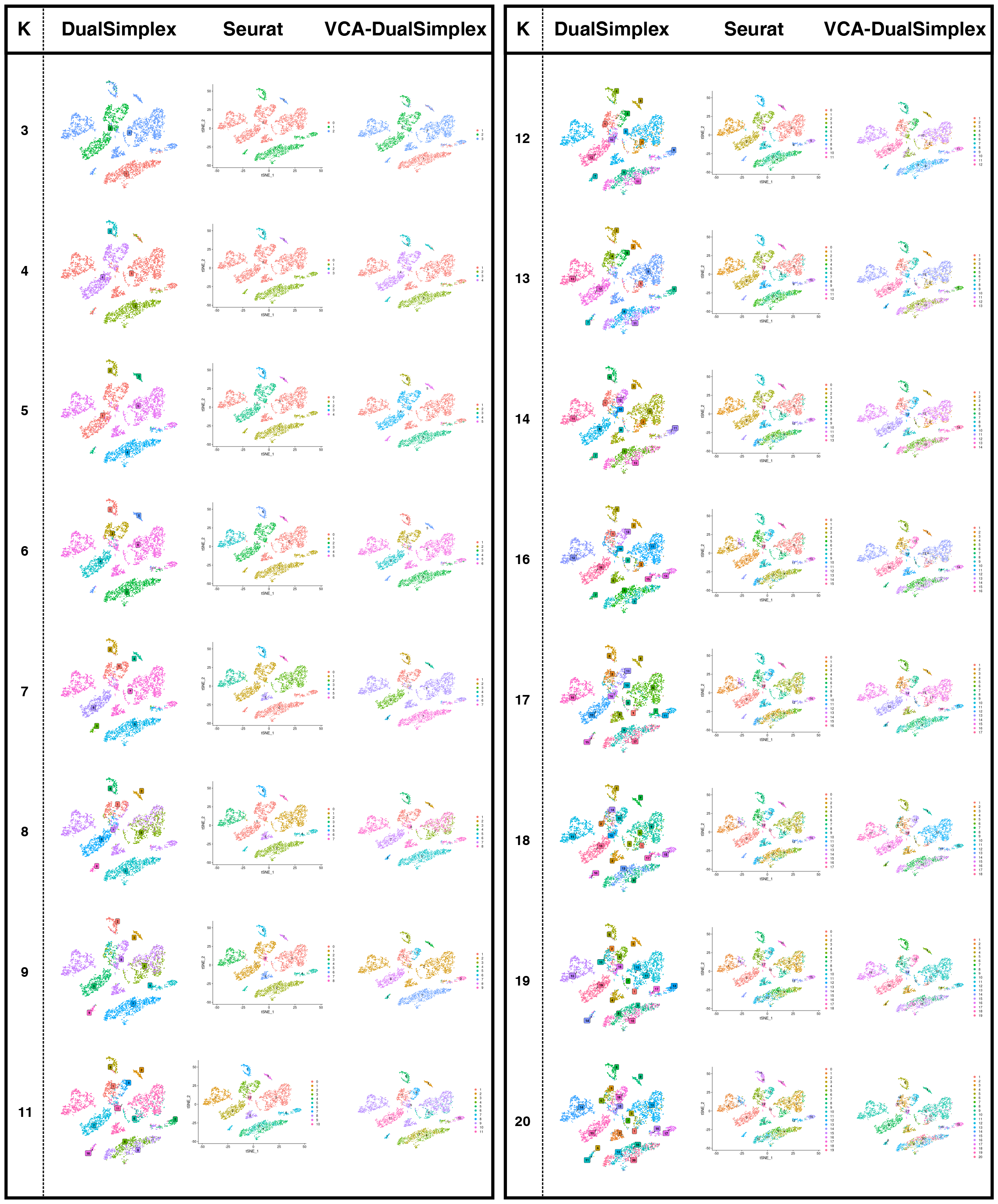
Proportion based clustering with dual simplex approach. The comparison of cell clusters obtained by dual simplex approach optimization procedure (DualSimplex column), geometrical simplex initialization procedure using VCA algorithm (Nascimento & Dias, 2005) applied to Sinkhorn transformed matrix (VCA column) and Seurat (Hao et al., 2020) pipeline (Seurat column) for different number of clusters (K).

**Supplementary Figure 5.**
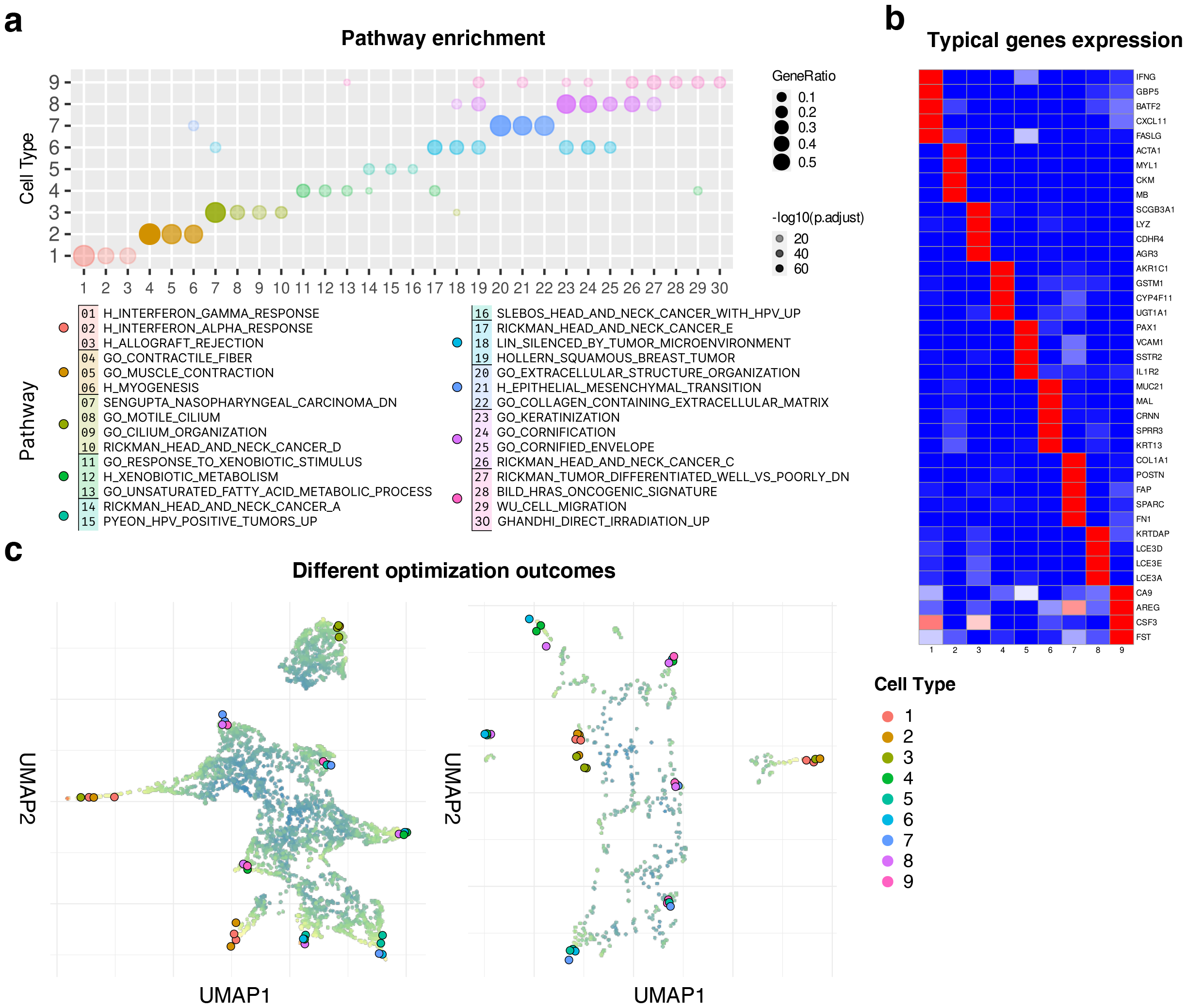
Further analysis of the bulk-HNCS dataset. (a) Enrichment of top-100 marker genes for each obtained cell type in pathways extracted from the Molecular Signatures Database (Liberzon et al., 2011). (b) Heatmap of expression values for selected genes for each individual cell type. (c) Assessment of the stability of the convergence for the dataset. Randomized initialization points converge to the same solution points of the simplex.

**Supplementary Figure 6.**
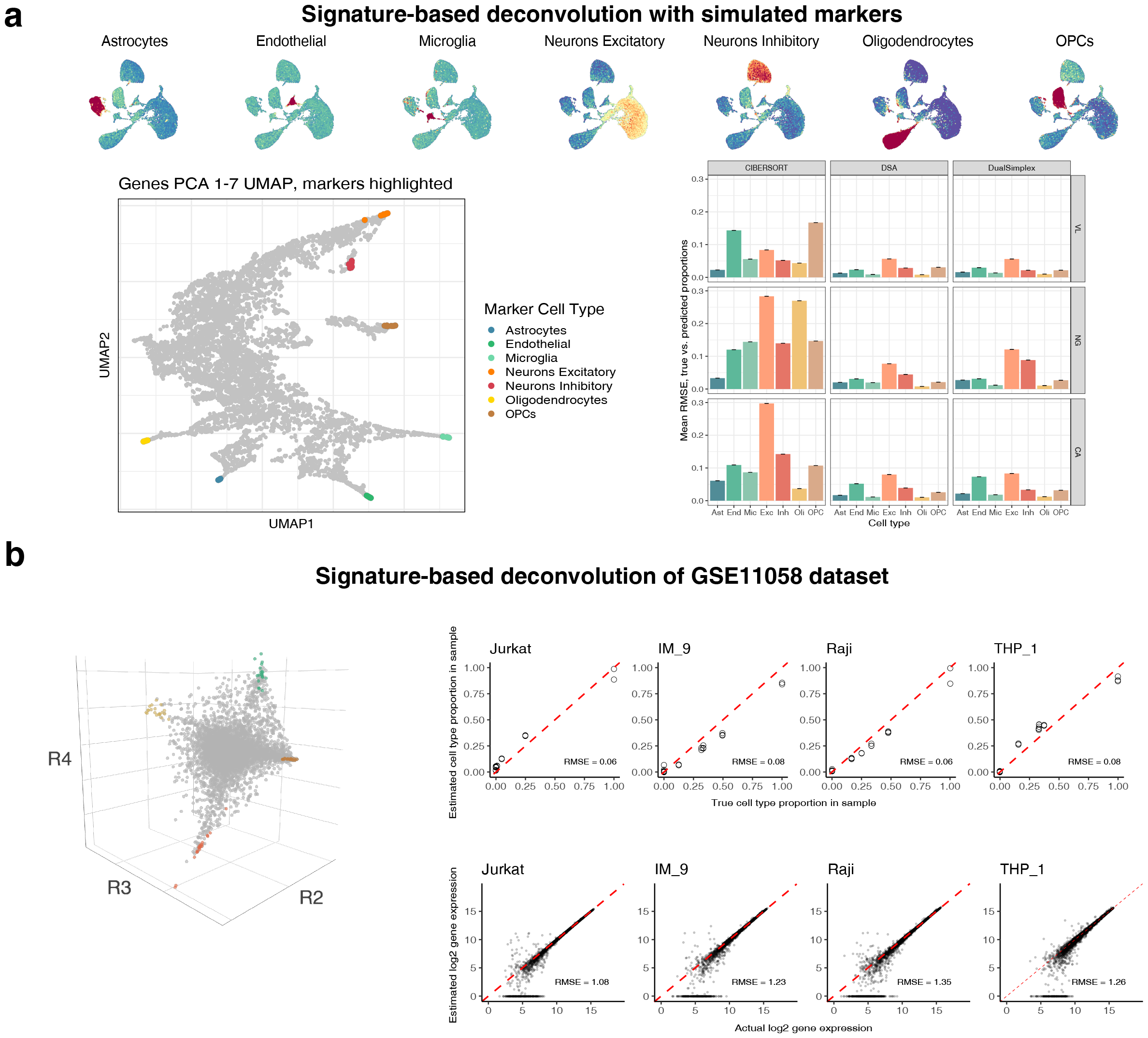
Signature-based deconvolution with dual simplex approach. (a) Signature based deconvolution of computationally mixed brain dataset by Sutton et al (Sutton et al., 2022) with 3 different marker gene sets VL (Velmeshev et al., 2019), NG (Nagy et al., 2020),CA (Hodge et al., 2019). Enrichment of marker genes in (Velmeshev et al., 2019) demonstrated at the top of the figure. These marker genes aligned with the corners of simplex structure of genes which is demonstrated on UMAP of genes (left). Dual simplex performance is comparable to DSA and CIBERSORT methods (Newman et al., 2015; Zhong et al., 2013) (right). (b) Signature based deconvolution of the benchmark GSE11058 dataset with markers extracted from the pure samples (left). Dual simplex approach is able to reconstruct the cell type proportions and gene expression profiles with a low RMSE error.

