## Supplementary Notes for "Non-negative matrix factorization and deconvolution as dual simplex problem"

### Supplementary contents


| | |
|---|---|
| Supplementary Note 1 | The proof of existence of a first simplex in samples space of row normalized matrix |
| Supplementary Note 2 | The proof of existence of a complementary simplex in features space of column normalized matrix |
| Supplementary Note 3 | Definition of Sinkhorn scaling transformation. Relation between two simplexes of Sinkhorn transformed matrix |
| Supplementary Note 4 | Definition of projection operation for $N$ dimensional samples' space to a lower $K$ dimensional space. Vectors $R$. Projective formulation of NMF/deconvolution in this space |
| Supplementary Note 5 | Definition of projection operation for $M$ dimensional features space to a lower $K$ dimensional space. Vectors $S$. Projective formulation of NMF/deconvolution in this space |
| Supplementary Note 6 | $(K-1)$-dimensional structure of a simplex in samples space leads to additional constraint for projection $(R)$ vectors we want to maintain |
| Supplementary Note 7 | $(K-1)$-dimensional structure of a simplex in features space leads to additional constraint for projection $(S)$ vectors we want to maintain |
| Supplementary Note 8 | Definition of the same problems for Sinkhorn transformed matrices. The constant relation allows for joining of two independent projection optimization problems into a single optimization problem. |
| Supplementary Note 9 | Sinkhorn transformed matrices inherently satisfy additional constraints we want to define, so singular vectors are possible solution for projection minimization. The main theorem. |
| Supplementary Note 10 | Gradient descent training formulations to search simplex corners in projected space. Update rules, hyperparameters and initialization strategies. Geometrical relation between simplexes |
| Supplementary Note 11 | Definition of reverse Sinkhorn transformation to return to original space solution matrices $W$ and $H$ |
| Supplementary Note 12 | Application: Guidance on data prefiltering |

### Supplementary Note 1. Normalized and transposed matrix $\widetilde{V}^T$ is a simplex in a samples space

The original data matrix could be represented as multiplication two matrices $W$ – feature matrix of pure components (e.g. cell types) and $H$ – coefficients matrix.

$$V = W \times H,$$

$$v_{i,j} \in \mathbb{R}^{M\times N}, \qquad w_{i.j} \in \mathbb{R}^{M\times K}, \qquad h_{i,j} \in \mathbb{R}^{K\times N}$$

$$K - number\ of\ pure\ components$$
$$N - number\ of\ samples$$
$$M - number\ of\ features$$

Where $v_{i,j}$ describes value of $i^{th}$ feature in $j^{th}$ sample, $w_{i.k}$ describes value of $i^{th}$ feature (e.g. gene expression) in $k^{th}$ pure component (e.g. pure cell type), $h_{k,j} \geq 0$ describes contribution value (e.g. proportion) of $k^{th}$ pure components (e.g. pure cell types),in $j^{th}$ sample.

#### <u>Introduce normalization for matrix *V*</u>

Each element of V could be rewritten as

$$\forall\ i \in [1, M], j \in [1, N]$$
$$v_{i.j} = \sum_{k=1}^{K} w_{i,k} h_{k,j}$$

For both matrices $V$ and $H$ the row normalization operation could be written in a matrix form:

$$\tilde{V} = D_v \times V$$

$$\widetilde{H} = D_h \times H$$

Where $D_v$, $D_h$ are normalizing matrices:

$$D_v = \begin{bmatrix} \frac{1}{\sum_{j=1}^{N} v_{1,j}} & 0 & \cdots & 0 \\ 0 & \frac{1}{\sum_{j=1}^{N} v_{2,j}} & \cdots & 0 \\ \vdots & \vdots & \ddots & \vdots \\ 0 & 0 & \cdots & \frac{1}{\sum_{j=1}^{N} v_{M,j}} \end{bmatrix} \quad D_h = \begin{bmatrix} \frac{1}{\sum_{j=1}^{N} h_{1,j}} & 0 & \cdots & 0 \\ 0 & \frac{1}{\sum_{j=1}^{N} h_{2,j}} & \cdots & 0 \\ \vdots & \vdots & \ddots & \vdots \\ 0 & 0 & \cdots & \frac{1}{\sum_{j=1}^{N} h_{K,j}} \end{bmatrix}$$

So, the original deconvolution equation could be rewritten as

$$\tilde{V} = D_v \times W \times D_h^{-1} \times \widetilde{H}$$

Let's denote:

$$\widetilde{W} = D_v \times W \times D_h^{-1}$$

Then we could rewrite this equation as. Defining the NMF equation for normalized matrix $\tilde{V}$

$$\boxed{\tilde{V} = \widetilde{W} \times \widetilde{H}}$$

Where $\tilde{V}$ and $\widetilde{H}$ are row normalized

$$\forall i \in [1, M] \sum_{j=1}^{N} \tilde{v}_{i,j} = 1 \; and \; \forall k \in [1, K] \sum_{j=1}^{N} \tilde{h}_{k,j} = 1$$

### **Proof of simplex property**

*Statement 1*

Note that $\widetilde{W}$ is also row normalized.

*Proof*

The sum of row $i$ for matrix $\tilde{V}$ could be rewritten in the following way:

$$\forall i \in [1, M] \sum_{j=1}^{N} \tilde{v}_{i.j} = 1 = \sum_{j=1}^{N} \sum_{k=1}^{K} \widetilde{w}_{i,k} \tilde{h}_{k,j} = \sum_{k=1}^{K} \widetilde{w}_{i,k} \sum_{j=1}^{N} \tilde{h}_{k,j} = \sum_{k=1}^{K} \widetilde{w}_{i,k}$$

Which means

$$\forall i \in [1, M], \qquad \sum_{k=1}^{K} \widetilde{w}_{i,k} = 1$$

Therefore all 3 matrices in the original equation are row normalized.

$$\boxed{\underset{row\ normalized}{\tilde{V}} = \underset{row\ normalized}{\widetilde{W}} \times \underset{row\ normalized}{\tilde{H}}}$$

*Statement 2*

Next, we will show that column vectors $(\tilde{v}^T)_{*,i}$ of matrix $\tilde{V}^T$ lie within a simplex.

*Proof*

Each element of $\tilde{V}$ could be represented as a linear combination of elements from $\tilde{H}$, with coefficients from $\widetilde{W}$.

$$\forall \; i \in [1, M], j \in [1, N]$$
$$\tilde{v}_{i.j} = \sum_{k=1}^{K} \widetilde{w}_{i,k} \tilde{h}_{k,j}$$

If we will look at the transposed matrix of $\tilde{V}^T$

$$(\tilde{v}^T)_{j.i} = \tilde{v}_{i.j} = \sum_{k} \widetilde{w}_{i,k} \tilde{h}_{k,j}$$

For each column $i$ of matrix $\tilde{V}^T$ we will use the same set of coefficients $\widetilde{w}_{i,1}, \dots, \widetilde{w}_{i,K}$ which are row elements of matrix $\widetilde{W}$. So, in a vector form each column vector $(\tilde{v}^T)_{*,i}$ will be represented as

$$(\tilde{v}^T)_{*,i} = (\tilde{v}_{i.*})^T = \begin{bmatrix} \sum_{k} \widetilde{w}_{i,k} * \tilde{h}_{k,1} \\ \vdots \\ \sum_{k} \widetilde{w}_{i,k} * \tilde{h}_{k,N} \end{bmatrix} = \widetilde{w}_{i,1} \begin{bmatrix} \tilde{h}_{1,1} \\ \vdots \\ \tilde{h}_{1,N} \end{bmatrix} + \widetilde{w}_{i,2} \begin{bmatrix} \tilde{h}_{2,1} \\ \vdots \\ \tilde{h}_{2,N} \end{bmatrix} + \dots + \widetilde{w}_{i,K} \begin{bmatrix} \tilde{h}_{K,1} \\ \vdots \\ \tilde{h}_{K,N} \end{bmatrix}$$

Note that coefficients of this representation are summing to 1 (since $\widetilde{W}$ is row-normalized).

$$\sum_{k=1}^{K} \widetilde{w}_{i,k} = 1$$

We also know that $\widetilde{w}_{i,k} > 0$ (by its definition)

### Summary

Therefore

We defined the NMF equation for row normalized matrix.

$$\boxed{\underset{row\ normalized}{\widetilde{V}} = \underset{row\ normalized}{\widetilde{W}} \times \underset{row\ normalized}{\widetilde{H}}}$$

Which holds simplex property:

$$\forall i \in [1, M], \forall j \in [1, N]\ \exists\ \left(\alpha_{i,1}, \dots, \alpha_{i,K}\right)$$
$$(\tilde{v}^T)_{*,i} = \sum_k \alpha_{i,k} \left(\tilde{h}^T\right)_{*,k}$$
$$\forall i \in [1, M], \forall k \in [1, K]\ \alpha_{i.k} = \widetilde{w}_{i,k} > 0\ and \sum_k \alpha_{i.k} = 1$$
$$(\tilde{v}^T)_{*,i} = \ (\tilde{v}_{i.*})^T = \begin{bmatrix} \sum_k \alpha_{i,k} * \tilde{h}_{k,1} \\ \vdots \\ \sum_k \alpha_{i,k} * \tilde{h}_{k,N} \end{bmatrix} = \alpha_{i,1} \begin{bmatrix} \tilde{h}_{1,1} \\ \vdots \\ \tilde{h}_{1,N} \end{bmatrix} + \ \alpha_{i,2} \begin{bmatrix} \tilde{h}_{2,1} \\ \vdots \\ \tilde{h}_{2,N} \end{bmatrix} + \dots + \ \alpha_{i,K} \begin{bmatrix} \tilde{h}_{K,1} \\ \vdots \\ \tilde{h}_{K,N} \end{bmatrix}$$

To sum up, what we have is $M$ column vectors in $N$ dimensional space which form a transposed matrix $\tilde{V}^T$. Each individual column vector of this matrix could be represented as a linear combination of $K$ column vectors ($N$ dimensional vectors) of transposed matrix $\widetilde{H}^T$. And all the coefficients $\alpha_{i,k}$ of this representation are positive and sum to $1$. This means that $\tilde{V}^T$ is a $(K-1)$ dimensional simplex and vectors $\left(\tilde{h}^T\right)_{*,k}$ define corners of this simplex.

### Supplementary Note 2. Existence of complementary simplex in a features space of column normalized matrix

What we did so far is we defined NMF equation.

$$V = W \times H,$$

$$v_{i,j} \in \mathbb{R}^{M \times N}, \qquad w_{i.j} \in \mathbb{R}^{M \times K}, \qquad h_{i,j} \in \mathbb{R}^{K \times N}$$

$$\begin{aligned} &K - number\ of\ pire\ components \\ &N - number\ of\ samples \\ &M - number\ of\ features \end{aligned}$$

And the new NMF equation for row normalized matrices $\tilde{V} = D_v \times V$

$$\underset{row\ normalized}{\tilde{V}} = \underset{row\ normalized}{\widetilde{W}} \times \underset{row\ normalized}{\widetilde{H}}$$

#### <u>Introduce column normalization for matrix *V*</u>

Now let's column normalize matrix $V$

And introduce a new matrix $\tilde{\tilde{V}}$

$$\tilde{\tilde{V}} = \tilde{V} \times D_v - column\ normalized\ matrix$$

Where $D_v$ is a diagonal matrix that column normalizes $\tilde{V}$

$$D_{v_1} = \begin{bmatrix} \frac{1}{\sum_{i=1}^{M} v_{i,1}} & 0 & \cdots & 0 \\ 0 & \frac{1}{\sum_{i=1}^{M} v_{i,2}} & \cdots & 0 \\ \vdots & \vdots & \ddots & \vdots \\ 0 & 0 & \cdots & \frac{1}{\sum_{i=1}^{M} v_{i,N}} \end{bmatrix}$$

Similarly, we can define $D_w$ which will column normalize $W$

$$\tilde{\tilde{W}} = W \times D_w$$

Using the same original deconvolution equation, we can rewrite this new matrix in a similar form through $\tilde{\tilde{W}}$ and $\tilde{\tilde{H}}$, forming the NMF formulation for column normalized matrices.

$$\tilde{\tilde{V}} = W \times H \times D_{v_1} = \underbrace{W \times D_w}_{\tilde{\tilde{W}}} \times \underbrace{D_w^{-1} \times H \times D_v}_{\tilde{\tilde{H}}} = \tilde{\tilde{W}} \times \tilde{\tilde{H}}$$

$$\boxed{\tilde{\tilde{V}} = \tilde{\tilde{W}} \times \tilde{\tilde{H}}}$$

Where $\tilde{\tilde{V}}$ and $\tilde{\tilde{W}}$ are column normalized

$$\forall j \in [1, N] \sum_{i=1}^{M} \tilde{\tilde{v}}_{i.j} = 1\ and\ \forall k \in [1, K] \sum_{i=1}^{M} \tilde{\tilde{w}}_{i.k} = 1$$

### Proof of simplex property

*Statement 1*

Note that $\tilde{\tilde{H}}$ is also column normalized.

*Proof*

The sum of column for matrix $\tilde{\tilde{V}}$ could be rewritten in the following way:

$$\forall j \in [1, N] \sum_{i=1}^{M} \tilde{\tilde{v}}_{i.j} = \sum_{i=1}^{M}\sum_{k=1}^{K} \tilde{\tilde{w}}_{i.k}\tilde{\tilde{h}}_{k.j} = \sum_{k=1}^{K} \tilde{\tilde{h}}_{k.j} \sum_{i=1}^{M} \tilde{\tilde{w}}_{i.k} = \sum_{k=1}^{K} \tilde{\tilde{h}}_{k.j} = 1$$

(i.e., $\tilde{\tilde{H}}$ is column normalized)

Therefore all 3 matrices in the original equation are column normalized.

$$\underset{column\ normalized}{\tilde{\tilde{V}}} = \underset{column\ normalized}{\tilde{\tilde{W}}} \times \underset{column\ normalized}{\tilde{\tilde{H}}}$$

*Statement 2*

Next, we will show that column vectors $\tilde{\tilde{v}}_{*,j}$of matrix $\tilde{\tilde{V}}$ lie within a simplex.

*Proof*

Each element of $\tilde{\tilde{V}}$ could be represented as a linear combination of elements from $\tilde{\tilde{W}}$ , with coefficients from $\tilde{\tilde{H}}$.

$$\forall i \in [1, M], \forall j \in [1, N]$$
$$\tilde{\tilde{v}}_{i,j} = \sum_{k=1}^{K} \tilde{\tilde{w}}_{i.k}\tilde{\tilde{h}}_{k.j}$$

For each column $j$ of matrix $\tilde{\tilde{V}}$ we will use the same set of coefficients $\tilde{\tilde{h}}_{1,j}, \dots, \tilde{\tilde{h}}_{K,j}$ which are column elements of matrix $\tilde{\tilde{H}}$. So, in a vector form each column vector $\tilde{\tilde{v}}_{*,j}$ will be represented as

$$\tilde{\tilde{v}}_{*,j} = \begin{bmatrix} \sum_{k} \tilde{\tilde{w}}_{1,k} * \tilde{\tilde{h}}_{k,j} \\ \vdots \\ \sum_{k} \tilde{\tilde{w}}_{M,k} * \tilde{\tilde{h}}_{k,j} \end{bmatrix} = \tilde{\tilde{h}}_{1,j} \begin{bmatrix} \tilde{\tilde{w}}_{1,1} \\ \vdots \\ \tilde{\tilde{w}}_{M,1} \end{bmatrix} + \tilde{\tilde{h}}_{2,j} \begin{bmatrix} \tilde{\tilde{w}}_{1,2} \\ \vdots \\ \tilde{\tilde{w}}_{M,2} \end{bmatrix} + \dots + \tilde{\tilde{h}}_{K,j} \begin{bmatrix} \tilde{\tilde{w}}_{1,K} \\ \vdots \\ \tilde{\tilde{w}}_{M,K} \end{bmatrix}$$

Note that coefficients of this representation are summing to 1 (since $\tilde{\tilde{H}}$ is column-normalized).

$$\sum_{k=1}^{K} \tilde{\tilde{h}}_{k.j} = 1$$

### Note: This property is true for consecutive normalizations

Since the simplex property is true for any positive and factorizable matrix, the consecutive application of normalization still holds this property.

We will denote previously defined $D_v$ as $D_{v_0}$:

$$\tilde{V} = D_v \times V = D_{v_0} \times V - row\ normalized\ matrix$$

Now let's column normalize matrix $\tilde{V}$(previously row normalized $V$)

$$\tilde{\tilde{V}} = \tilde{V} \times D_{v_1} = D_{v_0} \times V \times D_{v_1} - column\ normalized\ matrix$$

Where $D_{v_1}$ is a diagonal matrix that column normalizes $\tilde{V}$

$$D_{v_1} = \begin{bmatrix} \frac{1}{\sum_{i=1}^{M} \tilde{v}_{i,1}} & 0 & \dots & 0 \\ 0 & \frac{1}{\sum_{i=1}^{M} \tilde{v}_{i,2}} & \dots & 0 \\ \vdots & \vdots & \ddots & \vdots \\ 0 & 0 & \dots & \frac{1}{\sum_{i=1}^{M} \tilde{v}_{i,N}} \end{bmatrix}$$

Similarly, we can define $D_w$ which will column normalize $\widetilde{W}$

$$\tilde{\widetilde{W}} = \widetilde{W} \times D_w$$

Using the same original deconvolution equation, we can rewrite this new matrix in a similar form through new matrices $\tilde{\widetilde{W}}$ and $\tilde{\tilde{H}}$.

$$\tilde{\tilde{V}} = \widetilde{W} \times \tilde{H} \times D_{v_1} = \underbrace{\widetilde{W} \times D_w}_{\tilde{\widetilde{W}}} \times \underbrace{D_w^{-1} \times \tilde{H} \times D_{v_1}}_{\tilde{\tilde{H}}} = \tilde{\widetilde{W}} \times \tilde{\tilde{H}}$$

The same simplex property is true.

### Summary

Therefore

We defined the NMF equation for column normalized matrix.

$$\boxed{\underset{column\ normalized}{\tilde{\tilde{V}}} = \underset{column\ normalized}{\tilde{\widetilde{W}}} \times \underset{column\ normalized}{\tilde{\tilde{H}}}}$$

Which holds simplex property:

$$\forall i \in [1, M], \forall j \in [1, N]\ \exists\ (\alpha_{1,j}, \dots, \alpha_{K,j})$$
$$\tilde{\tilde{v}}_{*,j} = \sum_k \alpha_{k,j} \tilde{\tilde{w}}_{*,k}$$
$$\forall j \in [1, N]\ \forall k \in [1, K]\ \alpha_{k,j} = \tilde{\tilde{h}}_{k,j} > 0\ and \sum_k \alpha_{k,j} = 1$$
$$\tilde{\tilde{v}}_{*,j} = \begin{bmatrix} \sum_k \tilde{\tilde{w}}_{1,k} * \alpha_{k,j} \\ \vdots \\ \sum_k \tilde{\tilde{w}}_{M,k} * \alpha_{k,j} \end{bmatrix} = \alpha_{1,j} \begin{bmatrix} \tilde{\tilde{w}}_{1,1} \\ \vdots \\ \tilde{\tilde{w}}_{M,1} \end{bmatrix} + \alpha_{2,j} \begin{bmatrix} \tilde{\tilde{w}}_{1,2} \\ \vdots \\ \tilde{\tilde{w}}_{M,2} \end{bmatrix} + \dots + \alpha_{K,j} \begin{bmatrix} \tilde{\tilde{w}}_{1,K} \\ \vdots \\ \tilde{\tilde{w}}_{M,K} \end{bmatrix}$$

To sum up, what we have is $N$ column vectors in $M$ dimensional space which form a matrix $\tilde{\tilde{V}}$. Each individual column vector of this matrix could be represented as a linear combination of

$K$ column vectors ($M$ dimensional vectors) of matrix $\tilde{\tilde{W}}$. And all the coefficients $\alpha_{j,k}$ of this representation will sum to 1. Which means that $\tilde{\tilde{V}}$ is a $(K-1)$ dimensional simplex and vectors $\tilde{\tilde{w}}_{*,k}$ define corners of this simplex.

### Supplementary Note 3. Introducing Sinkhorn transformation operation

#### What we did so far

*Defined the NMF problem for normalized matrices* $\tilde{\mathbf{V}}$ *in a samples space and* $\tilde{\tilde{\mathbf{V}}}$ *in a features' space:*

$$V = W \times H \Rightarrow \tilde{V} = \widetilde{W} \times \widetilde{H} \Rightarrow \tilde{\mathbf{V}}^{\boldsymbol{T}} = \widetilde{\boldsymbol{H}}^{\boldsymbol{T}} \times \widetilde{\boldsymbol{W}}^{\boldsymbol{T}} \Rightarrow \tilde{\tilde{\boldsymbol{V}}} = \widetilde{\widetilde{\boldsymbol{W}}} \times \tilde{\tilde{\boldsymbol{H}}}$$

$$v_{i,j} \in \mathbb{R}^{M \times N}, \qquad w_{i.j} \in \mathbb{R}^{M \times K}, \qquad h_{i,j} \in \mathbb{R}^{K \times N}$$

$$K - number\ of\ pure\ components$$
$$N - number\ of\ samples$$
$$M - number\ of\ features$$

$$H - column\ normalized\ matrix$$
$$\tilde{V}, \widetilde{W}, \widetilde{H} - row\ normalized\ matrices$$
$$\tilde{V}^T, \widetilde{H}^T, \widetilde{W}^T - column\ normalized\ matrices$$
$$\widetilde{\widetilde{W}}, \tilde{\tilde{V}}, \tilde{\tilde{H}} - row\ normalized\ matrices$$

We also showed that points of these to matrices lie on two simplexes.

#### Subsequent normalizations reveal relation between matrices in features and samples space.

What we did so far is we took the original matrix $V$ and performed row normalization followed by column normalization.

$$\tilde{\tilde{V}} = \tilde{V} \times D_{v_1} = D_{v_0} \times V \times D_{v_1}$$

And showed that these two formulations for simplexes, even for consecutive normalizations.

First, we notice that if we will again row normalize matrix $\tilde{\tilde{V}}$ we will again get a simplex in a samples space. And if we will column normalize result of this operation, we will get a simplex in a features space. i.e. consecutive application of the row normalization – column normalization preserves simplex structures in corresponding space. Therefore, we wonder what happens if one considers repetitive application of these procedures ad infimum:

$$\underbrace{D_{v_0} \times V \times D_{v_1}}_{\substack{siplex\ in \\ features\ space}} \overset{\substack{row \\ normalization}}{\Longrightarrow} \underbrace{D_{v_3} \times D_{v_0} \times V \times D_{v_1}}_{\substack{simlex\ in \\ samples\ space}} \overset{\substack{column \\ normalization}}{\Longrightarrow} \underbrace{D_{v_2} \times D_{v_0} \times V \times D_{v_1} \times D_{v_3}}_{\substack{simplex\ in \\ features\ space}}$$

We can then define the iterative process of normalizations.

$$V \overset{\substack{row \\ norm}}{\Longrightarrow} \underbrace{\boldsymbol{D}_{\boldsymbol{v_0}} \times V}_{\substack{(1) \\ \tilde{V}\ or\ V \\ simplex\ in \\ samples\ space}} \overset{\substack{column \\ norm}}{\Longrightarrow} \underbrace{D_{v_0} \times V \times \boldsymbol{D}_{\boldsymbol{v_1}}}_{\substack{(2) \\ \tilde{\tilde{V}}\ or\ V \\ simplex\ in \\ features\ space}} \overset{\substack{row \\ norm}}{\Longrightarrow} \ldots$$

$$\dots \overset{\substack{column\\norm}}{\Longrightarrow} \underbrace{D_{v_{2n-2}} \times D_{v_{2n-4}} \dots \times D_{v_0} \times V \times D_{v_1} \times \dots \times D_{v_{2n-3}} \times \boldsymbol{D_{v_{2n-1}}}}_{\substack{\overset{(2n)}{V}\\ simplex\ in\\ features\ space}} \overset{\substack{row\\norm}}{\Longrightarrow} \dots$$

Which could be represented in iterative form for even and odd elements:

$$\begin{array}{|ll|}\hline & \overset{(0)}{V} = V \\ \forall i \in [1, 3, \dots 2n-1] & \overset{(i)}{V} = D_{v_{i-1}} \times \overset{(i-1)}{V} - row\ normalized \\ \forall i \in [2, 4, \dots 2n] & \overset{(i)}{V} = \overset{(i-1)}{V} \times D_{v_{i-1}} - column\ normalized \\ \hline \end{array}$$

According to Sinkhorn-Knopp theorem(Sinkhorn, 1967), for any strictly positive square matrix the result of these operations will converge to some matrix, which called doubly stochastic matrix. Which will have both rows and columns normalized.

For non-square matrices we are not guaranteed to have this property. But the process of iteratively scaling the rows and columns of $M \times N$ matrix $V$ to have row and column sums respectively $r_i$ and $c_j$will converge to two subsequences. First subsequence, in which column sums are scaled will converge to the matrix $D_1 \times V \times D_2$. Another subsequence, in which row sums are scaled will converge to $\frac{1}{\mu} D_1 \times V \times D_2$ (Sinkhorn, 1967)

Where

$$\mu = \frac{\sum_{j=1}^{N} c_j}{\sum_{i=1}^{M} r_i}$$

In our case $r_i = 1$ and $c_j = 1$

Then

$$\mu = \frac{\sum_{j=1}^{N} c_j}{\sum_{i=1}^{M} r_i} = \frac{N}{M}$$

*This means that we can get a proportionality relation between two subsequences during the Sinkhorn scaling procedure applied to the data matrix.*

Let's denote as $\overset{\infty}{V_{gs}}$ and $\overset{\infty}{V_{ss}}$ the result of these iterations after a sufficient number of steps $2n$ to reach convergence for both subsequences ($n$ row normalizations and $n$ column normalizations):

$$V \overset{\substack{row\\norm}}{\Longrightarrow} \dots \overset{\substack{row\\norm}}{\Longrightarrow} \underbrace{\boldsymbol{D_{v_{2n-2}}} \times D_{v_{2n-4}} \times \dots \times D_{v_0} \times V \times D_{v_1} \times \dots \times D_{v_{2n-3}}}_{\substack{\overset{(2n-1)}{V}\ or\ \overset{\infty}{V_{ss}}\\ simplex\ in\\ samples\ space}} \overset{\substack{column\\norm}}{\Longrightarrow}$$

$$\overset{\substack{column\\norm}}{\Longrightarrow} \underbrace{D_{v_{2n-2}} \times D_{v_{2n-4}} \times \dots \times D_{v_0} \times V \times D_{v_1} \times \dots \times D_{v_{2n-3}} \times \boldsymbol{D_{v_{2n-1}}}}_{\substack{\overset{(2n)}{V}\ or\ \overset{\infty}{V_{fs}}\\ simplex\ in\\ features\ space}} \overset{\substack{row\\norm}}{\Longrightarrow}$$

$$\overset{row}{\underset{norm}{\Longrightarrow}} \underset{\substack{simplex\ in\\ sample\ space}}{\overset{\infty}{V_{ss}}} \overset{column}{\underset{norm}{\Longrightarrow}} \underset{\substack{simplex\ in\\ features\ space}}{\overset{\infty}{V_{fs}}} \dots \overset{row}{\underset{norm}{\Longrightarrow}} \underset{\substack{simplex\ in\\ sample\ space}}{\overset{\infty}{V_{ss}}} \overset{column}{\underset{norm}{\Longrightarrow}} \underset{\substack{simplex\ in\\ features\ space}}{\overset{\infty}{V_{fs}}}$$

Where $\overset{\infty}{V_{ss}}$ and $\overset{\infty}{V_{fs}}$ are both $M \times N$ matrices.

Then

$$\overset{\infty}{V_{gs}} = D_1 \times V \times D_2$$

And

$$\overset{\infty}{V_{ss}} = \frac{M}{N} D_1 \times V \times D_2$$

Where $D_1 = D_{v_{2n-2}} \times D_{v_{2n-4}} \times \dots \times D_{v_0}$ and $D_2 = D_{v_1} \times \dots \times D_{v_{2n-3}} \times D_{v_{2n-1}}$ , and are diagonal matrices (according to Sinkhorn-Knopp theorem)

Which means that

$$\boxed{\overset{\infty}{V_{ss}} \times \frac{N}{M} = \overset{\infty}{V_{fs}}}$$

Note that normalizing matrix will be the following.

$$D_V = \begin{bmatrix} \frac{N}{M} & \dots & 0 \\ \vdots & \ddots & \vdots \\ 0 & \dots & \frac{N}{M} \end{bmatrix}$$

Similarly, to what we already showed we can modify initial matrices $H$ to keep desired normalization property for each iteration step.

$$H \Longrightarrow \underbrace{\boldsymbol{D_{h_0}} \times H}_{\substack{\overset{(1)}{H} \ or\ \tilde{H} \\ row\ normalized}} \Longrightarrow \underbrace{\boldsymbol{D_{w_1}^{-1}} \times D_{h_0} \times H \times \boldsymbol{D_{v_1}}}_{\substack{\overset{(2)}{H} \ or\ \tilde{\tilde{H}} \\ column\ normalized}} \Longrightarrow \cdots$$

$$\dots \Longrightarrow \underbrace{\boldsymbol{D_{h_{2n-2}}} \times D_{w_{2n-1}}^{-1} \times \dots \times D_{h_2} \times D_{w_1}^{-1} \times D_{h_0} \times H \times D_{v_1} \times D_{v_3} \times \dots \times D_{v_{2n-3}}}_{\substack{\overset{(2n-1)}{H} \ or\ \overset{\infty}{H_{ss}} \\ row\ normalized}} \Longrightarrow$$

$$\Longrightarrow \underbrace{\boldsymbol{D_{w_{2n-1}}^{-1}} \times D_{h_{2n-2}} \times \dots \times D_{h_2} \times D_{w_2}^{-1} \times D_{h_0} \times H \times D_{v_1} \times D_{v_3} \times \dots \times \boldsymbol{D_{v_{2n-1}}}}_{\overset{(2n)}{H} \ or\ \overset{\infty}{H_{fs}}}$$

As well as for $W$ matrix

$$W \Longrightarrow \underbrace{\boldsymbol{D_{v_0}} \times W \times \boldsymbol{D_{h_0}^{-1}}}_{\substack{\overset{(1)}{H} \ or\ \tilde{H} \\ row\ normalized}} \Longrightarrow \underbrace{D_{v_0} \times W \times D_{h_0}^{-1} \times \boldsymbol{D_{w_1}}}_{\substack{\overset{(2)}{H} \ or\ \tilde{\tilde{H}} \\ column\ normalized}} \Longrightarrow \cdots$$

$$\dots \Longrightarrow \underbrace{\boldsymbol{D_{v_{2n-2}}} \times D_{v_{2n-4}} \times \dots \times D_{v_4} \times D_{v_0} \times W \times D_{h_0}^{-1} \times D_{w_2} \times D_{h_3}^{-1} \times \dots \times \boldsymbol{D_{h_{2n-1}}^{-1}}}_{\substack{\overset{(2n-1)}{W} \ or\ \overset{\infty}{W_{fs}} \\ row\ normalized}} \Longrightarrow$$

$$\Longrightarrow \underbrace{D_{v_{2n-2}} \times D_{v_{2n-4}} \times \ldots \times D_{v_4} \times D_{v_0} \times W \times D_{h_0}^{-1} \times D_{w_2} \times D_{h_3}^{-1} \times \ldots \times D_{h_{2n-1}}^{-1} \times \boldsymbol{D}_{\boldsymbol{w_{2n}}}}_{\overset{(2n)}{W} \ or \ \overset{\infty}{W}_{fs}}$$

Which could be represented in iterative form for even and odd elements:

| | | | |
|---|---|---|---|
| | $\overset{(0)}{H} = H$ | $\overset{(0)}{W} = W$ | |
| $\forall i \in [1, 3, \ldots 2n-1]$ | $\overset{(i)}{H} = \boldsymbol{D}_{\boldsymbol{h_{i-1}}} \times \overset{(i-1)}{H}$ | $\overset{(i)}{W} = \boldsymbol{D}_{\boldsymbol{v_i}} \times \overset{(i-1)}{W} \times \boldsymbol{D}_{\boldsymbol{h_i}}^{-\boldsymbol{1}}$ | $-row\ normalized$ |
| $\forall i \in [2, 4, \ldots 2n]$ | $\overset{(i)}{H} = \boldsymbol{D}_{\boldsymbol{w_{i-1}}}^{-\boldsymbol{1}} \times \overset{(i-1)}{H} \times \boldsymbol{D}_{\boldsymbol{v_{i-1}}}$ | $\overset{(i)}{W} = \overset{(i-1)}{W} \times \boldsymbol{D}_{\boldsymbol{w_1}}$ | $-column\ normalized$ |

We can get the relation between $\overset{\infty}{W}_{ss}$ and $\overset{\infty}{W}_{fs}$

$$\overset{\infty}{V}_{fs} = \overset{\infty}{V}_{ss} \times D_V = \underbrace{\overset{\infty}{W}_{ss} \times \overset{\infty}{D}_w}_{\overset{\infty}{W}_{fs}} \times \underbrace{\overset{\infty}{D}_w^{-1} \times \overset{\infty}{H}_{ss} \times D_V}_{\overset{\infty}{H}_{fs}}$$

Where

$$\overset{\infty}{W}_{fs} = \overset{\infty}{W}_{ss} \times \overset{\infty}{D}_w$$

$$\overset{\infty}{W}_{ss} = \overset{\infty}{W}_{fs} \times \overset{\infty}{D}_w^{-1}$$

Where $\overset{\infty}{D}_w^{-1}$ is some diagonal matrix which is not know by default but will be calculated during algorithm. Let's denote this matrix as $D$

This indicates that Sinkhorn scaled expression matrix has very interesting properties – (1) it preserves geometric simplex structure in both features space and in samples space, (2) result matrices in each space are proportional up to scalar coefficient.

### Supplementary Note 4. Projective formulation of the NMF problem in a samples' space (Vectors $\boldsymbol{R}$)

#### What we did so far

*Defined the NMF problem:*

$$V = W \times H$$

$$v_{i,j} \in \mathbb{R}^{M\times N}, \qquad w_{i.j} \in \mathbb{R}^{M\times K}, \qquad h_{i,j} \in \mathbb{R}^{K\times N}$$

$$K - number\ of\ pure\ components$$
$$N - number\ of\ samples$$
$$M - number\ of\ features$$

Where $v_{i,j}$ describes value of $i^{th}$ feature in $j^{th}$ sample, $w_{i.k}$ describes value of $i^{th}$ feature (e.g. gene expression) in $k^{th}$ pure component (e.g. pure cell type), $h_{k,j} \geq 0$ describes contribution value (e.g. proportion) of $k^{th}$ pure components (e.g. pure cell types),in $j^{th}$ sample.

*Reformulated deconvolution problem for normalized matrix $\widetilde{\boldsymbol{V}}$ in a sample space*

We defined new deconvolution equations for normalized matrices.

$$V = W \times H \Rightarrow \tilde{V} = \widetilde{W} \times \widetilde{H} \Rightarrow \widetilde{\mathbf{V}}^{\boldsymbol{T}} = \widetilde{\boldsymbol{H}}^{\boldsymbol{T}} \times \widetilde{\boldsymbol{W}}^{\boldsymbol{T}}$$

$$H - column\ normalized\ matrix$$
$$\tilde{V}, \widetilde{W}, \widetilde{H} - row\ normalized\ matrices$$
$$\tilde{\mathrm{V}}^T, \widetilde{H}^T, \widetilde{W}^T - column\ normalized\ matrices$$

#### We can formulate the optimization problem to find these new NMF matrices.

In terms of optimization, our main goal is to find such matrices $\widetilde{H}$ and $\widetilde{W}$ so that the combination of these matrices will give reconstruct the initial data matrix with lowest possible error.

$$\min_{\widetilde{H}^T, \widetilde{W}^T} \left\| \tilde{V}^T - \widetilde{H}^T \times \widetilde{W}^T \right\|_{\substack{s.t.\ \ \widetilde{H}^T > 0 \\ \widetilde{W}^T > 0}}$$

Note that this formulation is equivalent to deconvolution problem, since $\widetilde{W}^T$ is column normalized. Therefore we might use the NMF and deconvolution terms interchangeably in the following section.

#### Note: $\widetilde{\boldsymbol{V}}^{\boldsymbol{T}}$ is projectable onto $\boldsymbol{K}$ dimensional space

In our work we assume that original data matrix is a mixture of $K$ different data matrices which represent main components of the original data matrix $V$. And these components are pure components.

We assume that $K < M \ll N$ so that.

$$rank(V) = \min\big(rank(W), rank(H)\big) = \min(K, K) = K$$

### We can formulate projection optimization problem to find these vectors

Since we assume that our initial matrix $\tilde{V}^T$ contains only $K$ main components hidden inside. We can try to represent this matrix in terms of these components.

First, we will define a set of $K$ $N$-dimensional row vectors, which will define new axes of our projected space.

$$R = \begin{bmatrix} r_{1,1} & r_{1,2} & \cdots & r_{1,N} \\ r_{2,1} & r_{2,2} & \cdots & r_{2,N} \\ \vdots & \vdots & \ddots & \vdots \\ r_{K,1} & r_{K,2} & \cdots & r_{K,N} \end{bmatrix} \begin{matrix} \rightarrow axis\ 1 \\ \rightarrow axis\ 2 \\ \vdots \\ \rightarrow axis\ K \end{matrix}$$

Where rows of $R$ are orthogonal

$$\mathrm{R} \times R^T = \mathbb{I}_{K \times K}$$

$R^T \times R$ is then a projection operator to this new reduced space spawned by these vectors.

Having this we will try to find this projection matrix $R$ (of $K$ row vectors) so that projected expression matrix will contain the maximum possible signal from the original matrix.

$$\boxed{\min_{R} \left\| \tilde{V}^T - R^T \times R \times \tilde{V}^T \right\|_{s.t.\quad R \times R^T = \mathbb{I}_{K \times K}}} \tag{1}$$

Note, that this projection minimization problem could be solved using SVD (Yuan & Oja, 2005). Vectors $R^T$ could be found as left singular vectors of original matrix $\tilde{V}^T$, which are length normalized eigenvectors of $\tilde{V}^T \times \tilde{V}$.

### We will reformulate the NMF/deconvolution optimization problem in a projected space.

We can reformulate this problem in terms of $\widetilde{H}$ and $\widetilde{W}$

$$\min_{R} \left\| \tilde{V}^T - R^T \times R \times \widetilde{H}^T \times \widetilde{W}^T \right\|$$

Once $R$ matrix is defined, we can project our initial basis column vectors $\widetilde{H}^T$on a new reduced space. Let's define new matrix $X$ in the following way:

$$\widetilde{H} = X \times R$$

Or in transposed form

$$\widetilde{H}^T = R^T \times X^T$$

Where columns of $X^T$ are coordinates of original column vectors $\widetilde{H}^T$ in a $K$-dimensional space spawned by column vectors $R^T$.

Note that since projection is a linear operation and coordinates of vectors $\left(\tilde{h}^T\right)_{*,k}$ in initial space were defining corners of the simplex, then coordinates of projected vectors $(\tilde{x}^T)_{*,k}$ will also be defining corners of the simplex in projected space.

We can also find equation for $X$

$$\widetilde{H}^T = R^T \times X^T \Leftrightarrow R \times \widetilde{H}^T = \underbrace{R \times R^T}_{\mathbb{I}_{K \times K}} \times X^T$$

Which means that

$$X^T = R \times \widetilde{H}^T \xrightarrow{transpose} X = \widetilde{H} \times R^T$$

Where $X$ is a $K \times K$ dimensional matrix which rows contain coordinates of the original vectors in a new space.

$$X = \begin{bmatrix} x_{1,1} & x_{1,2} & \cdots & x_{1,K} \\ x_{2,1} & x_{2,2} & \cdots & x_{2,K} \\ \vdots & \vdots & \ddots & \vdots \\ x_{K,1} & x_{K,2} & \cdots & x_{K,K} \end{bmatrix} \begin{matrix} 1 \\ 2 \\ \vdots \\ K \end{matrix}$$

Once we found projection vectors $R$ (1) and coordinates $X$ we can now solve our NMF problem in projected space.

Using $X$ we can rewrite initial optimization problem:

$$\min_{R} \left\| \tilde{V}^T - R^T \times R \times \tilde{V}^T \right\| \Rightarrow \min_{\substack{R \\ \widetilde{W}}} \left\| \tilde{V}^T - R^T \times \underbrace{R \times \boldsymbol{\widetilde{H}^T}}_{X^T} \times \boldsymbol{\widetilde{W}^T} \right\| \Rightarrow \min_{\substack{X \\ \widetilde{W}}} \left\| \tilde{V}^T - R^T \times \boldsymbol{X^T} \times \widetilde{W}^T \right\|$$

Therefore, the optimization problem in a space of $X$ and $W$ could be formulated:

$$\boxed{\min_{\substack{X \\ \widetilde{W}}} \left\| \tilde{V}^T - R^T \times X^T \times \widetilde{W}^T \right\| \quad s.t. \quad \begin{matrix} \widetilde{W} > 0 \\ \widetilde{H} = X \times R > 0 \end{matrix}}$$

Which could be reformulated in projected space:

$$\min_{\substack{X \\ \widetilde{W}}} \left\| R \times \tilde{V}^T - R \times R^T \times X^T \times \widetilde{W}^T \right\| \Rightarrow \min_{\substack{X \\ \widetilde{W}}} \left\| R \times \tilde{V}^T - X^T \times \widetilde{W}^T \right\| \Rightarrow \min_{\substack{X \\ \widetilde{W}}} \left\| \tilde{V} \times R^T - \widetilde{W} \times X \right\|$$

Which leads us to

$$\boxed{\min_{\substack{X \\ \widetilde{W}}} \left\| \tilde{V} \times R^T - \widetilde{W} \times X \right\| \quad s.t. \quad \begin{matrix} \widetilde{W} > 0 \\ \widetilde{H} = X \times R > 0 \end{matrix}}$$

### <u>Summary</u>

So, we formulated the NMF optimization for normalized matrices as a problem of optimization in some projected space where we first will find projection operator vectors $R$ and then will solve the NMF/deconvolution problem in a projected space where instead of optimizing in $K \times N$ matrix $\widetilde{H}$ we are optimizing $K \times K$ matrix $X$.

1. *Find projection vectors R for $\tilde{V}$ so that*

$$\min_{R} \left\| \tilde{V}^T - R^T \times R \times \tilde{V}^T \right\|_{s.t. \quad R \times R^T = \mathbb{I}_{K \times K}}$$

Where $R^T$ could be found as left singular vectors of $\tilde{V}^T$ from SVD.

2. *Solve the NMF/deconvolution problem for this projected space*

$$\min_{\substack{X \\ \widetilde{W}}} \left\| \tilde{V} \times R^T - \widetilde{W} \times X \right\|_{\substack{s.t. \quad \widetilde{W} > 0 \\ \widetilde{H} = X \times R > 0}}$$

Where $X$ defines corners of the simplex in projected space

$$\widetilde{H} = X \times R$$

$$X^T = R \times \widetilde{H}^T$$

### Supplementary Note 5. Projective formulation of the NMF problem in a features' space (Vectors $S$)

#### What we did so far

*Defined the NMF problem:*

$$V = W \times H$$

$$v_{i,j} \in \mathbb{R}^{M \times N}, \qquad w_{i.j} \in \mathbb{R}^{M \times K}, \qquad h_{i,j} \in \mathbb{R}^{K \times N}$$

$$\begin{aligned} &K - number\ of\ pure\ components \\ &N - number\ of\ samples \\ &M - number\ of\ features \end{aligned}$$

Where $v_{i,j}$ describes value of $i^{th}$ feature in $j^{th}$ sample, $w_{i.k}$ describes value of $i^{th}$ feature (e.g. gene expression) in $k^{th}$ pure component (e.g. pure cell type), $h_{k,j} \geq 0$ describes contribution value (e.g. proportion) of $k^{th}$ pure components (e.g. pure cell types),in $j^{th}$ sample.

*Defined the NMF problem for normalized matrices $\tilde{\boldsymbol{V}}$ in a samples space and $\tilde{\tilde{\boldsymbol{V}}}$ in a features space:*

What we did so far is we defined new NMF equations for normalized matrices.

$$V = W \times H \Rightarrow \tilde{V} = \widetilde{W} \times \widetilde{H} \Rightarrow \tilde{V}^T = \widetilde{H}^T \times \widetilde{W}^T \Rightarrow \tilde{\tilde{\boldsymbol{V}}} = \widetilde{\widetilde{\boldsymbol{W}}} \times \tilde{\tilde{\boldsymbol{H}}}$$

$$\begin{aligned} &H - column\ normalized\ matrix \\ &\tilde{V}, \widetilde{W}, \widetilde{H} - row\ normalized\ matrices \\ &\tilde{V}^T, \widetilde{H}^T, \widetilde{W}^T - column\ normalized\ matrices \\ &\widetilde{\widetilde{W}}, \tilde{\tilde{V}}, \tilde{\tilde{H}} - row\ normalized\ matrices \end{aligned}$$

#### We formulate the optimization problem to find these new NMF matrices

In terms of optimization, our main goal is to find such matrices $\tilde{\tilde{H}}$ and $\widetilde{\widetilde{W}}$ so that combination of these matrices will give us initial data matrix with highest possible accuracy (i.e. lowest error).

$$\min_{\tilde{\tilde{H}}, \widetilde{\widetilde{W}}} \left\| \tilde{\tilde{V}} - \widetilde{\widetilde{W}} \times \tilde{\tilde{H}} \right\| \quad s.t. \quad \tilde{\tilde{H}} > 0, \ \widetilde{\widetilde{W}} > 0$$

Note that this formulation is equivalent to deconvolution problem, since $\tilde{\tilde{H}}$ is column normalized.

#### Note: $\tilde{\tilde{\boldsymbol{V}}}$ is projectable onto $\boldsymbol{K}$ dimensional space.

Similar to what we described for $\tilde{V}^T$, we assume that matrix $\tilde{\tilde{V}}$ contains $K$ main components hidden inside. And we can try to represent this matrix in terms of these components.

### We can formulate projection optimization problem to find these vectors

First, we will define a set of $K$ $M$-dimensional row vectors, which will define new axes of our projected space.

$$S = \begin{bmatrix} s_{1,1} & s_{1,2} & \cdots & s_{1,M} \\ s_{2,1} & s_{2,2} & \cdots & s_{2,M} \\ \vdots & \vdots & \ddots & \vdots \\ s_{K,1} & s_{K,2} & \cdots & s_{K,M} \end{bmatrix} \begin{matrix} \rightarrow axis\ 1 \\ \rightarrow axis\ 2 \\ \vdots \\ \rightarrow axis\ K \end{matrix}$$

Where rows of $S$ are orthogonal

$$\mathrm{S} \times S^T = \mathbb{I}_{K\times K}$$

$S^T \times S$ is then a projection operator to this new reduced space spawned by these vectors.

Having this we will try to find this projection matrix $S$ (of $K$ row vectors) so that projected data matrix will contain the maximum possible signal from the original matrix.

$$\boxed{\min_{S} \left\| \tilde{\tilde{V}} - S^T \times S \times \tilde{\tilde{V}} \right\|_{s.t.\quad S\times S^T = \mathbb{I}_{K\times K}}} \qquad (2)$$

Note, that this projection minimization problem could be solved using SVD (Yuan & Oja, 2005). Vectors $S^T$ could be found as left singular vectors of original matrix $\tilde{\tilde{V}}$, which are length normalized eigenvectors of $\tilde{\tilde{V}} \times \tilde{\tilde{V}}^T$.

### We will reformulate the NMF/deconvolution optimization in a projected space.

We can reformulate this problem in terms of $\tilde{\tilde{W}}$ and $\tilde{\tilde{H}}$

$$\min_{S} \left\| \tilde{\tilde{V}} - S^T \times S \times \tilde{\tilde{W}} \times \tilde{\tilde{H}} \right\|$$

Once $S$ matrix is defined it then could be used to project our initial basis column vectors $\tilde{\tilde{W}}$ on a new reduced space. Let's define new matrix $\Omega$ in the following way:

$$\tilde{\tilde{W}} = S^T \times \Omega$$

Where columns of $\Omega$ are coordinates of original column vectors $\tilde{\tilde{W}}$ in a $K$-dimensional space spawned by column vectors $S^T$.

Note that since projection is a linear operation and coordinates of vectors $\left(\tilde{\tilde{w}}\right)_{*,k}$ in initial space were defining corners of the simplex, then coordinates of projected vectors $(\omega)_{*,k}$ will also be defining corners of the simplex in projected space.

We can also find equation for $\Omega$

$$\tilde{\tilde{W}} = S^T \times \Omega \Leftrightarrow S \times \tilde{\tilde{W}} = \underbrace{S \times S^T}_{\mathbb{I}_{K\times K}} \times \Omega$$

Which means that

$$\Omega = S \times \tilde{\tilde{W}}$$

Where $\Omega$ is a $K \times K$ dimensional matrix which columns contains coordinates of the original vectors in a new space.

$$\Omega = \begin{bmatrix} \omega_{1,1} & \omega_{1,2} & \cdots & \omega_{1,K} \\ \omega_{2,1} & \omega_{2,2} & \cdots & \omega_{2,K} \\ \vdots & \vdots & \ddots & \vdots \\ \omega_{K,1} & \omega_{K,2} & \cdots & \omega_{K,K} \end{bmatrix}$$
$$\quad 1 \qquad 2 \quad \ldots \quad K$$

Once we found projection vectors $S$ (2) and coordinates $\Omega$ we can now solve our NMF/deconvolution problem in projected space.

Using $\Omega$, the optimization problem could be rewritten as:

$$\min_{S} \left\| \tilde{\tilde{V}} - S^T \times S \times \tilde{\tilde{V}} \right\| \Rightarrow \min_{\substack{S \\ \tilde{\tilde{H}}}} \left\| \tilde{\tilde{V}} - S^T \times \underbrace{S \times \tilde{\tilde{W}}}_{\Omega} \times \tilde{\tilde{H}} \right\| \Rightarrow \min_{\substack{\Omega \\ \tilde{\tilde{H}}}} \left\| \tilde{\tilde{V}} - S^T \times \Omega \times \tilde{\tilde{H}} \right\|$$

Therefore, the optimization problem in a space of $\Omega$ and $W$ could be formulated as

$$\boxed{\min_{\substack{\Omega \\ \tilde{\tilde{H}}}} \left\| \tilde{\tilde{V}} - S^T \times \Omega \times \tilde{\tilde{H}} \right\| \quad s.t. \quad \tilde{\tilde{H}} > 0, \ \tilde{\tilde{W}} = S^T \times \Omega > 0}$$

Which could be reformulated to projected space:

$$\min_{\substack{\Omega \\ \tilde{\tilde{H}}}} \left\| S \times \tilde{\tilde{V}} - S \times S^T \times \Omega \times \tilde{\tilde{H}} \right\| \Rightarrow \min_{\substack{\Omega \\ \tilde{\tilde{H}}}} \left\| S \times \tilde{\tilde{V}} - \Omega \times \tilde{\tilde{H}} \right\|$$

Which leads us to

$$\min_{\substack{\Omega \\ \tilde{\tilde{H}}}} \left\| S \times \tilde{\tilde{V}} - \Omega \times \tilde{\tilde{H}} \right\| \quad s.t. \quad \tilde{\tilde{H}} > 0, \ \tilde{\tilde{W}} = S^T \times \Omega > 0$$

### **Summary**

So, we reformulated our optimization problem as a problem of optimization in some projected space where we first find projection operator vectors $S$ and then will solve NMF/deconvolution problem in a projected space where instead of optimizing in $M \times K$ matrix we are optimizing $K \times K$ matrix $\Omega$.

*1. Find projection vectors S for $\tilde{\tilde{V}}$ so that*

$$\min_{S} \left\| \tilde{\tilde{V}} - S^T \times S \times \tilde{\tilde{V}} \right\|_{s.t. \quad S \times S^T = \mathbb{I}_{K \times K}}$$

Where $S^T$ could be found as left singular vectors of $\tilde{\tilde{V}}$ from SVD.

*2. Solve the NMF/deconvolution problem for this projected space*

$$\min_{\substack{\Omega \\ \tilde{\tilde{H}}}} \left\| S \times \tilde{\tilde{V}} - \Omega \times \tilde{\tilde{H}} \right\|_{\substack{s.t. \quad \tilde{\tilde{H}} > 0 \\ \tilde{\tilde{W}} = S^T \times \Omega > 0}}$$

Where Ω defines corners of the simplex in projected space

$$\tilde{\tilde{W}} = S^T \times \Omega$$
$$\Omega = S \times \tilde{\tilde{W}}$$

### Supplementary Note 6. Additional sum-to-one constraint for projection vectors $\boldsymbol{R}$

#### What we did so far

*Defined the NMF/deconvolution problem for matrix $\tilde{\boldsymbol{V}}$ in a samples space:*

$$V = W \times H \Rightarrow \tilde{V} = \widetilde{W} \times \widetilde{H} \Rightarrow \tilde{\mathbf{V}}^{\boldsymbol{T}} = \widetilde{\boldsymbol{H}}^{\boldsymbol{T}} \times \widetilde{\boldsymbol{W}}^{\boldsymbol{T}}$$

$$v_{i,j} \in \mathbb{R}^{M \times N}, \qquad w_{i.j} \in \mathbb{R}^{M \times K}, \qquad h_{i,j} \in \mathbb{R}^{K \times N}$$

$$K - number\ of\ pure\ components$$
$$N - number\ of\ samples$$
$$M - number\ of\ features$$

$$H - column\ normalized\ matrix$$
$$\tilde{V}, \widetilde{W}, \widetilde{H} - row\ normalized\ matrices$$
$$\tilde{\mathrm{V}}^T, \widetilde{H}^T, \widetilde{W}^T - column\ normalized\ matrices$$

*Decomposed the NMF/deconvolution problem into two steps in order to solve this task in projected space*

*1.Find projection vectors $R$ for $\tilde{V}$ so that*

$$\min_{R} \left\| \tilde{V}^T - R^T \times R \times \tilde{V}^T \right\|_{s.t.\quad R \times R^T = \mathbb{I}_{K \times K}}$$

Where $R$ could be found as left singular vectors of $\tilde{V}^T$ from SVD.

*2. Solve NMF/deconvolution problem for this projected space*

$$\min_{\substack{X \\ \widetilde{W}}} \left\| \tilde{V} \times R^T - \widetilde{W} \times X \right\|_{\substack{s.t.\quad \widetilde{W} > 0 \\ \widetilde{H} = X \times R > 0}}$$

Where $X$ defines corners of the simplex in projected space

$$\widetilde{H} = X \times R$$

$$X^T = R \times \widetilde{H}^T$$

#### Intuition for moving into $\boldsymbol{K} - \mathbf{1}$ dimensional space.

We already showed that matrix $\tilde{V}^T$ forms a simplex and this simplex is practically $(K-1)$ – dimensional due to sum-to-one normalization.

We have represented each feature vector $(\tilde{v}^T)_{*,i}$ as a combination of $K$ coefficients vectors for each pure type with sample dependent coefficients. But coefficients are not arbitrary. Since these coefficients sum to 1, the last coefficient could be derived from this sum.

$$(\tilde{v}^T)_{*,i} = (\tilde{v}_{i,*})^T = \begin{bmatrix} \sum_k \alpha_{i,k} * \tilde{h}_{k,1} \\ \vdots \\ \sum_k \alpha_{i,k} * \tilde{h}_{k,N} \end{bmatrix} = \alpha_{i,1} \begin{bmatrix} \tilde{h}_{1,1} \\ \vdots \\ \tilde{h}_{1,N} \end{bmatrix} + \alpha_{i,2} \begin{bmatrix} \tilde{h}_{2,1} \\ \vdots \\ \tilde{h}_{2,N} \end{bmatrix} + \cdots + \alpha_{i,K} \begin{bmatrix} \tilde{h}_{K,1} \\ \vdots \\ \tilde{h}_{K,N} \end{bmatrix}$$

$$\sum\nolimits_{k=1}^{K}\alpha_{i,k}=1\Leftrightarrow\alpha_{i,K}=1-\sum\nolimits_{k=1}^{K-1}\alpha_{i,k}$$

Intuitively this means that we can get rid of 1 dimension in our task, since this dimension is predefined by others. Therefore, the whole structure of the geometrical set of points is $(K-1)$-dimensional. Next, we will show how this transition in dimensions from $K$ to $(K-1)$ could be considered in a defined projection operator.

### Transition to a (K-1) dimensional formulation via additional orthogonality constraint.

Geometrically speaking we can get $(K-1)$-dimensional representation of our data by centering the whole dataset.

SVD vectors $R$ we got as a solution for the projection minimization problem do not consider this property. $(K-1)$-dimensional structure of a simplex (e.g. $2D$-triangle) is still described by these $K$ vectors, but the whole structure is shifted from zero and rotated (e.g. 3 vectors describe 2D triangle rotated in some way). The natural way to describe such $(K-1)$ dimensional structure is to consider orthogonal vector $R_1$ which will only represent the shift of the whole $(K-1)$ structure away from the zero point, but not its structure.

For general SVD vectors this property is not met, since first vector is usually just pointing to a mean value of the dataset (Kim & You, 2023).

$$\vec{R}_c=\frac{\sum_{j=1}^{N}\tilde{V}_{*,i}^{T}}{M}$$

This vector is not mandatory orthogonal to a simplex hyperplane (Fig. S2).

However, once the data is centered, the shift for all points is eliminated, the mean value is now zero and all $(K-1)$ vectors represent the entire simplex structure and the hyperplane, since no other variability left(Fig. S2).

$$\tilde{V_c}^{T}=\tilde{V}^{T}-\frac{\sum_{i=1}^{M}\tilde{V}_{*,i}^{T}}{M}\mathbb{I}_M \quad R''=\min_{R}\left\|\tilde{V}_c^{T}-R^{T}\times R\times\tilde{V}_c^{T}\right\|_{s.t.\quad R\times R^{T}=\mathbb{I}_{(K-1)\times(K-1)}}$$

Vectors $R''$ are in the same hyperplane where the points are, but this system describes points only in a space of centered matrix. We can represent initial points in original space using vectors $(\vec{R}_c,\vec{R}_1'',\dots,\vec{R}_{K-1}'')$ but these vectors are not orthogonal. One of the possible ways to make this system orthogonal is to perform orthogonalization procedure.

In this work we added orthogonality constraint directly to projection minimization problem.

### Sum-to-one property leads to constraint for vector $R_1$

Geometrically speaking orthogonality of the first vector to the simplex hyperplane means that scalar projection of each of the column vectors, forming $\tilde{V}^{T}$ onto vector $\vec{R}_1$ is a constant.

$$\forall j\in[1,M]\ \left|proj_{\vec{R}_1}\left(\tilde{V}^{T}\right)_{*,j}\right|=\begin{bmatrix}\tilde{V}_{1,j}^{T} & \dots & \tilde{V}_{N,j}^{T}\end{bmatrix}\begin{bmatrix}r_{1,1}\\ \vdots\\ r_{1,N}\end{bmatrix}=\begin{bmatrix}\tilde{V}_{j,1} & \dots & \tilde{V}_{j,N}\end{bmatrix}\begin{bmatrix}r_{1,1}\\ \vdots\\ r_{1,N}\end{bmatrix}=c$$

In a matrix form this could be represented for the entire matrix $\tilde{V}$

$$\boxed{\tilde{V}\vec{R}_1 = \begin{bmatrix} c \\ \vdots \\ c \end{bmatrix} = c\mathbb{I}_M}$$

This constraint could be directly included into projection optimization formulation to define optimization problem in space of the original matrix, which is not centered.

$$R^* = \min_R \left\| \tilde{V}^T - R^T \times R \times \tilde{V}^T \right\|_{\substack{s.t. \quad R \times R^T = \mathbb{I}_{K \times K} \\ \tilde{V}\vec{R}_1 = c\mathbb{I}_M}}$$

Importantly solution vectors $\vec{R}_1^*, \dots, \vec{R}_K^*$ are now will be orthogonal and vectors $\vec{R}_2^*, \dots, \vec{R}_K^*$ will describe the entire $(K-1)$-dimensiona hyperplane, where simplex is located (Fig S2).

### Summary

All above means that deconvolution problem is now formulated through the projection operator which has specific property we want to enforce.

1. *Find projection vectors $R$ for $\tilde{V}^T$ so that*

$$\min_R \left\| \tilde{V}^T - R^T \times R \times \tilde{V}^T \right\|_{\substack{s.t. \quad R \times R^T = \mathbb{I}_{K \times K} \\ \tilde{V}\vec{R}_1 = c\mathbb{I}_M}}$$

Where $R^T$ now is different from singular vectors of $\tilde{V}^T$.However $\vec{R}_2, \dots, \vec{R}_K$ can be obtained as left singular vectors of the centered matrix $\tilde{V}_c^T$

2. *Solve deconvolution problem for this projected space*

$$\min_{\substack{X \\ \widetilde{W}}} \left\| \tilde{V} \times R^T - \widetilde{W} \times X \right\|_{\substack{s.t. \quad \widetilde{W} > 0 \\ \widetilde{H} = X \times R > 0}}$$

Where $X$ defines corners of the simplex in projected space

$$\widetilde{H} = X \times R$$
$$X^T = R \times \widetilde{H}^T$$

### Supplementary Note 7. Additional sum-to-one constraint for projection vectors $S$

#### What we did so far

*Defined the NMF/deconvolution problem for normalized matrix $\tilde{\tilde{\boldsymbol{V}}}$ in a features' space:*

$$V = W \times H \Rightarrow \tilde{V} = \widetilde{W} \times \widetilde{H} \Rightarrow \widetilde{V}^T = \widetilde{H}^T \times \widetilde{W}^T \Rightarrow \tilde{\tilde{\boldsymbol{V}}} = \widetilde{\widetilde{\boldsymbol{W}}} \times \tilde{\tilde{\boldsymbol{H}}}$$

$$v_{i,j} \in \mathbb{R}^{M\times N}, \qquad w_{i.j} \in \mathbb{R}^{M\times K}, \qquad h_{i,j} \in \mathbb{R}^{K\times N}$$

$$K - number\ of\ pure\ components$$
$$N - number\ of\ samples$$
$$M - number\ of\ features$$

$$H - column\ normalized\ matrix$$
$$\tilde{V}, \widetilde{W}, \widetilde{H} - row\ normalized\ matrices$$
$$\widetilde{V}^T, \widetilde{H}^T, \widetilde{W}^T - column\ normalized\ matrices$$
$$\widetilde{\widetilde{W}}, \tilde{\tilde{V}}, \tilde{\tilde{H}} - row\ normalized\ matrices$$

*Decomposed the NMF/deconvolution problem into two problems to solve this task in projected space*

*1.Find projection vectors $S$ for $\tilde{\tilde{V}}$ so that*

$$\min_{S} \left\| \tilde{\tilde{V}} - S^T \times S \times \tilde{\tilde{V}} \right\|_{s.t.\quad S\times S^T = \mathbb{I}_{K\times K}}$$

Where $\Omega$ could be found as left singular vectors of $\tilde{\tilde{V}}$

*2. Solve deconvolution problem for this projected space*

$$\min_{\substack{X \\ \tilde{\tilde{H}}}} \left\| S \times \tilde{\tilde{V}} - \Omega \times \tilde{\tilde{H}} \right\|_{\substack{s.t.\quad \tilde{\tilde{H}} > 0 \\ \widetilde{\widetilde{W}} = S^T \times \Omega > 0}}$$

Where $\Omega$ defines corners of the simplex in projected space

$$\widetilde{\widetilde{W}} = S^T \times \Omega$$

$$\Omega = S \times \widetilde{\widetilde{W}}$$

#### Intuition for moving into $K - 1$ dimensional space.

Note, that again we showed that matrix $\tilde{\tilde{V}}$ forms a simplex and this simplex is practically $K - 1$ dimensional.

We have represented our sample vector $\tilde{\tilde{v}}_{*,j}$ as a combination of $K$ pure component vectors with coefficients depending on contribution values. And since these coefficients sum to 1, the last coefficient could be derived from this sum.

$$\tilde{\tilde{v}}_{*,j} = \begin{bmatrix} \sum_k \tilde{\tilde{w}}_{1,k} * \alpha_{k,j} \\ \vdots \\ \sum_k \tilde{\tilde{w}}_{M,k} * \alpha_{k,j} \end{bmatrix} = \alpha_{1,j} \begin{bmatrix} \tilde{\tilde{w}}_{1,1} \\ \vdots \\ \tilde{\tilde{w}}_{M,1} \end{bmatrix} + \alpha_{2,j} \begin{bmatrix} \tilde{\tilde{w}}_{1,2} \\ \vdots \\ \tilde{\tilde{w}}_{M,2} \end{bmatrix} + \cdots + \alpha_{K,j} \begin{bmatrix} \tilde{\tilde{w}}_{1,K} \\ \vdots \\ \tilde{\tilde{w}}_{M,K} \end{bmatrix}$$

$$\sum_{k=1}^{K} \alpha_{k,j} = 1 \Leftrightarrow \alpha_{K,j} = 1 - \sum_{k=1}^{K-1} \alpha_{k,j}$$

Intuitively this means that we can get rid of 1 dimension in our task, since this dimension is predefined by others.

Next, we will show how this transition in dimensions from $K$ to $(K-1)$ could be considered in a defined projection operator.

### Transition to a $(K-1)$- dimensional formulation via additional orthogonality constraint

Similarly for original problem formulation, the first vector is pointing to the mean of the whole dataset.

$$\vec{S}_c = \frac{\sum_{j=1}^{N} \tilde{\tilde{V}}_{*,j}}{N}$$

And it is not mandatory orthogonal to the simplex hyperplane.

However, once the data is centered, the shift for all points is eliminated, the mean value is now zero and all $(K-1)$ vectors represent the entire simplex structure and the hyperplane, since no other variability left (Fig. S2).

$$\tilde{\tilde{V}}_c = \tilde{\tilde{V}} - \frac{\sum_{j=1}^{N} \tilde{\tilde{V}}_{*,j}}{N} \mathbb{I}_N \quad S'' = \min_{S} \left\| \tilde{\tilde{V}}_c - S^T \times S \times \tilde{\tilde{V}}_c \right\| \quad s.t. \quad S \times S^T = \mathbb{I}_{(K-1)\times(K-1)}$$

Similarly, we can include the search of these vectors into original problem by considering orthogonality of the first vector $\vec{S}_1$ to the hyperplane.

In this work we added orthogonality constraint directly to projection minimization problem.

### Sum-to-one property leads to constraint for vector $S_1$

Geometrically speaking orthogonality of the first vector to the simplex hyperplane to means that scalar projection of each of the column vectors, forming $\tilde{\tilde{V}}$ onto vector $\vec{S}_1$ is a constant.

$$\forall\, i \in [1, N] \left| proj_{\vec{S}_1} \left( \tilde{\tilde{V}} \right)_{*,i} \right| = \begin{bmatrix} \tilde{\tilde{V}}_{1,i} & \dots & \tilde{\tilde{V}}_{N,i} \end{bmatrix} \begin{bmatrix} s_{1,1} \\ \vdots \\ s_{1,N} \end{bmatrix} = c$$

In a matrix form this could be represented for the entire matrix $\tilde{V}$

$$\boxed{\tilde{\tilde{V}}^T \vec{S}_1 = \begin{bmatrix} c \\ \vdots \\ c \end{bmatrix} = c\mathbb{I}_N}$$

This constraint could be directly included into projection optimization formulation to define optimization problem in space of the original matrix, which is not centered.

$$S^* = \min_{S} \left\| \tilde{\tilde{V}} - S^T \times S \times \tilde{\tilde{V}} \right\| \quad \underset{\substack{\tilde{\tilde{V}}^T \vec{S}_1 = c\mathbb{I}_N}}{s.t.\quad S \times S^T = \mathbb{I}_{K \times K}}$$

Importantly solution vectors $\vec{S}_1^*, \ldots, \vec{S}_K^*$ are now will be orthogonal and vectors $\vec{S}_2^*, \ldots, \vec{S}_K^*$ will describe the entire $(K-1)$-dimensiona hyperplane, where simplex is located (Fig S2).

### Summary

All above means that deconvolution problem is now formulated as

1. *Find projection vectors S for $\tilde{\tilde{V}}$ so that*

$$\min_{S} \left\| \tilde{\tilde{V}} - S^T \times S \times \tilde{\tilde{V}} \right\| \quad \underset{\substack{\tilde{\tilde{V}}^T \vec{S}_1 = c\mathbb{I}_N}}{s.t.\quad S \times S^T = \mathbb{I}_{K \times K}}$$

Where $S^T$ now is different from singular vectors of $\tilde{\tilde{V}}$. However, $\vec{S}_2, \ldots, \vec{S}_K$ could be found as left singular vectors of the centered matrix $\tilde{\tilde{V}}_c$.

2. *Solve deconvolution problem for this projected space*

$$\min_{\substack{\Omega \\ \tilde{\tilde{H}}}} \left\| S \times \tilde{\tilde{V}} - \Omega \times \tilde{\tilde{H}} \right\| \quad \underset{\substack{\tilde{\tilde{W}} = S^T \times \Omega > 0}}{s.t.\quad \tilde{\tilde{H}} > 0}$$

Where Ω defines corners of the simplex in projected space

$$\tilde{\tilde{W}} = S^T \times \Omega$$
$$\Omega = S \times \tilde{\tilde{W}}$$

### Supplementary Note 8. Sinkhorn transformation links two optimization problems together

#### What we did so far

*Defined the NMF/deconvolution problem for normalized matrices $\tilde{\boldsymbol{V}}$ in a samples' space and $\tilde{\tilde{\boldsymbol{V}}}$ in a features' space:*

$$V = W \times H \Rightarrow \tilde{V} = \tilde{W} \times \tilde{H} \Rightarrow \tilde{\mathbf{V}}^{\boldsymbol{T}} = \tilde{\boldsymbol{H}}^{\boldsymbol{T}} \times \tilde{\boldsymbol{W}}^{\boldsymbol{T}} \Rightarrow \tilde{\tilde{\boldsymbol{V}}} = \tilde{\tilde{\boldsymbol{W}}} \times \tilde{\tilde{\boldsymbol{H}}}$$

$$v_{i,j} \in \mathbb{R}^{M \times N}, \qquad w_{i.j} \in \mathbb{R}^{M \times K}, \qquad h_{i,j} \in \mathbb{R}^{K \times N}$$

$$K - number\ of\ pure\ components$$
$$N - number\ of\ samples$$
$$M - number\ of\ features$$

$$H - column\ normalized\ matrix$$
$$\tilde{V}, \tilde{W}, \tilde{H} - row\ normalized\ matrices$$
$$\tilde{\mathrm{V}}^T, \tilde{H}^T, \tilde{W}^T - column\ normalized\ matrices$$
$$\tilde{\tilde{W}}, \tilde{\tilde{V}}, \tilde{\tilde{H}} \ - row\ normalized\ matrices$$

The optimization goal for first matrix:

$$\min_{\tilde{H}^T, \tilde{W}^T} \left\| \tilde{V}^T - \tilde{H}^T \times \tilde{W}^T \right\|_{\substack{s.t. \quad \tilde{H}^T > 0 \\ \tilde{W}^T > 0}}$$

And for second matrix:

$$\min_{\tilde{\tilde{H}}, \tilde{\tilde{W}}} \left\| \tilde{\tilde{V}} - \tilde{\tilde{W}} \times \tilde{\tilde{H}} \right\|_{\substack{s.t. \quad \tilde{\tilde{H}} > 0 \\ \tilde{\tilde{W}} > 0}}$$

We also showed that points of these to matrices lie on two simplexes.

*Decomposed each optimization problem into two steps*

Each optimization problem was approached through the projection optimization problem with additional constraints.

For the first matrix:

1. *Find projection vectors $R$ for $\tilde{V}^T$ so that*

$$\min_{R}\left\|\tilde{V}^T - R^T \times R \times \tilde{V}^T\right\|_{\substack{s.t.\quad R\times R^T=\mathbb{I}_{K\times K}\\ \tilde{V}\vec{R}_1=c\mathbb{I}_M}}$$

Where $R^T$ now is different from singular vectors of $\tilde{V}^T$.However $\vec{R}_2, \dots, \vec{R}_K$ can be obtained as left singular vectors of the centered matrix $\tilde{V}_c^T$

2. *Solve deconvolution problem for this projected space*

$$\min_{\substack{X\\ \widetilde{W}}}\left\|\tilde{V} \times R^T - \widetilde{W} \times X\right\|_{\substack{s.t.\quad \widetilde{W}>0\\ \widetilde{H}=X\times R>0}}$$

Where $X$ defines corners of the simplex in projected space

$$\widetilde{H} = X \times R$$
$$X^T = R \times \widetilde{H}^T$$

For the second matrix:

1. *Find projection vectors S for $\tilde{\tilde{V}}$ so that*

$$\min_{S}\left\|\tilde{\tilde{V}} - S^T \times S \times \tilde{\tilde{V}}\right\|_{\substack{s.t.\quad S\times S^T=\mathbb{I}_{K\times K}\\ \tilde{\tilde{V}}^T\vec{S}_1=c\mathbb{I}_N}}$$

Where $S^T$ now is different from singular vectors of $\tilde{\tilde{V}}$. However, $\vec{S}_2, \dots, \vec{S}_K$ could be found as left singular vectors of the centered matrix $\tilde{\tilde{V}}_c$.

2. *Solve deconvolution problem for this projected space*

$$\min_{\substack{\Omega\\ \tilde{\tilde{H}}}}\left\|S \times \tilde{\tilde{V}} - \Omega \times \tilde{\tilde{H}}\right\|_{\substack{s.t.\quad \tilde{\tilde{H}}>0\\ \tilde{\tilde{W}}=S^T\times\Omega>0}}$$

Where $\Omega$ defines corners of the simplex in projected space

$$\tilde{\tilde{W}} = S^T \times \Omega$$
$$\Omega = S \times \tilde{\tilde{W}}$$

### *Introduced iterative Sinkhorn transformation for positive data matrices*

$$V \overset{\substack{row\\norm}}{\Longrightarrow} \underbrace{D_{v_0} \times V}_{\substack{\tilde{V}\\ simplex\ in\\ samples\ space}} \overset{\substack{column\\norm}}{\Longrightarrow} \underbrace{D_{v_0} \times V \times D_{v_1}}_{\substack{\tilde{\tilde{V}}\\ simplex\ in\\ features\ space}} \overset{\substack{row\\norm}}{\Longrightarrow} \dots \overset{\substack{row\\norm}}{\Longrightarrow} \underset{\substack{simplex\ in\\ samples\ space}}{\overset{\infty}{V}_{ss}} \overset{\substack{column\\norm}}{\Longrightarrow} \underset{\substack{simplex\ in\\ features\ space}}{\overset{\infty}{V}_{fs}}$$

From which we got the constant relation between two simplexes defined.

$$\overset{\infty}{V}_{ss} \times \frac{N}{M} = \overset{\infty}{V}_{fs}$$

**Projection minimization and deconvolution problems could be formulated for Sinkhorn-transformed matrices**

*For simplex of features in a samples space*

*1. Find projection vectors $R$ for $\overset{\infty}{V}_{ss}^{T}$ so that*

$$\min_{R} \left\| \overset{\infty}{V}_{ss}^{T} - R^{T} \times R \times \overset{\infty}{V}_{ss}^{T} \right\| \quad \underset{\overset{\infty}{V}_{ss}\vec{R}_1 = c\mathbb{I}_M}{s.t.\ \ R\times R^{T} = \mathbb{I}_{K\times K}}$$

Where: $\vec{R}_2, \ldots, \vec{R}_K$ could be found as left singular vectors of the centered matrix ${\overset{\infty}{V}_{ss}^{T}}_{c}$.

*2. Solve deconvolution problem for this projected space*

$$\min_{\overset{\infty}{W}_{ss},\, X} \left\| \overset{\infty}{V}_{ss} \times R^{T} - \overset{\infty}{W}_{ss} \times X \right\| \quad s.t.\ \ \overset{\infty}{W}_{ss} > 0,\ \ \overset{\infty}{H}_{ss} = X \times R > 0$$

Where $X$ defines corners of the simplex in projected space

$$\overset{\infty}{H}_{ss} = X \times R$$

$$X^{T} = R \times \overset{\infty}{H}_{ss}^{T}$$

$\overset{\infty}{V}_{ss}, \overset{\infty}{W}_{ss}, \overset{\infty}{H}_{ss}$ – row normalized matrices

produced by Sinkhorn procedure ($\overset{\infty}{V}_{ss} = \overset{\infty}{W}_{ss} \times \overset{\infty}{H}_{ss}$)

*For simplex of samples in a features space*

*1. Find projection vectors $S$ for $\overset{\infty}{V}_{gs}$ so that*

$$\min_{S} \left\| \overset{\infty}{V}_{fs} - S^{T} \times S \times \overset{\infty}{V}_{fs} \right\| \quad \underset{\overset{\infty}{V}_{fs}^{T}\vec{S}_1 = c\mathbb{I}_N}{s.t.\ \ S\times S^{T} = \mathbb{I}_{K\times K}}$$

Where: $\vec{S}_2, \ldots, \vec{S}_K$ could be found as left singular vectors of the centered matrix ${\overset{\infty}{V}_{fs}}_{c}$.

*2. Solve the NMF/deconvolution problem for this projected space*

$$\min_{\overset{\infty}{H}_{gs},\, \Omega} \left\| S \times \overset{\infty}{V}_{fs} - \Omega \times \overset{\infty}{H}_{fs} \right\| \quad s.t.\ \ \overset{\infty}{W}_{fs} = S^{T} \times \Omega > 0,\ \ \overset{\infty}{H}_{fs} > 0$$

Where $\Omega$ defines corners of the simplex in projected space

$$\overset{\infty}{W}_{fs} = S^{T} \times \Omega$$

$$\Omega = S \times \overset{\infty}{W}_{fs}$$

$\overset{\infty}{V}_{fs}, \overset{\infty}{W}_{fs}, \overset{\infty}{H}_{fs}$ – column normalized matrices

produced by Sinkhorn procedure ($\overset{\infty}{V}_{fs} = \overset{\infty}{W}_{fs} \times \overset{\infty}{H}_{fs}$)

### Merge two problems to obtain single optimization equation.

Once we found $R$, we can say that:

$$\overset{\infty}{V}_{ss}{}^{T} - R^T \times R \times \overset{\infty}{V}_{ss}{}^{T} = 0$$

Which in transposed form could be represented as

$$\overset{\infty}{V}_{ss} - \overset{\infty}{V}_{ss} \times R^T \times R = 0$$

Alternatively, if we found $S$ we can say that:

$$\overset{\infty}{V}_{fs} - S^T \times S \times \overset{\infty}{V}_{fs} = 0$$

But $\overset{\infty}{V}_{fs} = \overset{\infty}{V}_{ss} \times \frac{N}{M}$

$$\overset{\infty}{V}_{ss} - \frac{M}{N} S^T \times S \times \overset{\infty}{V}_{ss} \times \frac{N}{M} = 0$$

Which means

$$\overset{\infty}{V}_{ss} = S^T \times S \times \overset{\infty}{V}_{ss}$$

Then we can merge equations for $S$ and $R$ into single equation

$$\boxed{\overset{\infty}{V}_{ss} - S^T \times S \times \overset{\infty}{V}_{ss} \times R^T \times R = 0}$$

Since we know that $\overset{\infty}{W}_{ss} = \overset{\infty}{W}_{fs} \times D$ and $\overset{\infty}{V}_{ss} = \overset{\infty}{W}_{ss} \times \overset{\infty}{H}_{ss}$, this equation could be written as

$$\overset{\infty}{V}_{ss} - S^T \times S \times \overset{\infty}{W}_{fs} \times D \times \overset{\infty}{H}_{ss} \times R^T \times R = 0$$

Which is equivalent to

$$S \times \overset{\infty}{V}_{ss} \times R^T - S \times \overset{\infty}{W}_{fs} \times D \times \overset{\infty}{H}_{ss} \times R^T = 0$$

Let's define projection coordinates as we already did before:

$$\Omega = S \times \overset{\infty}{W}_{fs}$$

$$\overset{\infty}{H}_{ss} = X \times R \Leftrightarrow X = \overset{\infty}{H}_{ss} \times R^T$$

$$\boxed{S \times \overset{\infty}{V}_{ss} \times R^T = \Omega \times D \times X}$$

So, the result optimization problem for X and Omega can formulated as

$$\boxed{\begin{array}{l} \min\limits_{\substack{\Omega \\ X \\ D}} \left\| S \times \overset{\infty}{V}_{ss} \times R^T - \Omega \times D \times X \right\| \\ \qquad s.t. \quad \overset{\infty}{V}_{ss} \vec{R}_1 = c\mathbb{I}_M \\ \qquad\qquad \overset{\infty}{V}{}_{ss}^{T} \vec{S}_1 = c\frac{M}{N}\mathbb{I}_N \\ \qquad\qquad \overset{\infty}{H}_{ss} = X \times R > 0 \\ \qquad\qquad \overset{\infty}{W}_{gs} = S^T \times \Omega > 0 \end{array}}$$

### Summary

Using properties of Sinkhorn-transformed matrices we were able to connect two optimization problems into a single one. But for this problem it is still not clear how to find $R$ and $S$

*1.Find projection vectors $R$ and $S$ for $\overset{\infty}{V}_{ss}$ and $\overset{\infty}{V}_{fs}$ so that*

$$\min_{\substack{S\\R}} \left\| \overset{\infty}{V}_{ss} - S^T \times S \times \overset{\infty}{V}_{ss} \times R^T \times R \right\| \quad s.t. \quad \begin{array}{l} S \times S^T = \mathbb{I}_{K\times K} \\ R \times R^T = \mathbb{I}_{K\times K} \\ \overset{\infty}{V}_{ss}\vec{R}_1 = c\mathbb{I}_M \\ \overset{\infty}{V}{}_{ss}^{T}\vec{S}_1 = c\frac{M}{N}\mathbb{I}_N \end{array}$$

*2. Solve the NMF/deconvolution problem for this projected space*

$$\min_{\substack{X\\ \Omega\\ D}} \left\| S \times \overset{\infty}{V}_{ss} \times R^T - \Omega \times D \times X \right\| \quad s.t. \quad \begin{array}{l} \overset{\infty}{W}_{fs} = S^T \times \Omega > 0 \\ \overset{\infty}{H}_{ss} = X \times R > 0 \end{array}$$

Where $X$ and $\Omega$ define corners of the simplexes in projected space

$\overset{\infty}{V}_{ss}, \overset{\infty}{W}_{ss}, \overset{\infty}{H}_{ss}$ – row normalized matrices produced by Sinkhorn procedure ($\overset{\infty}{V}_{ss} = \overset{\infty}{W}_{ss} \times \overset{\infty}{H}_{ss}$)

$\overset{\infty}{V}_{fs}, \overset{\infty}{W}_{fs}, \overset{\infty}{H}_{fs}$ – column normalized matrices produced by Sinkhorn procedure ($\overset{\infty}{V}_{fs} = \overset{\infty}{W}_{fs} \times \overset{\infty}{H}_{fs}$)

$$\overset{\infty}{V}_{fs} = \overset{\infty}{V}_{ss} \times D_V = \underbrace{\overset{\infty}{W}_{ss} \times \overset{\infty}{D}_w}_{\overset{\infty}{W}_{fs}} \times \underbrace{\overset{\infty}{D}{}_w^{-1} \times \overset{\infty}{H}_{ss} \times D_V}_{\overset{\infty}{H}_{fs}}$$

$D = \overset{\infty}{D}{}_w^{-1}$ is diagonal matrix which connects matrices $\overset{\infty}{W}_{ss}$ and $\overset{\infty}{W}_{fs}$

$$\overset{\infty}{W}_{ss} = \overset{\infty}{W}_{fs} \times D$$

### Supplementary Note 9. Sum-to-one property is guaranteed for singular vectors of Sinkhorn transformed matrices

#### <u>What we did so far</u>

*<u>Introduced iterative Sinkhorn transformation for expression matrices</u>*

$$V \overset{\substack{row\\norm}}{\Longrightarrow} \underbrace{D_{v_0} \times V}_{\substack{\tilde{V}\\ simplex\ in\\ samples\ space}} \overset{\substack{column\\norm}}{\Longrightarrow} \underbrace{D_{v_0} \times V \times D_{v_1}}_{\substack{\tilde{\tilde{V}}\\ simplex\ in\\ features\ space}} \overset{\substack{row\\norm}}{\Longrightarrow} \ldots \overset{\substack{row\\norm}}{\Longrightarrow} \underset{\substack{simplex\ in\\ sample\ space}}{\overset{\infty}{V}_{ss}} \overset{\substack{column\\norm}}{\Longrightarrow} \underset{\substack{simplex\ in\\ features\ space}}{\overset{\infty}{V}_{fs}}$$

From which we got the constant relation between two simplexes defined.

$$\overset{\infty}{V}_{ss} \times \frac{N}{M} = \overset{\infty}{V}_{fs}$$

*<u>Formulated combined deconvolution problem in terms of new matrices</u>*

1. *Find projection vectors $R$ and $S$ for $\overset{\infty}{V}_{ss}$ and $\overset{\infty}{V}_{gs}$ so that*

$$\min_{\substack{S\\R}} \left\| \overset{\infty}{V}_{ss} - S^T \times S \times \overset{\infty}{V}_{ss} \times R^T \times R \right\| \quad s.t. \quad \begin{array}{l} S\times S^T=\mathbb{I}_{K\times K} \\ R\times R^T=\mathbb{I}_{K\times K} \\ \overset{\infty}{V}_{ss}\vec{R}_1=c\mathbb{I}_M \\ \overset{\infty}{V}{}_{ss}^T\vec{S}_1=c\frac{M}{N}\mathbb{I}_N \end{array}$$

2. *Solve the NMF/deconvolution problem for this projected space*

$$\min_{\substack{X\\ \Omega}} \left\| S \times \overset{\infty}{V}_{ss} \times R^T - \Omega \times D \times X \right\| \quad s.t. \quad \begin{array}{l} \overset{\infty}{W}_{fs}=S^T\times\Omega>0 \\ \overset{\infty}{H}_{ss}=X\times R>0 \end{array}$$

Where $X$ and $\Omega$ define corners of the simplexes in projected space

$\overset{\infty}{V}_{ss}, \overset{\infty}{W}_{ss}, \overset{\infty}{H}_{ss}$ – row normalized matrices

produced by Sinkhorn procedure ($\overset{\infty}{V}_{ss} = \overset{\infty}{W}_{ss} \times \overset{\infty}{H}_{ss}$)

$\overset{\infty}{V}_{fs}, \overset{\infty}{W}_{fs}, \overset{\infty}{H}_{gs}$ – column normalized matrices

produced by Sinkhorn procedure ($\overset{\infty}{V}_{fs} = \overset{\infty}{W}_{fs} \times \overset{\infty}{H}_{fs}$)

$$\overset{\infty}{V}_{fs} = \overset{\infty}{V}_{ss} \times D_V = \underbrace{\overset{\infty}{W}_{ss} \times \overset{\infty}{D}_w}_{\overset{\infty}{W}_{fs}} \times \underbrace{\overset{\infty}{D}{}_w^{-1} \times \overset{\infty}{H}_{ss} \times D_V}_{\overset{\infty}{H}_{fs}}$$

$D = \overset{\infty}{D}{}_w^{-1}$ is diagonal matrix which connects matrices $\overset{\infty}{W}_{ss}$ and $\overset{\infty}{W}_{fs}$

$$\overset{\infty}{W}_{ss} = \overset{\infty}{W}_{fs} \times D$$

### Lemma on SVD relation for Sinkhorn transformed matrices

#### *Lemma 1*

**Lemma 1.** Due to constant relation between Sinkhorn transformed matrices matrices $\overset{\infty}{V}_{ss}$ *and* $\overset{\infty}{V}_{fs}$ *have the same singular vectors and proportional singular values*

$$\overset{\infty}{V}_{ss} = \mathcal{U} \times \Sigma_{ss} \times \mathcal{V}^T$$

$$\overset{\infty}{V}_{fs} = \mathcal{U} \times \Sigma_{gs} \times \mathcal{V}^T = \mathcal{U} \times \frac{M}{N} \Sigma_{ss} \times \mathcal{V}^T$$

#### *Proof*

*For right singular vectors*

Let $\overset{\infty}{V}_{ss} = \mathcal{U} \times \Sigma_{ss} \times \mathcal{V}^T$ be singular value decomposition of matrix $\overset{\infty}{V}_{ss}$

Since $\mathcal{V}$ is a matrix of right singular vectors for matrix $\overset{\infty}{V}_{ss}$ and $\overset{\infty}{V}_{ss} = \frac{M}{N} \times \overset{\infty}{V}_{fs}$

$$\overset{\infty}{V}{}_{fs}^{T} \times \overset{\infty}{V}_{fs} \times \mathcal{V} = \frac{N^2}{M^2} \overset{\infty}{V}{}_{ss}^{T} \times \overset{\infty}{V}_{ss} \times \mathcal{V} = \frac{N^2}{M^2} \times \mathcal{V} \times \Sigma_{ss}^2$$

Where $\Sigma_{ss}^2$ is a diagonal matrix of squared singular values of matrix $\overset{\infty}{V}_{ss}$

This means that $\mathcal{V}$ is also a matrix of right singular vectors for matrix $\overset{\infty}{V}_{fs}$.with singular values $\Sigma_{fs} = \frac{N}{M} \Sigma_{ss}$. So right singular vector matrices are the same.

*For left singular vectors*

Let $\overset{\infty}{V}_{fs} = \mathcal{U} \times \Sigma_{fs} \times \mathcal{V}^T$ be the be singular value decomposition of matrix $\overset{\infty}{V}_{fs}$

Since $\mathcal{U}$ is a matrix of left singular vectors $\overset{\infty}{V}_{fs}$ and $\overset{\infty}{V}_{fs} = \frac{N}{M} \times \overset{\infty}{V}_{ss}$

$$\overset{\infty}{V}_{ss} \times \overset{\infty}{V}{}_{ss}^{T} \times S\mathcal{U}^T = \frac{M^2}{N^2} \overset{\infty}{V}_{fs} \times \overset{\infty}{V}{}_{fs}^{T} \times \mathcal{U}^T = \frac{M^2}{N^2} \times \mathcal{U}^T \times \Sigma_{fs}^2$$

This means that $\mathcal{U}$ is also a matrix of left singular vectors for matrix $\overset{\infty}{V}_{ss}$.with singular values $\Sigma_{ss} = \frac{M}{N} \Sigma_{ss}$. So left singular vector matrices are the same.

#### *Lemma 2*

**Lemma 2.** For Sinkhorn transformed matrices $\overset{\infty}{V}_{ss}$, $\overset{\infty}{V}_{gs}$, we know exact form of one of the vectors composing right ($\mathcal{V}^T$) and left ($\mathcal{U}$) singular vectors which are diagonal vectors $\vec{\mathcal{V}}_1^T = \begin{bmatrix} \frac{1}{\sqrt{N}} & \dots & \frac{1}{\sqrt{N}} \end{bmatrix}$ and $\vec{\mathcal{U}}_1^T = \begin{bmatrix} \frac{1}{\sqrt{M}} & \dots & \frac{1}{\sqrt{M}} \end{bmatrix}$

#### *Proof for a simplex of features in a samples space*

**Statement*:*** one of the vectors in $\mathcal{V}$ is $\frac{1}{\sqrt{N}} \mathbb{I}_N$

**Proof:** Since $\overset{\infty}{V}_{ss}$ is row normalized:

$$\overset{\infty}{V}_{ss} \times \mathbb{I}_N = \mathbb{I}_M$$

Then

$$\overset{\infty}{V}_{ss}^{T} \times \overset{\infty}{V}_{ss} \times \mathbb{I}_N = \overset{\infty}{V}_{ss}^{T} \times \mathbb{I}_M$$

But $\overset{\infty}{V}_{ss} = \frac{M}{N} \times \overset{\infty}{V}_{gs}$ and $\overset{\infty}{V}_{ss}^{T} = \frac{M}{N} \times \overset{\infty}{V}_{gs}^{T}$

$$\overset{\infty}{V}_{ss}^{T} \times \mathbb{I}_M = \frac{M}{N} \times \overset{\infty}{V}_{gs}^{T} \times \mathbb{I}_M$$

Since $\overset{\infty}{V}_{gs}$ is column normalized, $\overset{\infty}{V}_{gs}^{T}$ is row normalized.

$$\overset{\infty}{V}_{gs}^{T} \times \mathbb{I}_M = \mathbb{I}_N$$

This means that

$$\frac{M}{N} \times \overset{\infty}{V}_{gs}^{T} \times \mathbb{I}_M = \frac{M}{N} \times \mathbb{I}_N$$

This means

$$\overset{\infty}{V}_{ss}^{T} \times \overset{\infty}{V}_{ss} \times \mathbb{I}_N = \frac{M}{N} \times \mathbb{I}_N$$

Which means that $\mathbb{I}_N$ is an eigenvector for $\overset{\infty}{V}_{ss}^{T} \times \overset{\infty}{V}_{ss}$. Then normalized vector $\frac{1}{\sqrt{N}} \times \mathbb{I}_N$ is also an eigen vector for $\overset{\infty}{V}_{ss}^{T} \times \overset{\infty}{V}_{ss}$.

$$\overset{\infty}{V}_{ss}^{T} \times \overset{\infty}{V}_{ss} \times \frac{1}{\sqrt{N}} \times \mathbb{I}_N = \frac{M}{N} \times \frac{1}{\sqrt{N}} \times \mathbb{I}_N$$

This also means that $\frac{1}{\sqrt{N}} \times \mathbb{I}_N$ is a right singular vector for $\overset{\infty}{V}_{ss}$ and left singular vector for $\overset{\infty}{V}_{ss}^{T}$ with singular value $\sigma = \sqrt{\frac{M}{N}}$

Since $\overset{\infty}{V}_{ss}^{T} = \frac{M}{N} \times \overset{\infty}{V}_{gs}^{T}$, a vector $\frac{1}{\sqrt{N}} \times \mathbb{I}_N$ is also a left singular vector for $\overset{\infty}{V}_{gs}^{T}$ (or right singular vector for $\overset{\infty}{V}_{gs}$) with corresponding singular value $\sigma = \sqrt{\frac{N}{M}}$

Therefore, one of the vectors in the $\mathcal{V}$, e.g. $\mathcal{V}_1$, should be $\frac{1}{\sqrt{N}}\mathbb{I}_N$.

*Proof for a simplex of samples in a features space*

**Statement:** one of the vectors in $\mathcal{U}$ is $\frac{1}{\sqrt{M}}\mathbb{I}_M$

**Proof:** We know that $\overset{\infty}{V}_{gs}$ is column normalized then $\overset{\infty}{V}_{gs}^{T}$ is row normalized.

$$\overset{\infty}{V}_{gs}^{T} \times \mathbb{I}_M = \mathbb{I}_N$$

Then

$$\overset{\infty}{V}_{gs} \times \overset{\infty}{V}_{gs}^{T} \times \mathbb{I}_M = \overset{\infty}{V}_{gs} \times \mathbb{I}_N$$

But $\overset{\infty}{V}_{ss} \times \frac{N}{M} = \overset{\infty}{V}_{gs}$

$$\overset{\infty}{V}_{gs} \times \mathbb{I}_N = \frac{N}{M} \times \overset{\infty}{V}_{ss} \times \mathbb{I}_N$$

Since $\overset{\infty}{V}_{ss}$ is row normalized

$$\overset{\infty}{V}_{ss} \times \mathbb{I}_N = \mathbb{I}_M$$

Therefore

$$\frac{N}{M} \times \overset{\infty}{V}_{ss} \times \mathbb{I}_N = \frac{N}{M} \times \mathbb{I}_M$$

Which means

$$\overset{\infty}{V}_{gs} \times \overset{\infty}{V}{}_{gs}^{T} \times \mathbb{I}_M = \frac{N}{M} \times \mathbb{I}_M$$

Which means that $\mathbb{I}_M$ is an eigenvector for $\overset{\infty}{V}_{gs} \times \overset{\infty}{V}{}_{gs}^{T}$. Then normalized vector $\frac{1}{\sqrt{M}} \times \mathbb{I}_M$ is also an eigen vector for $\overset{\infty}{V}_{gs} \times \overset{\infty}{V}{}_{gs}^{T}$.

$$\overset{\infty}{V}_{gs} \times \overset{\infty}{V}{}_{gs}^{T} \times \frac{1}{\sqrt{M}} \times \mathbb{I}_M = \frac{N}{M} \times \frac{1}{\sqrt{M}} \times \mathbb{I}_M$$

This also means that $\frac{1}{\sqrt{M}} \times \mathbb{I}_M$ is a left singular vector for $\overset{\infty}{V}_{gs}$ and a right singular vector for $\overset{\infty}{V}{}_{gs}^{T}$ with singular value $\sigma = \sqrt{\frac{N}{M}}$.

Since $\overset{\infty}{V}_{ss} \times \frac{N}{M} = \overset{\infty}{V}_{gs}$, a vector $\frac{1}{\sqrt{M}} \times \mathbb{I}_M$ is also a left singular vector for $\overset{\infty}{V}_{ss}$ (or right singular vector for $\overset{\infty}{V}{}_{ss}^{T}$) with corresponding singular value $\sigma = \sqrt{\frac{M}{N}}$.

That's why $\vec{\mathcal{U}}_1$ should be $\frac{1}{\sqrt{M}}\mathbb{I}_M$.

### Theorem on SVD properties of Sinkhorn transformed matrices

We will show that matrices $\boldsymbol{R}$ and $\boldsymbol{S}$ obtained from Sikhorn-transformed matrices using SVD already incorporate sum-to-one restrictions.

#### *The main theorem*

***The main theorem.*** *For Sinkhorn transformed matrices $\overset{\infty}{V}_{ss}$, $\overset{\infty}{V}_{gs}$ $\left(\overset{\infty}{V}_{gs} = \overset{\infty}{V}_{ss} \times D_V\right)$ of non-negative matrix V solution to projection optimization problems.*

$$\min_{R} \left\| \overset{\infty}{V}{}_{ss}^{T} - R^T \times R \times \overset{\infty}{V}{}_{ss}^{T} \right\|_{\substack{s.t.\ R \times R^T = \mathbb{I}_{K\times K} \\ \overset{\infty}{V}_{ss}\vec{R}_1 = c\mathbb{I}_M}} \quad \min_{S} \left\| \overset{\infty}{V}_{fs} - S^T \times S \times \overset{\infty}{V}_{fs} \right\|_{\substack{s.t.\ S \times S^T = \mathbb{I}_{K\times K} \\ \overset{\infty}{V}{}_{gs}^{T}\vec{S}_1 = c\mathbb{I}_N}}$$

*can be found as follows:*

*$R^T$ could be found as left singular matrix of $\overset{\infty}{V}{}_{ss}^{T}$ and right singular vectors of $\overset{\infty}{V}_{fs}$*

*$S^T$ could be found as right singular matrix of $\overset{\infty}{V}{}_{ss}^T$ and left singular vectors of $\overset{\infty}{V}_{fs}$*

*<u>Proof</u>*

The unconstrained form of each optimization problem

$$\min_{\substack{R \\ s.t.\quad R\times R^T=\mathbb{I}_{K\times K}}} \left\| \overset{\infty}{V}{}_{ss}^T - R^T \times R \times \overset{\infty}{V}{}_{ss}^T \right\| \quad \min_{\substack{S \\ s.t.\quad S\times S^T=\mathbb{I}_{K\times K}}} \left\| \overset{\infty}{V}_{fs} - S^T \times S \times \overset{\infty}{V}_{gs} \right\| \qquad (*)$$

could be solved using SVD vectors (Yuan & Oja, 2005) where $R^T$ matrix consists of left singular vectors for $\overset{\infty}{V}{}_{ss}^T$ and $S^T$ matrix consists of left singular vectors for $\overset{\infty}{V}_{fs}$ so that

$$R^T \times R \times \overset{\infty}{V}{}_{ss}^T = \overset{\infty}{V}{}_{ss}^T \quad S^T \times S \times \overset{\infty}{V}_{fs} = \overset{\infty}{V}_{fs}$$

Since $\overset{\infty}{V}_{fs}$ and $\overset{\infty}{V}_{ss}$ are Sinkhorn transformed matrices, then according to **Lemma 1** their left and right singular vectors matrices are the same.

$$\overset{\infty}{V}_{ss} = S^T \times \Sigma_{ss} \times R$$

$$\overset{\infty}{V}_{gs} = S^T \times \Sigma_{gs} \times R$$

$$\Sigma_{ss} = \frac{N}{M}\Sigma_{gs}$$

Moreover since $R^T$ is the left singular matrix for $\overset{\infty}{V}{}_{ss}^T$, then it is also the right singular matrix for $\overset{\infty}{V}_{ss}$ and vise versa since $S^T$ is the left singular matrix for $\overset{\infty}{V}_{gs}$, then it is also the right singular matrix for $\overset{\infty}{V}{}_{gs}^T$.

This symmetry makes matrices $R$ and $S$ the sought after projection vectors for both features and samples in unconstrained case.

Furthermore, according to **Lemma 2** $\frac{1}{\sqrt{N}}\mathbb{I}_N$ and $\frac{1}{\sqrt{M}}\mathbb{I}_M$ are within these matrices $R$ and $S$. Since SVD decomposition is unique up to the order of the vectors, we can select this pair of vectors as first factors of our projection operator and name these vectors as $\vec{R}_1$ and $\vec{S}_1$ respectively.

*For simplex of features in a samples space*

Since ($\vec{R}_1$,$\vec{S}_1$) is a pair of right and left singular vectors for $\overset{\infty}{V}_{ss}$ with singular value $\sigma_1 = \sqrt{\frac{M}{N}}$, then

$$\overset{\infty}{V}_{ss} \times \vec{R}_1 = \sqrt{\frac{M}{N}} \times \vec{S}_1$$

But $\vec{S}_1 = \frac{1}{\sqrt{M}}\mathbb{I}_M$

$$\overset{\infty}{V}_{ss} \times \vec{R}_1 = \sqrt{\frac{M}{N}} \times \frac{1}{\sqrt{M}}\mathbb{I}_M$$

Which is equivalent to

$$\overset{\infty}{V}_{ss} \times \vec{R}_1 = \frac{1}{\sqrt{N}} \mathbb{I}_M$$

Which means that projection of all row vectors from $\overset{\infty}{V}_{ss}$ (column vectors of $\overset{\infty}{V}{}_{ss}^{T}$) is constant.

Therefore, the matrix of left singular vectors $R^T$(for matrix $\overset{\infty}{V}{}_{ss}^{T}$) which deliver global minimum value for unconstrained problem (*) formulated for Sinkhorn transformed matrix $\overset{\infty}{V}{}_{ss}^{T}$ always satisfies the constrains we defined for $R^T$ matrix in constrained case. Which means that singular vectors solution also delivers optimum for constrained problem solution. As what to be shown.

*For simplex of samples in features space*

Since $(\vec{S}_1, \vec{R}_1)$ is pair of right and left singular vectors for $\overset{\infty}{V}{}_{fs}^{T}$ with singular value $\sigma_1 = \sqrt{\frac{N}{M}}$, then

$$\overset{\infty}{V}{}_{fs}^{T} \times \vec{S}_1 = \sqrt{\frac{N}{M}} \times \vec{R}_1$$

But $\vec{R}_1 = \frac{1}{\sqrt{N}} \mathbb{I}_N$

$$\overset{\infty}{V}{}_{fs}^{T} \times \vec{S}_1 = \sqrt{\frac{N}{M}} \times \frac{1}{\sqrt{N}} \mathbb{I}_N$$

Which is equivalent to

$$\overset{\infty}{V}{}_{fs}^{T} \times \vec{S}_1 = \frac{1}{\sqrt{M}} \mathbb{I}_N$$

Which means that projection of all row vectors from $\overset{\infty}{V}{}_{fs}^{T}$ (column vectors of $\overset{\infty}{V}_{fs}$) is constant. Therefore, left singular vectors matrix $S^T$(for matrix $\overset{\infty}{V}_{gs}$) which deliver global minimum value for unconstrained problem (*) formulated for Sinkhorn transformed matrix $\overset{\infty}{V}_{gs}$ always satisfies the constrains we defined for $S^T$ matrix in constrained case. Which means that singular vectors solution also delivers optimum for constrained problem solution. As what to be shown.

### <u>Corollary on geometrical interpretation of obtained SVD projection solution</u>

***Corollary***. *For Sinkhorn transformed matrices $\overset{\infty}{V}_{ss}$, $\overset{\infty}{V}_{gs}$ are diagonal vectors $\vec{R}_1^T = \begin{bmatrix} \frac{1}{\sqrt{N}} & \dots & \frac{1}{\sqrt{N}} \end{bmatrix}$ and $\vec{S}_1^T = \begin{bmatrix} \frac{1}{\sqrt{M}} & \dots & \frac{1}{\sqrt{M}} \end{bmatrix}$ reflect the shift of the simplex hyperplane from the zero point.*

*<u>Geometrical interpretation of features projection in samples space</u>*

Due to exact diagonal form and orthogonality property obtained for $\vec{R}_1^T$ in the main theorem we can derive the for one of the coordinate values for each NMF/deconvolution solution vector.

Since $X^T = R \times \overset{\infty}{H}{}_{ss}^{T}$

The first row of the result $X^T$ is a product of a row vector $\vec{R}_1{}^T$ and columns of $\overset{\infty}{H}{}_{ss}^{T}$.However $\overset{\infty}{H}{}_{ss}^{T}$ is column normalized, which makes first row of $X^T$ equal to $\frac{1}{\sqrt{N}} \mathbb{I}_N$.

$$\forall k \in [1, K] \;\; x_{1,k} = \sum_{i=1}^{N} r_{1,i} * \left(\overset{\infty}{h_{ss}^{T}}\right)_{i,k} = \sum_{i=1}^{N} \frac{1}{\sqrt{N}} * \left(\overset{\infty}{h_{ss}^{T}}\right)_{i,k} = \frac{1}{\sqrt{N}} \sum_{i=1}^{N} \left(\overset{\infty}{h_{ss}^{T}}\right)_{i,k} = \frac{1}{\sqrt{N}}$$

Meaning that $X$ matrix has predefined first column.

$$X = \begin{bmatrix} \frac{1}{\sqrt{N}} & x_{1,2} & \cdots & x_{1,K} \\ \vdots & \vdots & \ddots & \vdots \\ \frac{1}{\sqrt{N}} & x_{K,2} & \cdots & x_{K,K} \end{bmatrix}$$

This means that first coordinate for each of the projected points in the basis $R_1, R_2, \dots, R_K$ will always be constant, specifying the shift of the simplex corners (and the whole hyperplane) from the center of coordinates. At the same time vectors $\vec{R}_2, \dots, \vec{R}_K$ still describe the entire $(K - 1)$-dimensional simplex structure.

*<u>Geometrical interpretation of samples projection in features space</u>*

Due to exact diagonal form and orthogonality property obtained for $\vec{S}_1^T$ in the main theorem we can derive the for one of the coordinate values for each NMF/deconvolution solution vector

Similarly, to what we did for $X$ matrix, since $\Omega = S \times \overset{\infty}{W_{gs}}$

The first row of the result $\Omega$ is a product of a row vector $\vec{S}_1^{\,T}$ and columns of $\overset{\infty}{W_{gs}}$, but $\overset{\infty}{W_{gs}}$ is column normalized, which makes first row of $\Omega$ equal to $\frac{1}{\sqrt{M}} \mathbb{I}_M$.

$$\forall k \in [1, K] \;\; \omega_{1,k} = \sum_{j=1}^{M} s_{1,j} * \left(\overset{\infty}{w_{gs}}\right)_{j,k} = \sum_{j=1}^{M} \frac{1}{\sqrt{M}} * \left(\overset{\infty}{w_{gs}}\right)_{j,k} = \frac{1}{\sqrt{M}} \sum_{j=1}^{M} \left(\overset{\infty}{w_{gs}}\right)_{j,k} = \frac{1}{\sqrt{M}}$$

Meaning that $\Omega$ matrix has predefined first row, specifying the shift of the simplex corners (and the whole hyperplane) from the center of coordinates.

$$\Omega = \begin{bmatrix} \frac{1}{\sqrt{M}} & \cdots & \frac{1}{\sqrt{M}} \\ \omega_{2,1} & \cdots & \omega_{2,K} \\ \vdots & \ddots & \vdots \\ \omega_{K,1} & \cdots & \omega_{K,K} \end{bmatrix}$$

At the same time vectors $\vec{S}_2, \dots, \vec{S}_K$ still describe the entire $(K - 1)$-dimensional simplex structure.

**<u>Incorporate known vectors into combined problem formulations.</u>**

With all this we can update the original task with already known vectors and coordinate values

*<u>1.Find projection vectors $R$ and $S$ for $\overset{\infty}{V}_{ss}$ and $\overset{\infty}{V}_{gs}$ so that</u>*

$$\min_{\substack{S\\R}} \left\| \overset{\infty}{V}_{ss} - S^T \times S \times \overset{\infty}{V}_{ss} \times R^T \times R \right\| \quad s.t. \quad \begin{array}{l} S\times S^T = \mathbb{I}_{K\times K} \\ R\times R^T = \mathbb{I}_{K\times K} \\ \overset{\infty}{V}_{ss}\vec{R}_1 = c\mathbb{I}_M \\ \overset{\infty}{V}{}_{ss}^{T}\vec{S}_1 = c\frac{M}{N}\mathbb{I}_N \end{array}$$

Where

$$S^T \text{ could be found as left singular vectors of } \overset{\infty}{V}_{ss} \text{ using SVD}$$

$$R^T \text{ could be found as right singular vectors of } \overset{\infty}{V}_{ss} \text{ using SVD}$$

$$R = \begin{bmatrix} \frac{1}{\sqrt{N}} & \frac{1}{\sqrt{N}} & \cdots & \frac{1}{\sqrt{N}} \\ r_{2,1} & r_{2,2} & \cdots & r_{2,N} \\ \vdots & \vdots & \ddots & \vdots \\ r_{K,1} & r_{K,2} & \cdots & r_{K,N} \end{bmatrix} \quad S = \begin{bmatrix} \frac{1}{\sqrt{M}} & \frac{1}{\sqrt{M}} & \cdots & \frac{1}{\sqrt{M}} \\ s_{2,1} & s_{2,2} & \cdots & s_{2,M} \\ \vdots & \vdots & \ddots & \vdots \\ s_{K,1} & s_{K,2} & \cdots & s_{K,M} \end{bmatrix}$$

*<u>2. Solve deconvolution problem for this projected space</u>*

$$\min_{\substack{X\\ \Omega\\ D}} \left\| S \times \overset{\infty}{V}_{ss} \times R^T - \Omega \times D \times X \right\| \quad s.t. \quad \begin{array}{l} \overset{\infty}{W}_{gs} = S^T \times \Omega > 0 \\ \overset{\infty}{H}_{ss} = X \times R > 0 \end{array}$$

Where

$X$ and $\Omega$ define corners of the simplexes in projected space

$$X = \begin{bmatrix} \frac{1}{\sqrt{N}} & x_{1,2} & \cdots & x_{1,K} \\ \vdots & \vdots & \ddots & \vdots \\ \frac{1}{\sqrt{N}} & x_{K,2} & \cdots & x_{K,K} \end{bmatrix} \quad \Omega = \begin{bmatrix} \frac{1}{\sqrt{M}} & \cdots & \frac{1}{\sqrt{M}} \\ \omega_{2,1} & \cdots & \omega_{2,K} \\ \vdots & \ddots & \vdots \\ \omega_{K,1} & \cdots & \omega_{K,K} \end{bmatrix}$$

$\overset{\infty}{V}_{ss}, \overset{\infty}{W}_{ss}, \overset{\infty}{H}_{ss}$ – row normalized matrices produced by Sinkhorn procedure

$$(\overset{\infty}{V}_{ss} = \overset{\infty}{W}_{ss} \times \overset{\infty}{H}_{ss})$$

$\overset{\infty}{V}_{gs}, \overset{\infty}{W}_{gs}, \overset{\infty}{H}_{gs}$ – column normalized matrices produced by Sinkhorn procedure

$$(\overset{\infty}{V}_{fs} = \overset{\infty}{W}_{fs} \times \overset{\infty}{H}_{fs})$$

$$\overset{\infty}{V}_{fs} = \overset{\infty}{V}_{ss} \times D_V = \underbrace{\overset{\infty}{W}_{ss} \times \overset{\infty}{D}_w}_{\overset{\infty}{W}_{fs}} \times \underbrace{\overset{\infty}{D}{}_w^{-1} \times \overset{\infty}{H}_{ss} \times D_V}_{\overset{\infty}{H}_{fs}}$$

$D = \overset{\infty}{D}{}_w^{-1} > 0$ is diagonal matrix which connects matrices $\overset{\infty}{W}_{ss}$ and $\overset{\infty}{W}_{fs}$

$$\overset{\infty}{W}_{ss} = \overset{\infty}{W}_{fs} \times D$$

### Supplementary Note 10. Computational formulation of the optimization problem

So far, we got a single optimization equation for our deconvolution problem:

*1.Find projection vectors $R$ and $S$ for $\overset{\infty}{V}_{ss}$ and $\overset{\infty}{V}_{gs}$ so that*

$$\min_{\substack{S\\R}} \left\| \overset{\infty}{V}_{ss} - S^T \times S \times \overset{\infty}{V}_{ss} \times R^T \times R \right\| \quad \begin{array}{l} s.t. \quad S \times S^T = \mathbb{I}_{K\times K} \\ \qquad\; R \times R^T = \mathbb{I}_{K\times K} \\ \qquad\; \overset{\infty}{V}_{ss}\vec{R}_1 = c\mathbb{I}_M \\ \qquad\; \overset{\infty}{V}{}_{ss}^{T}\vec{S}_1 = c\frac{M}{N}\mathbb{I}_N \end{array}$$

Where

$$S^T \text{ could be found as left singular vectors of } \overset{\infty}{V}_{ss} \text{ using SVD}$$

$$R^T \text{ could be found as right singular vectors of } \overset{\infty}{V}_{ss} \text{ using SVD}$$

*2. Solve deconvolution problem for this projected space*

$$\min_{\substack{X\\ \Omega\\ D}} \left\| S \times \overset{\infty}{V}_{ss} \times R^T - \Omega \times D \times X \right\| \quad \begin{array}{l} s.t. \quad \overset{\infty}{W}_{fs} = S^T \times \Omega > 0 \\ \qquad\; \overset{\infty}{H}_{ss} = X \times R > 0 \end{array} \tag{1}$$

Where $X$ and $\Omega$ define corners of the simplexes in projected space

$\overset{\infty}{V}_{ss}$ – row normalized matrix produced by Sinkhorn procedure, representing simplex in a sample space.

$\overset{\infty}{V}_{fs}$ – column normalized matrix produced by Sinkhorn procedure, representing simplex in a features space.

$\overset{\infty}{W}_{ss}, \overset{\infty}{H}_{ss}$ – row normalized matrices found during optimization ($\overset{\infty}{V}_{ss} = \overset{\infty}{W}_{ss} \times \overset{\infty}{H}_{ss}$)

$\overset{\infty}{W}_{gs}, \overset{\infty}{H}_{gs}$ – column normalized matrices found during optimization ($\overset{\infty}{V}_{fs} = \overset{\infty}{W}_{fs} \times \overset{\infty}{H}_{fs}$)

$$\overset{\infty}{V}_{gs} = \overset{\infty}{V}_{ss} \times D_V = \underbrace{\overset{\infty}{W}_{ss} \times D_w}_{\overset{\infty}{W}_{fs}} \times \underbrace{\overset{\infty}{D}{}_w^{-1} \times \overset{\infty}{H}_{ss} \times D_V}_{\overset{\infty}{H}_{fs}}$$

$D = \overset{\infty}{D}{}_w^{-1}$ is diagonal matrix which connects matrices $\overset{\infty}{W}_{ss}$ and $\overset{\infty}{W}_{fs}$

$$\overset{\infty}{W}_{ss} = \overset{\infty}{W}_{fs} \times D$$

Next, we want to computationally solve this optimization problem (1). Gradient descent has been chosen as an algorithm to accomplish this task.

Let's note that it is possible to solve this problem in multiple ways. Originally, we have 3 variable matrices to optimize ($X, \Omega, D$). but in this section, we show that due to geometrical properties of Sinkhorn transformed matrices it is possible to simplify optimization problem significantly.

We start with definition of a cost function in a general way and proceed with a most straightforward and interpretable training procedure which formulates update rules for each variable according to cost function derivatives. As the following step we discuss alternative approaches to optimization which involve substitution of original optimization variables and how these substitutions change original cost function and respective derivatives.

The last step will be discussing the hyperparameters of the training procedure.

### The cost function.

To apply gradient descent algorithm, we need to define a cost function, which we will optimize with respect to all 3 variables $(X, \Omega, D)$.

The main part of the cost function is naturally following from the unconstrained form of the given problem and could be expressed as a deconvolution error in terms of Frobenius norm.

$$\boxed{C(X, \Omega, D) = \left\| S \times \overset{\infty}{V}_{ss} \times R^T - \Omega \times D \times X \right\|_F^2} \quad (2)$$

#### *Incorporate non-negativity constraints.*

Then we want to add constraints defined.

Following biological nature of data, we used constraints of positivity for matrix of proportions $\overset{\infty}{H}_{ss}$ and pure components $\overset{\infty}{W}_{gs}$.

$$\overset{\infty}{H}_{ss} = X \times R > 0$$

$$\overset{\infty}{W}_{fs} = S^T \times \Omega > 0$$

Such constraints could be considered in the total optimization function as a hinge function terms. Where hinge function for matrix is formulated as sum of absolute values of negative elements:

$$\|A\|_h = \sum_i \sum_j \max(-A_{i,j}, 0)$$

This function allows us to control number of negative elements and their values at the same time.

Since initial optimized function does not incorporate given constraints, hinge terms should be added explicitly and are subject to optimization. Penalty method was used to include into const function (2) these additional terms.

$$\boxed{C(X, \Omega, D) = \left\| S \times \overset{\infty}{V}_{ss} \times R^T - \Omega \times D \times X \right\|_F^2 + \beta * \|S^T \times \Omega\|_h + \lambda * \|X \times R\|_h} \quad (3)$$

Where $\beta$ and $\lambda$ – penalty coefficients for negativity cost functions.

### The training process and update rules

After formulation of the cost function (3) we need define the training procedure. Let’s note that in fact multiple approaches to optimization possible.

Since our optimization requires updates in multiple spaces simultaneously we applied block-wise gradient descent to perform separate updates for each variable. We kept these updates in strict order to keep search synchronized between 2 simplexes.

$$X \rightarrow D \rightarrow \Omega \rightarrow D \rightarrow X \rightarrow \cdots$$

Once we decided with strategies, we need to calculate partial derivative of the cost function (5) with respect to all variables $(X, \Omega, D)$

*Partial derivative with respect to $X$ and update rule*

For each iteration $i$ of the learning algorithm the update rule for $X$will be the following:

$$\boxed{X_{i+1} = X_i - \chi * \nabla F_{X_i}} \tag{4}$$

Where $\chi$ is a learning rate for $X$, which will be defined in detail further below.

And $\nabla F_{X_i}$ is a derivative of cost function with respect to matrix $X$

$$\boxed{\begin{gathered}\nabla F_{X_i} = -2 * D \times \Omega_i^{\mathrm{T}} \times (S \times V_{SS} \times R^T - \Omega_i \times D \times X_i) \\ +\lambda * \frac{\partial \|X \times R\|_h}{\partial X}\end{gathered}} \tag{5}$$

Term $\frac{\partial \|X \times R\|_h}{\partial X}$ in equation (5) is a $K \times K$ matrix where rows are partial derivatives of the hinge function with respect to rows of $X$:

$$\frac{\partial \|X \times R\|_h}{\partial X} = \begin{bmatrix} \frac{\|X \times R\|_h}{\partial X_{1,*}} \\ \vdots \\ \frac{\|X \times R\|_h}{\partial X_{K,*}} \end{bmatrix}$$

$$\forall l \in [1, K] \left( \frac{\|X \times R\|_h}{\partial X_{l,*}} \right)^T = \sum_{j=1}^{N} \begin{cases} -\vec{R}_{*,j}, & if\ (X \times R)_{l,j} < 0 \\ 0, & if\ (X \times R)_{l,j} \geq 0 \end{cases}$$

*Partial derivative with respect to $\Omega$ and update rule*

Similarly, for each iteration $i$ of the learning algorithm the update rule for $\Omega$ will be the following:

$$\boxed{\Omega_{i+1} = \Omega_i - \phi * \nabla F_{\Omega_i}} \tag{6}$$

Where $\phi$ is a learning rate for $\Omega$, which will be defined in detail further below.

And $\nabla F_{\Omega_i}$ is a derivative of cost function with respect to matrix $\Omega$

$$\boxed{\begin{gathered}\nabla F_{\Omega_i} = -2 * (S \times V_{SS} \times R^T - \Omega_i \times D \times X_{i+1}) \times X_{i+1}^T \times D \\ +\beta * \frac{\partial \|S^T \times \Omega\|_h}{\partial \Omega}\end{gathered}} \tag{7}$$

Term $\frac{\partial \|S^T \times \Omega\|_h}{\partial \Omega}$ in equation (7) is a $K \times K$ matrix, where columns are partial derivatives of the hinge function with respect to columns of $\Omega$

$$\forall l \in [1, K] \left(\frac{\|S^T \times \Omega\|_h}{\partial \Omega_{*,l}}\right)^T = \sum_{j=1}^{N} \begin{cases} -\vec{S}_{*,j}, & if\ (S^T \times \Omega)_{l,j} < 0 \\ 0, & if\ (S^T \times \Omega)_{l,j} \geq 0 \end{cases}$$

$$\frac{\partial \|S^T \times \Omega\|_h}{\partial \Omega} = \left[\frac{\|S^T \times \Omega\|_h}{\partial \Omega_{*,1}} \quad \cdots \quad \frac{\|S^T \times \Omega\|_h}{\partial \Omega_{*,K}}\right]$$

*Update rule for* $D$

For $D$ matrix we do updates without gradients.

Our goal is to optimize the Frobenius norm (2) with respect to $D$.

$$\min_{D} \left\| S \times \overset{\infty}{V_{ss}} \times R^T - \Omega \times D \times X \right\|_F^2$$

But D is diagonal:

$$D = \begin{bmatrix} d_1 & \cdots & 0 \\ \vdots & \ddots & \vdots \\ 0 & \cdots & d_k \end{bmatrix}$$

So, the term $\Omega \times D \times X$ could be written in terms of these diagonal elements:

$$\Omega \times D \times X = \begin{bmatrix} \sum_{i=1}^{K} d_i \omega_{1,i} x_{i,1} & \dots & \sum_{i=1}^{K} d_i \omega_{1,i} x_{i,K} \\ \vdots & \ddots & \vdots \\ \sum_{i=1}^{K} d_i \omega_{K,i} x_{i,1} & \dots & \sum_{i=1}^{K} d_i \omega_{K,i} x_{i,K} \end{bmatrix} =$$

$$= \sum_{i=1}^{K} d_i \begin{bmatrix} \omega_{1,i} x_{i,1} & \dots & \omega_{1,i} x_{i,K} \\ \vdots & \ddots & \vdots \\ \omega_{K,i} x_{i,K} & \dots & \omega_{K,i} x_{i,K} \end{bmatrix} = \sum_{j=1}^{K} d_j \times \begin{bmatrix} \omega_{1,j} \\ \vdots \\ \omega_{K,j} \end{bmatrix} \times [x_{j,1} \quad \cdots \quad x_{j,K}]$$

Which means

$$\boxed{\Omega \times D \times X = \sum_{j=1}^{K} d_j \times \begin{bmatrix} \omega_{1,j} \\ \vdots \\ \omega_{K,j} \end{bmatrix} \times [x_{j,1} \quad \cdots \quad x_{j,K}]}$$

Therefore, initial problem could be rewritten as

$$\min_{\vec{D}} \left\| S \times \overset{\infty}{V_{ss}} \times R^T - \sum_{j=1}^{K} d_j \times \begin{bmatrix} \omega_{1,j} \\ \vdots \\ \omega_{K,j} \end{bmatrix} \times [x_{j,1} \quad \cdots \quad x_{j,K}] \right\|_F^2$$

Let's denote $Q_j = \begin{bmatrix} \omega_{1,j} \\ \vdots \\ \omega_{K,j} \end{bmatrix} \times [x_{j,1} \quad \cdots \quad x_{j,K}]$

$$\min_{\vec{D}} \left\| S \times \overset{\infty}{V_{ss}} \times R^T - \sum_{j=1}^{K} d_j \times Q_j \right\|_F^2$$

Minimization of Frobenius norm of matrices is equivalent of minimization of vectorized matrices, where values of $K \times K$ matrices are represented as row vectors of size $K^2$. The same problem could be formulated in vectorized form.

$$\min_{\vec{D}} \left\| vec(S \times \overset{\infty}{V_{ss}} \times R^T) - \sum_{j=1}^{K} d_j \times vec(Q_j) \right\|_F^2$$

Not let's denote as $Q(\mathrm{X}, \Omega)$ matrix whose columns are given by.

$$\forall j \in [1, K] \mathrm{Q}_{*,j} = vec(Q_j)$$

Then problem could be rewritten as

$$\boxed{\min_{\vec{D}} \left\| vec\left(S \times \overset{\infty}{V_{ss}} \times R^T\right) - Q(\mathrm{X}, \Omega) \times \vec{D} \right\|_F^2} \qquad (8)$$

$$\begin{aligned} Where \quad & vec\left(S \times \overset{\infty}{V_{ss}} \times R^T\right) - \ row\ vector\ of\ size\ K^2 \\ & Q(\mathrm{X}, \Omega) - matrix\ of\ size\ K^2 \times K \\ & \vec{D} - row\ vector\ of\ size\ \ K \end{aligned}$$

Moreover $\vec{D}$ is non-negative by its definition.

Then this optimization task with non-negativity constraint for $\vec{D}$ is exactly a formulation of non-negative least squares problem. Which means that $\vec{D}$ could be found as solution of this problem.

$$\vec{D} = \underset{x}{argmin} \left\| vec\left(S \times \overset{\infty}{V_{ss}} \times R^T\right) - Q(X, \Omega) \times x \right\|_F^2 \qquad (9)$$

Where

$$\forall j \in [1, K]$$
$$Q(\mathrm{X}, \Omega)_{*,j} = vec(\Omega_{*,j} \times \mathrm{X}_{j,*})$$

And solution vector $\vec{D}$ will be found as NNLS solution.

$$\boxed{\vec{D} = NNLS(Q(\mathrm{X}, \Omega), \mathrm{vec}(S \times \overset{\infty}{V_{ss}} \times R^T))} \qquad (10)$$

In our optimization strategy we update $D$ twice per iteration, after each update for variables $X$ and $\Omega$.

This means that in practice we will deal with different values of these variables.

For given iteration $i$

- The first update involves calculations with already updated value of $X_{i+1}$ and old value of $\Omega_i$

$$\vec{D} = NNLS(Q(\mathrm{X}_{i+1}, \Omega_{\mathrm{i}}), \mathrm{vec}(S \times \overset{\infty}{V_{ss}} \times R^T))$$

- For the second update both variables have already been updated to $X_{i+1}$ and $\Omega_{i+1}$

$$\vec{D} = NNLS(Q(X_{\mathrm{i+1}}, \Omega_{\mathrm{i+1}}), \mathrm{vec}(S \times \overset{\infty}{V}_{ss} \times R^T))$$

### Optional elimination of matrix D during optimization.

While training process defined above is highly interpretable, since $X, D$ and $\Omega$ represent actual simplex coordinates in projected space and normalization matrix, this procedure is redundant and could be simplified computationally.

Note that $D$ by its definition should be a constant which represent the relation between two simplexes (coordinate matrices $X$ and $\Omega$). This means that we can obtain this matrix whenever we want from $X$ and $\Omega$. Therefore, it is worth it to eliminate this matrix from the entire optimization problem.

Note that matrix $S \times \overset{\infty}{V}_{ss} \times R^T$ is a precomputed matrix of singular values for matrix $\overset{\infty}{V}_{ss}$.

$$\overset{\infty}{V}_{ss} = S^T \times \Sigma_{ss} \times R \Longrightarrow S \times \overset{\infty}{V}_{ss} \times R^T = \underbrace{S \times S^T}_{\mathbb{I}_{K\times K}} \times \Sigma_{ss} \times \underbrace{R \times R^T}_{\mathbb{I}_{K\times K}} = \Sigma_{ss}$$

Which means that optimization equation could be rewritten.

$$S \times \overset{\infty}{V}_{ss} \times R^T = \Sigma_{ss} = \Omega \times D \times X$$

Or in extended from

$$\begin{bmatrix} \sigma_1 & 0 & \dots & 0 \\ 0 & \sigma_2 & \dots & 0 \\ \vdots & \vdots & \ddots & \vdots \\ 0 & 0 & \dots & \sigma_k \end{bmatrix} = \begin{bmatrix} \frac{1}{\sqrt{M}} & \dots & \frac{1}{\sqrt{M}} \\ \omega_{2,1} & \dots & \omega_{2,K} \\ \vdots & \ddots & \vdots \\ \omega_{K,1} & \dots & \omega_{K,K} \end{bmatrix} \times \begin{bmatrix} d_1 & 0 & \dots & 0 \\ 0 & d_2 & \dots & 0 \\ \vdots & \vdots & \ddots & \vdots \\ 0 & 0 & \dots & d_k \end{bmatrix} \times \begin{bmatrix} \frac{1}{\sqrt{N}} & x_{1,2} & \dots & x_{1,K} \\ \vdots & \vdots & \ddots & \vdots \\ \frac{1}{\sqrt{N}} & x_{K,2} & \dots & x_{K,K} \end{bmatrix}$$

We then can represent matrix $D$ as a multiplication of two matrices.

$$D = \begin{bmatrix} \sqrt{d_1} & 0 & \dots & 0 \\ 0 & \sqrt{d_2} & \dots & 0 \\ \vdots & \vdots & \ddots & \vdots \\ 0 & 0 & \dots & \sqrt{d_k} \end{bmatrix} \begin{bmatrix} \sqrt{d_1} & 0 & \dots & 0 \\ 0 & \sqrt{d_2} & \dots & 0 \\ \vdots & \vdots & \ddots & \vdots \\ 0 & 0 & \dots & \sqrt{d_k} \end{bmatrix} = \sqrt{D} \times \sqrt{D}$$

And then define new coordinate matrices.

$$\tilde{X} = \sqrt{D} \times X$$

$$\tilde{\Omega} = \Omega \times \sqrt{D}$$

This substitution allows us to rewrite original equation in a simplified manner.

$$\Sigma_{ss} = \tilde{\Omega} \times \tilde{X}$$

Note that $\Sigma_{ss}$ is a constant as well and we can further separate each $\sigma_i$ into $\sqrt{\sigma_i} * \sqrt{\sigma_i}$

$$\begin{bmatrix} \sigma_1 & \dots & 0 \\ \vdots & \ddots & \vdots \\ 0 & \dots & \sigma_k \end{bmatrix} = \begin{bmatrix} \sqrt{\sigma_1} & \dots & 0 \\ \vdots & \ddots & \vdots \\ 0 & \dots & \sqrt{\sigma_K} \end{bmatrix} \times \begin{bmatrix} 1 & \dots & 0 \\ \vdots & \ddots & \vdots \\ 0 & \dots & 1 \end{bmatrix} \times \begin{bmatrix} \sqrt{\sigma_1} & \dots & 0 \\ \vdots & \ddots & \vdots \\ 0 & \dots & \sqrt{\sigma_K} \end{bmatrix} = \tilde{X} \times \tilde{\Omega}$$

Then multiplying this equation by two inverse matrices we can move all $\sigma_i$ to a right part.

$$\begin{bmatrix} 1 & \dots & 0 \\ \vdots & \ddots & \vdots \\ 0 & \dots & 1 \end{bmatrix} = \begin{bmatrix} \frac{1}{\sqrt{\sigma_1}} & \dots & 0 \\ \vdots & \ddots & \vdots \\ 0 & \dots & \frac{1}{\sqrt{\sigma_K}} \end{bmatrix} \times \tilde{\Omega} \times \tilde{X} \times \begin{bmatrix} \frac{1}{\sqrt{\sigma_1}} & \dots & 0 \\ \vdots & \ddots & \vdots \\ 0 & \dots & \frac{1}{\sqrt{\sigma_K}} \end{bmatrix}$$

If we again define new coordinate matrices.

$$\tilde{\tilde{X}} = \tilde{X} \times \begin{bmatrix} \frac{1}{\sqrt{\sigma_1}} & \dots & 0 \\ \vdots & \ddots & \vdots \\ 0 & \dots & \frac{1}{\sqrt{\sigma_K}} \end{bmatrix}$$

$$\tilde{\tilde{\Omega}} = \begin{bmatrix} \frac{1}{\sqrt{\sigma_1}} & \dots & 0 \\ \vdots & \ddots & \vdots \\ 0 & \dots & \frac{1}{\sqrt{\sigma_K}} \end{bmatrix} \times \tilde{\Omega}$$

We can get even more simplified equation to optimize:

$$\boxed{\Sigma_{ss} = \Omega \times D \times X \iff \mathbb{I}_{K\times K} = \tilde{\tilde{\Omega}} \times \tilde{\tilde{X}}}$$

Or in extended form

$$\begin{bmatrix} 1 & \dots & 0 \\ \vdots & \ddots & \vdots \\ 0 & \dots & 1 \end{bmatrix} = \begin{bmatrix} \frac{\sqrt{d_1}}{\sqrt{\sigma_1}\sqrt{M}} & \dots & \frac{\sqrt{d_k}}{\sqrt{\sigma_1}\sqrt{M}} \\ \frac{\sqrt{d_1}\omega_{2,1}}{\sqrt{\sigma_2}} & \dots & \frac{\sqrt{d_K}\omega_{2,K}}{\sqrt{\sigma_2}} \\ \vdots & \ddots & \vdots \\ \frac{\sqrt{d_1}\omega_{K,1}}{\sqrt{\sigma_K}} & \dots & \frac{\sqrt{d_K}\omega_{K,K}}{\sqrt{\sigma_K}} \end{bmatrix} \times \begin{bmatrix} \frac{\sqrt{d_1}}{\sqrt{\sigma_1}\sqrt{N}} & \frac{\sqrt{d_1}x_{1,2}}{\sqrt{\sigma_2}} & \dots & \frac{\sqrt{d_1}x_{1,K}}{\sqrt{\sigma_K}} \\ \vdots & \vdots & \ddots & \vdots \\ \frac{\sqrt{d_K}}{\sqrt{\sigma_1}\sqrt{N}} & \frac{\sqrt{d_K}x_{K,2}}{\sqrt{\sigma_2}} & \dots & \frac{\sqrt{d_K}x_{K,K}}{\sqrt{\sigma_K}} \end{bmatrix}$$

Note that for this equation it is important to keep first column of the $\tilde{\tilde{X}}$ and first row of the $\tilde{\tilde{\Omega}}$ related. Specifically:

$$\sqrt{N}\tilde{\tilde{\mathrm{X}}}_{*,1} = \sqrt{M}\tilde{\tilde{\Omega}}_{1,*}$$

Resulting in a optimization problem with one more additional constraint

$$\min_{\substack{\tilde{\tilde{X}} \\ \tilde{\tilde{\Omega}}}} \left\| \mathbb{I}_{K\times K} - \tilde{\tilde{\Omega}} \times \tilde{\tilde{X}} \right\| \quad s.t. \quad \begin{array}{l} \overset{\infty}{W}_{gs} = S^T \times \sqrt{\Sigma_{ss}} \times \tilde{\tilde{\Omega}} \times \sqrt{D} > 0 \\ \overset{\infty}{H}_{ss} = \sqrt{D} \times \tilde{\tilde{X}} \times \sqrt{\Sigma_{ss}} \times R > 0 \\ \sqrt{N}\tilde{\tilde{\mathrm{X}}}_{*,1} = \sqrt{M}\tilde{\tilde{\Omega}}_{1,*} \end{array}$$

Note that positivity constraints for $\overset{\infty}{W}_{gs}$ and $\overset{\infty}{H}_{ss}$ does not change with or without $\sqrt{D}$ due to positivity of matrix $D$

$$S^T \times \sqrt{\Sigma_{ss}} \times \Omega \times \sqrt{D} > 0 \iff S^T \times \sqrt{\Sigma_{ss}} \times \Omega > 0$$

$$\sqrt{D} \times X \times \sqrt{\Sigma_{ss}} \times R > 0 \iff X \times \sqrt{\Sigma_{ss}} \times R > 0$$

This fact allows us to simplify these constraints, eliminating D completely from the problem

$$\min_{\substack{\tilde{\tilde{X}} \\ \tilde{\tilde{\Omega}}}} \left\| \mathbb{I}_{K\times K} - \tilde{\tilde{\Omega}} \times \tilde{\tilde{X}} \right\| \quad s.t. \quad \begin{array}{l} S^T \times \sqrt{\Sigma_{ss}} \times \tilde{\tilde{\Omega}} > 0 \\ \tilde{\tilde{X}} \times \sqrt{\Sigma_{ss}} \times R > 0 \\ \sqrt{N}\tilde{\tilde{X}}_{*,1} = \sqrt{M}\tilde{\tilde{\Omega}}_{1,*} \end{array}$$

Note for this substitution the cost function and update rules should be changed according to new constraints.

### *The new cost function*

New const function for optimization could be defined in a similar way as we defined it above.

$$C(X,\Omega) = \left\| \mathbb{I}_{K\times K} - \tilde{\tilde{\Omega}} \times \tilde{\tilde{X}} \right\|_F^2 + \beta * \left\| S^T \times \sqrt{\Sigma_{ss}} \times \tilde{\tilde{\Omega}} \right\|_h + \lambda * \left\| \tilde{\tilde{X}} \times \sqrt{\Sigma_{ss}} \times R \right\|_h$$

This cost function should be optimized in a scenario, when $\sqrt{N}\tilde{\tilde{X}}_{*,1} = \sqrt{M}\tilde{\tilde{\Omega}}_{1,*}$

### *The new training process and update rules*

Elimination of $D$ allows us to perform conventional gradient descent with simultaneous parameters updates.

Where $\tilde{\tilde{X}}$ is updated according to gradient of defined cost function with respect to $\tilde{\tilde{X}}$

$$\nabla F_{\tilde{\tilde{X}}_i} = -2 * \tilde{\tilde{\Omega}}_i^{\mathrm{T}} \times \left( \mathbb{I}_{K\times K} - \tilde{\tilde{\Omega}}_i \times \tilde{\tilde{X}}_i \right) + \lambda * \frac{\partial \left\| \tilde{\tilde{X}} \times \sqrt{\Sigma_{ss}} \times R \right\|_h}{\partial \tilde{\tilde{X}}}$$

Where $\tilde{\tilde{\Omega}}$ is updated according to gradient of defined cost function with respect to $\tilde{\tilde{\Omega}}$

$$\nabla F_{\Omega_i} = -2 * \left( \mathbb{I}_{K\times K} - \tilde{\tilde{\Omega}}_i \times \tilde{\tilde{X}}_i \right) \times \tilde{\tilde{X}}_i^T + \beta * \frac{\partial \left\| S^T \times \sqrt{\Sigma_{ss}} \times \tilde{\tilde{\Omega}} \right\|_h}{\partial \tilde{\tilde{\Omega}}}$$

Derivatives for hinge terms could be calculated in a same way as previously

$$\forall l \in [1,K] \left( \frac{\left\| \tilde{\tilde{X}} \times \sqrt{\Sigma_{ss}} \times R \right\|_h}{\partial \tilde{\tilde{X}}_{l,*}} \right)^T = \sum_{j=1}^{N} \begin{cases} -\overrightarrow{\left(\sqrt{\Sigma_{ss}} \times R\right)_{*,j}}, & if \left( \tilde{\tilde{X}} \times \sqrt{\Sigma_{ss}} \times R \right)_{l,j} < 0 \\ 0, & if \left( \tilde{\tilde{X}} \times \sqrt{\Sigma_{ss}} \times R \right)_{l,j} \geq 0 \end{cases}$$

$$\forall l \in [1,K] \left( \frac{\left\| S^T \times \sqrt{\Sigma_{ss}} \times \tilde{\tilde{\Omega}} \right\|_h}{\partial \tilde{\tilde{\Omega}}_{*,l}} \right)^T = \sum_{j=1}^{N} \begin{cases} -\overrightarrow{S^T \times \sqrt{\Sigma_{ss}}_{*,j}}, & if \left( S^T \times \sqrt{\Sigma_{ss}} \times \tilde{\tilde{\Omega}} \right)_{l,j} < 0 \\ 0, & if \left( S^T \times \sqrt{\Sigma_{ss}} \times \tilde{\tilde{\Omega}} \right)_{l,j} \geq 0 \end{cases}$$

### *Obtain matrix D*

However, optimization result we got might not satisfy the new constraint involving relation between $\tilde{\tilde{X}}_{*,1}$ and $\tilde{\tilde{\Omega}}_{1,*}$ which we defined above. Practically could always be restored by correcting result variables.

Using our initial assumption about original variables, we always can rewrite multiplication of $\tilde{\tilde{\Omega}}$ and $\tilde{\tilde{X}}$ in a following way:

$$\mathbb{I}_{K\times K} = \tilde{\tilde{\Omega}} \times \tilde{\tilde{X}} = \sqrt{\Sigma_{ss}} \times \Omega \times \sqrt{D_\Omega} \times \sqrt{D_X} \times X \times \sqrt{\Sigma_{ss}}$$

Where $\sqrt{\Sigma_{ss}}$ is a constant matrix, and $(X, \Omega)$ is a pair of matrices we defined as coordinate matrices for simplexes.

$\sqrt{D_\Omega}$ is a matrix, which represent estimation of $\sqrt{D}$ given $\Omega$ and could be obtained from a first row of $\tilde{\tilde{\Omega}}$ since first row of $\Omega$ is predefined.

$$\left(\tilde{\tilde{\Omega}}_{1,*}\right)^T = \begin{bmatrix} \frac{\sqrt{d_1}}{\sqrt{\sigma_1}\sqrt{M}} \\ \vdots \\ \frac{\sqrt{d_k}}{\sqrt{\sigma_1}\sqrt{M}} \end{bmatrix} \Rightarrow \sqrt{D_\Omega} = \left(\tilde{\tilde{\Omega}}_{1,*}\right)^T \sqrt{M}\sqrt{\sigma_1}$$

$\sqrt{D_\mathrm{X}}$ is a matrix, which represent estimation of $\sqrt{D}$ given $X$ and could be obtained from a first column of $\tilde{\tilde{\mathrm{X}}}$ since first column of $X$ is predefined.

$$\tilde{\tilde{\mathrm{X}}}_{*,1} = \begin{bmatrix} \frac{\sqrt{d_1}}{\sqrt{\sigma_1}\sqrt{N}} \\ \vdots \\ \frac{\sqrt{d_k}}{\sqrt{\sigma_1}\sqrt{N}} \end{bmatrix} \Rightarrow \sqrt{D_\mathrm{X}} = \tilde{\tilde{\mathrm{X}}}_{*,1}\sqrt{N}\sqrt{\sigma_1}$$

According to original problem $\sqrt{D_\Omega}$ and $\sqrt{D_X}$ should give $D$ as a result but they are not equal by default. However, we can correct these matrices to be equal introducing new matrix $\sqrt{D}$ and enforcing desired property. This could be done due to properties of diagonal matrices without any harm for result cost function value.

$$\sqrt{D} = \sqrt{\sqrt{D_\Omega} \times \sqrt{D_X}}$$

Which then can be used to update values of $\tilde{\tilde{X}}$ and $\tilde{\tilde{\Omega}}$

$$\tilde{\tilde{X}}_{new} = \sqrt{D}\sqrt{D_X^{-1}}\tilde{\tilde{X}}$$

$$\tilde{\tilde{\Omega}}_{new} = \tilde{\tilde{\Omega}}\sqrt{D_\Omega^{-1}}\sqrt{D}$$

Note that this correction does not change the value of the cost function

$$\tilde{\tilde{X}}', \tilde{\tilde{\Omega}}' = \underset{\substack{\tilde{\tilde{X}} \\ \tilde{\tilde{\Omega}}}}{\operatorname{argmin}} \left\| \mathbb{I}_{K\times K} - \tilde{\tilde{\Omega}} \times \tilde{\tilde{X}} \right\|_{\substack{\infty \\ s.t.\ W_{gs} = S^T \times \sqrt{\Sigma_{ss}} \times \Omega > 0 \\ \infty \\ H_{ss} = X \times \sqrt{\Sigma_{ss}} \times R > 0}} \Leftrightarrow \tilde{\tilde{X}}', \tilde{\tilde{\Omega}}' = \underset{\substack{\tilde{\tilde{X}} \\ \tilde{\tilde{\Omega}}}}{\operatorname{argmin}} \left\| \mathbb{I}_{K\times K} - \tilde{\tilde{\Omega}} \times \sqrt{D_\Omega^{-1}}\sqrt{D}\sqrt{D}\sqrt{D_X^{-1}} \times \tilde{\tilde{X}} \right\|_{\substack{\infty \\ s.t.\ W_{gs} = S^T \times \sqrt{\Sigma_{ss}} \times \Omega > 0 \\ \infty \\ H_{ss} = X \times \sqrt{\Sigma_{ss}} \times R > 0}}$$

### Optional elimination of matrix Ω during optimization.

Note that relation deconvolution equation we got in previous section means that $\tilde{\tilde{\Omega}}$ is equal to inversed matrix $\tilde{\tilde{X}}$.

$$\mathbb{I}_{K\times K} = \tilde{\tilde{\Omega}} \times \tilde{\tilde{X}} \Leftrightarrow \tilde{\tilde{\Omega}} = \tilde{\tilde{X}}^{-1}$$

Which means that if original $\tilde{\tilde{X}}$ estimation is invertible matrix, then deconvolution term of cost function is always optimal if we consider $\tilde{\tilde{\Omega}} = \tilde{\tilde{X}}^{-1}$.

Note, this propert also reveals the connection between simplex shapes for both pairs $(\tilde{\tilde{X}}, \tilde{\tilde{\Omega}})$ and $(X, \Omega)$ (see Appendix 9b).

This substitution yields new optimization formulation which does not involve both $\tilde{\tilde{\Omega}}$ and $D$

$$\min_{\tilde{\tilde{X}}} \left\| \mathbb{I}_{K\times K} - \tilde{\tilde{X}}^{-1} \times \tilde{\tilde{X}} \right\| \quad s.t. \quad S^T \times \sqrt{\Sigma_{ss}} \times \tilde{\tilde{X}}^{-1} > 0, \quad \tilde{\tilde{X}} \times \sqrt{\Sigma_{ss}} \times R > 0, \quad \sqrt{N}\tilde{\tilde{\mathrm{X}}}_{*,1} = \sqrt{M}\tilde{\tilde{X}}^{-1}_{1,*}$$

*The new cost function*

Since we can always maintain the relation between $\tilde{\tilde{X}}^{-1}$ and $\tilde{\tilde{X}}$ , the new const function for optimization could be defined without deconvolution term.

$$C(X) = \beta * \left\| S^T \times \sqrt{\Sigma_{ss}} \times \tilde{\tilde{X}}^{-1} \right\|_h + \lambda * \left\| \tilde{\tilde{X}} \times \sqrt{\Sigma_{ss}} \times R \right\|_h$$

This cost function again should be optimized in a scenario, when $\sqrt{N}\tilde{\tilde{\mathrm{X}}}_{*,1} = \sqrt{M}\tilde{\tilde{X}}^{-1}{}_{1,*}$

*The new training process and update rules*

Training process now involves gradient descent with respect to single matrix. However derivative now involves calculation for both should be calculated with respect to both positivity terms.

For simplicity we applied for this case same computational approach as we did for three variables, considering $\tilde{\tilde{X}}^{-1}$ as separate variable while performing alternating updates in a blockwise manner.

Where $\tilde{\tilde{X}}$ is updated according to gradient of defined cost function with respect to $\tilde{\tilde{X}}$

$$\nabla F_{\tilde{\tilde{X}}_i} = \lambda * \frac{\partial \left\| \tilde{\tilde{X}}_i \times \sqrt{\Sigma_{ss}} \times R \right\|_h}{\partial \tilde{\tilde{X}}}$$

Then $\tilde{\tilde{X}}^{-1}$ is updated according to gradient of defined cost function with respect to $\tilde{\tilde{X}}^{-1}$

$$\nabla F_{\tilde{\tilde{X}}_i^{-1}} = \beta * \frac{\partial \left\| S^T \times \sqrt{\Sigma_{ss}} \times \tilde{\tilde{X}}_i^{-1} \right\|_h}{\partial \tilde{\tilde{X}}_i^{-1}}$$

### Hyperparameters selection

#### *Selection of coefficients for update steps $\chi, \phi$*

As in all machine learning tasks the purpose of learning rates is to ensure smooth convergence. By making too big steps – we might miss the optimum and might not converge.

In this section we discuss the choice of update step coefficients defined as $\chi$ (4) and $\phi$(6) previously. Which are learning rates in our problem.

$$\boxed{X_{i+1} = X_i - \chi * \nabla F_{X_i}} \quad \boxed{\Omega_{i+1} = \Omega_i - \phi * \nabla F_{\Omega_i}}$$

In practice we found that absolute values of projection coordinates for features and samples are of different scale. For GSE19830 examples showed above, gene points coordinates are in the range $[-1 * 10^{-1}, +1 * 10^{-1}]$ and sample points have coordinates in the range $[-1 * 10^{-3}, +1 * 10^{-3}]$. The question we want to answer is how to establish step size for updates of variables of different scale.

Since our simplex points are defined to be distributed around zero, we can estimate the actual limits of the simplex space through the average proximal distance to zero of a projected point. The hypersphere with a radius equal to an average Euclidean norm of projected feature vectors was chosen to be an estimation of the simplex borders.

For first space it is:

$$\tilde{R}_X = \frac{\sum_{j=1}^{M} \sqrt{\sum_{i=1}^{K} \left(\overset{\infty}{V}_{ss} \times R^T\right)^2_{j,i}}}{M}$$

For second space it is

$$\tilde{R}_\Omega = \frac{\sum_{j=1}^{N} \sqrt{\sum_{i=1}^{K} \left(S \times \overset{\infty}{V}_{ss}\right)^2_{i,j}}}{N}$$

We can then use this distance to adjust the update step to make it related to the variance of points in a search space. Generally, we want step size to be fraction of defined $\tilde{R}_X(\tilde{R}_\Omega)$ (for example 1% of $\tilde{R}_X(\tilde{R}_\Omega)$). For this we will use scaling factors $\mu$ and $\nu$. Which are typically the same.

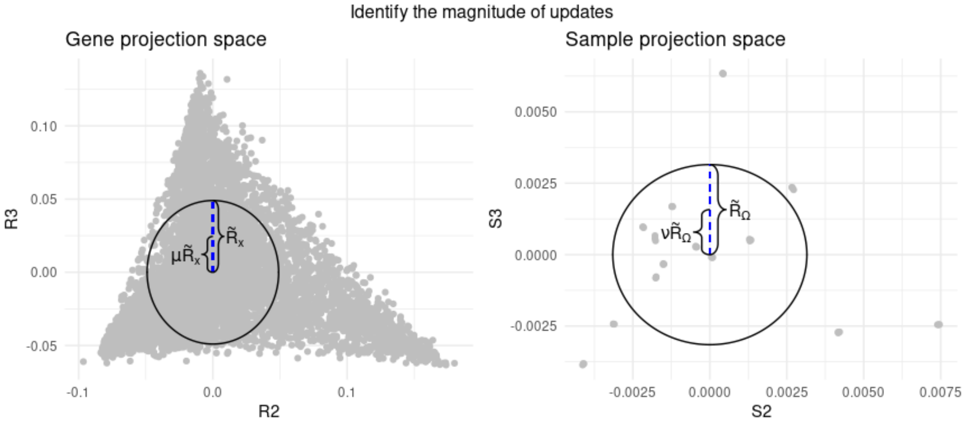

Therefore, we propose a new strategy for updates, which involves gradient clipping procedure. During update we use only unit vectors of calculated gradients: $\frac{\nabla F_{X_i}}{\|\nabla F_{X_i}\|_2}$ and $\frac{\nabla F_{\Omega_i}}{\|\nabla F_{\Omega_i}\|_2}$, which will define update directions for each vector of matrices $X$ and $\Omega$ respectively. Then we will multiply this unit vector by step size multiplier calculated using defined limits of simplex coordinates ($\tilde{R}_X(\tilde{R}_\Omega)$). The result vector will be used as a gradient vector for update.

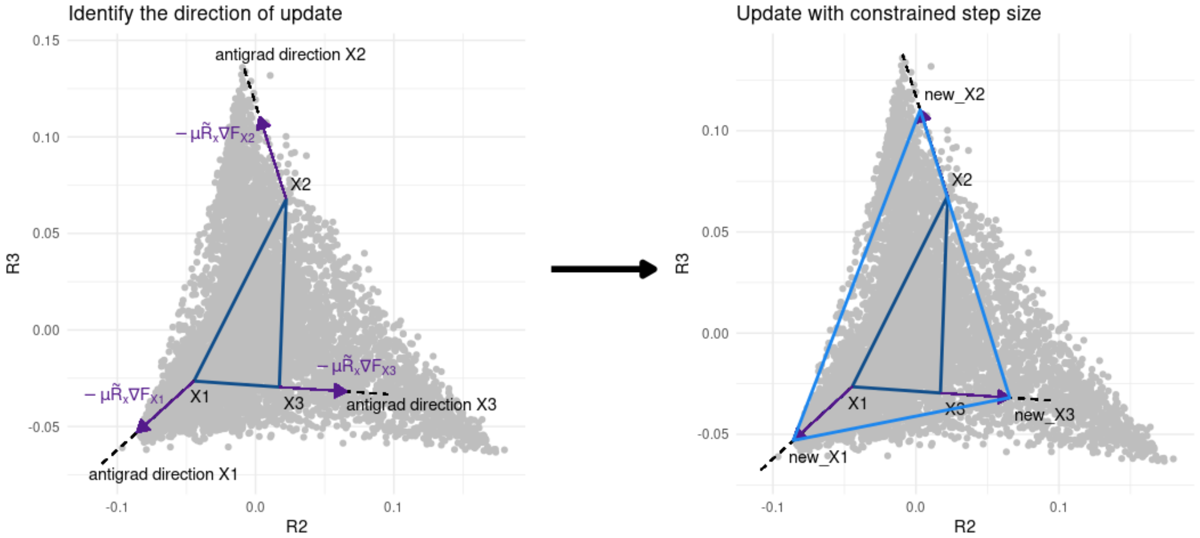


Thus, new update rule for $X$ is represented as

$$X_{i+1} = X_i - \mu * \tilde{R}_X * \frac{\nabla F_{X_i}}{\|\nabla F_{X_i}\|_2} \tag{11}$$

Where:

$\frac{\nabla F_{X_i}}{\|\nabla F_{X_i}\|_2}$ $-$ *unit direction vectors of gradient*

$\tilde{R}_X$ $-$ *average norm of projected vector*

$\mu$ $-$ *scaling constant for step (fraction of an average norm)*

Similarly, the new update rule for $\Omega$

$$\Omega_{i+1} = \Omega_i - \nu * \tilde{R}_\Omega * \frac{\nabla F_{\Omega_i}}{\|\nabla F_{\Omega_i}\|_2} \tag{12}$$

Where:

$\frac{\nabla F_{\Omega_i}}{\|\nabla F_{\Omega_i}\|_2}$ $-$ *unit direction vectors of gradient*

$\tilde{R}_\Omega$ $-$ *average norm of projected point vector*

$\nu$ $-$ *scaling constant for step (fraction of an average norm)*

If we will use the previous notation of update rules, $\chi(\phi)$ will now become the adaptive learning rate, which is dependent on the norm of gradient matrix $\nabla F_{X_i}(\nabla F_{\Omega_i})$

$$\boxed{\chi = \mu \frac{\tilde{R}_{\mathrm{X}}}{\left\|\nabla F_{\mathrm{X}_i}\right\|_2}} \quad \boxed{\phi = \nu \frac{\tilde{R}_{\Omega}}{\left\|\nabla F_{\Omega_i}\right\|_2}}$$

In our examples we found that good choices for $\mu$ and $\nu$ will be values in the range $[0.001, 0.01]$ (i.e. the length of the optimization step will be from 0.1% to 1% of the average norm values $\tilde{R}_{\mathrm{X}}, \tilde{R}_{\Omega}$).

*<u>Selection of penalty coefficients $\beta, \lambda$</u>*

As we already described, during the training penalty coefficients $\beta, \lambda$ should also be selected for the defined cost function (3)

$$C(X, \Omega, D) = \left\| S \times \overset{\infty}{V_{ss}} \times R^T - \Omega \times D \times X \right\|_F^2 + \beta * \|S^T \times \Omega\|_h + \lambda * \|X \times R\|_h$$

By varying penalty coefficients, we can restrict or relax negativity constraints during optimization process. Usage of two independent coefficients allows us to control negativity separately for proportions and expression in pure components.

We can plot values of each term with respect for each training step:

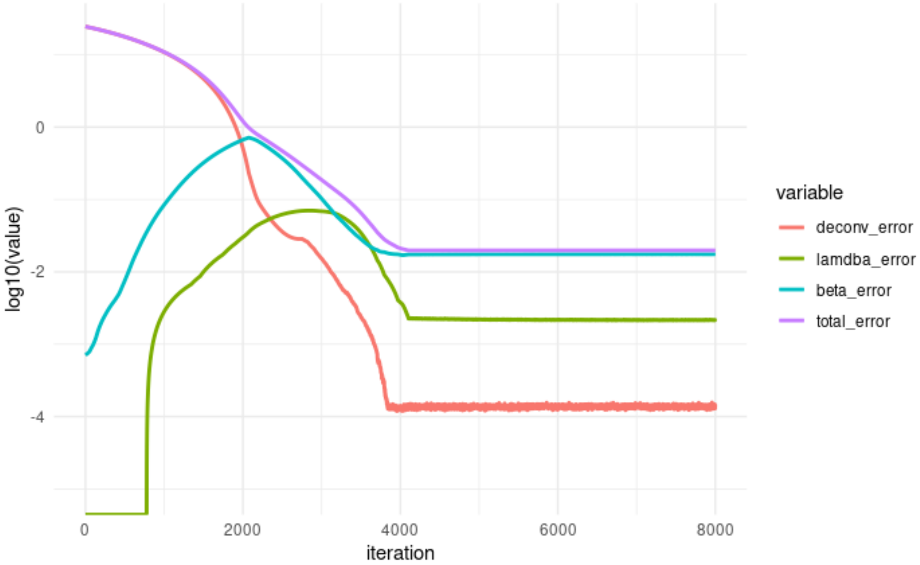


(Simulated data sd=2.5, no filtering, $\lambda = 1,\ \beta = 10,\ \mu = 0.001,\ \nu = 0.001,\ M = 10000,\ N = 40,\ K = 3$)

Where

$$deconvolution\ error = \left\| S \times \overset{\infty}{V_{ss}} \times R^T - \Omega \times D \times X \right\|_F^2$$
$$lambda\ error = \lambda * \|X \times R\|_h$$
$$beta\ error = \beta * \|S^T \times \Omega\|_h$$
$$total\ error = C(X, \Omega, D)$$

Larger values of deconvolution error correspond to how close (similar) is factorization result to input data.

For negativity errors, we cannot do such statement, since these terms represent number of negative elements and their absolute values. To monitor what caused increase of this function researcher should make two plots: the first will represent percentage of negative elements, and second will describe distribution of values for result matrices.

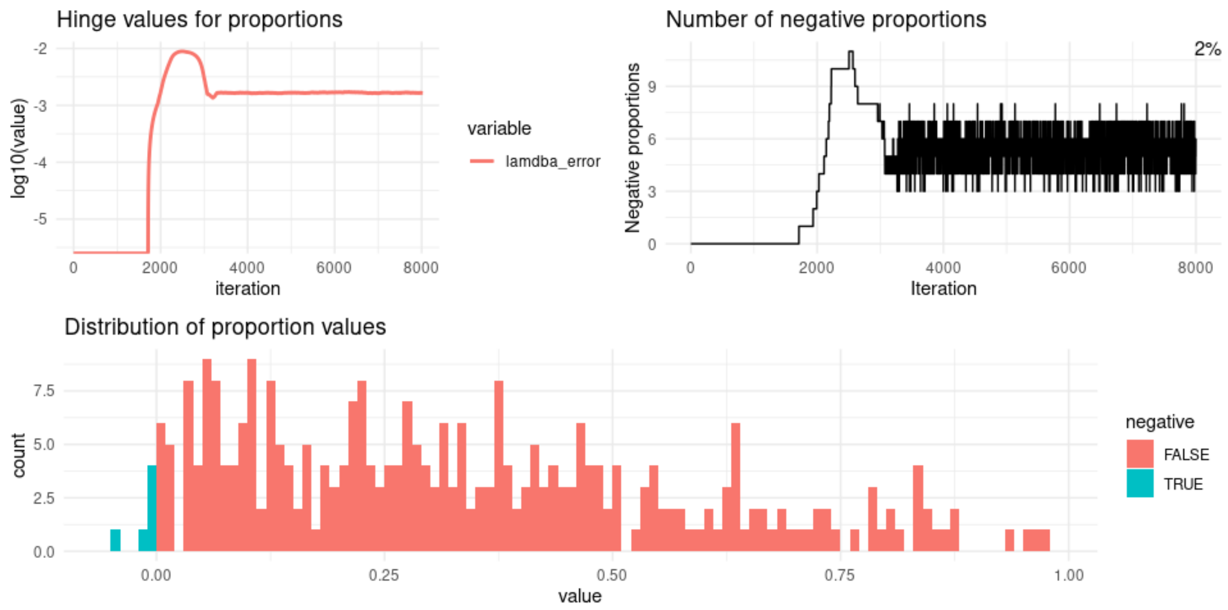


(Simulated data sd=2.5, no filtering, $\lambda = 1,\ \beta = 10,\ \mu = 0.001,\ \nu = 0.001,\ M = 10000,\ N = 40,\ K = 3$)

In general, all penalty coefficients should be picked independently based on the input data size. We can track errors during training and choose coefficients based on the behavior of the model.

The whole optimization for our task is a process of finding an equilibrium between all three terms of the cost function (3). A term with a highest value will contribute more to the next optimization step, which will lead to decreasing of this term. This will be continued until function will reach plateau, where term values are close enough to each other (in terms of gradient magnitudes), so that change of deconvolution parameters will be blocked by gradient of negativity constraints and vice versa.

In this setting the meeting point of deconvolution error and both lambda and beta errors are deciding. From this point the model will start joined optimization towards described state of equilibrium, in which optimization will be finished. This is harder the model to make significant improvements for deconvolution term in joined optimization. So, this joint problem should be understood as a slight adjustment of solution according to constraints. Chosen penalty coefficients will decide the amount of steps deconvolution error will be optimized freely.

To get some intuition about error terms we can inspect the behavior of the model in three main scenarios:

1. No negativity constrains ($\lambda = 0, \beta = 0$). For this scenario only deconvolution error will be optimized.
2. Negativity constraints are too strong. (e.g., $\lambda = 50, \beta = 50$)
3. Negativity penalty coefficients are optimal.

**Without negativity constrains.** The model will fail to find the right solution, since we will lose the important property of positivity for matrices. $\overset{\infty}{H}_{ss}$ and $\overset{\infty}{W}_{gs}$. These negative elements will make our $X$ coordinates free of actual simplex corner coordinates since any wrong basis vector coordinate assumption could be just corrected by negative element of another vector.

$$(\overset{\infty}{v}_{ss_{i,*}})^T = \begin{bmatrix} \sum_k \overset{\infty}{w}_{ss_{i,k}} * \overset{\infty}{h}_{ss_{k,1}} \\ \vdots \\ \sum_k \overset{\infty}{w}_{ss_{i,k}} * \overset{\infty}{h}_{ss_{k,N}} \end{bmatrix} = \alpha_{i,1} \begin{bmatrix} (X \times R)_{1,1} \\ \vdots \\ (X \times R)_{1,N} \end{bmatrix} + \cdots + \alpha_{i,K} \begin{bmatrix} (X \times R)_{K,1} \\ \vdots \\ (X \times R)_{K,N} \end{bmatrix}$$

$$\overset{\infty}{v}_{gs_{*,j}} = \begin{bmatrix} \sum_k \overset{\infty}{w}_{gs_{1,k}} * \overset{\infty}{h}_{gs_{k,j}} \\ \vdots \\ \sum_k \overset{\infty}{w}_{gs_{M,k}} * \overset{\infty}{h}_{gs_{k,j}} \end{bmatrix} = \alpha_{1,j} \begin{bmatrix} (S^T \times \Omega)_{1,1} \\ \vdots \\ (S^T \times \Omega)_{M,1} \end{bmatrix} + \cdots + \alpha_{K,j} \begin{bmatrix} (S^T \times \Omega)_{1,K} \\ \vdots \\ (S^T \times \Omega)_{M,K} \end{bmatrix}$$

In practice we found that solution will converge some local minimum value, where $X$ and $\Omega$ might be far from the actual corners.

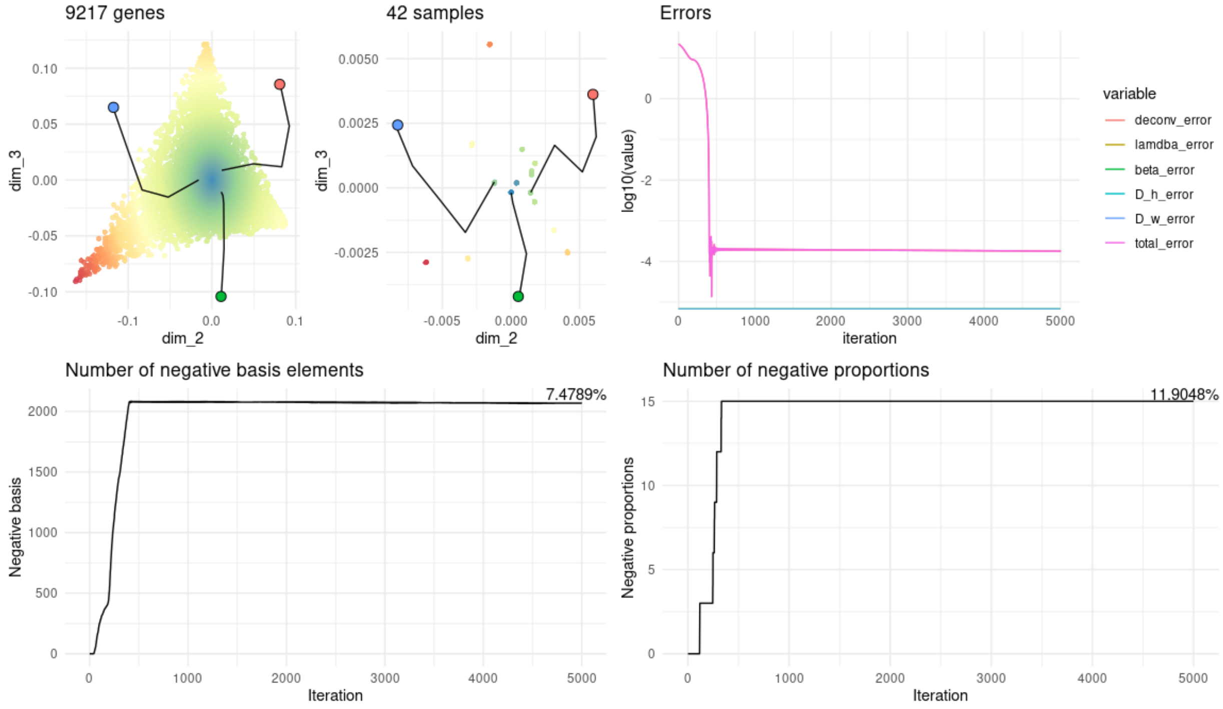


(GSE19830, $\lambda = 0,\ \beta = 0,\ \mu = 0.01,\ \nu = 0.01,\ M = 9217,\ N = 42,\ K = 3$)

**High values of $\lambda$ and $\beta$** will prevent total error from optimization. In this scenario this is too expensive for the model to leave equilibrium state, so we have no improvements at all.

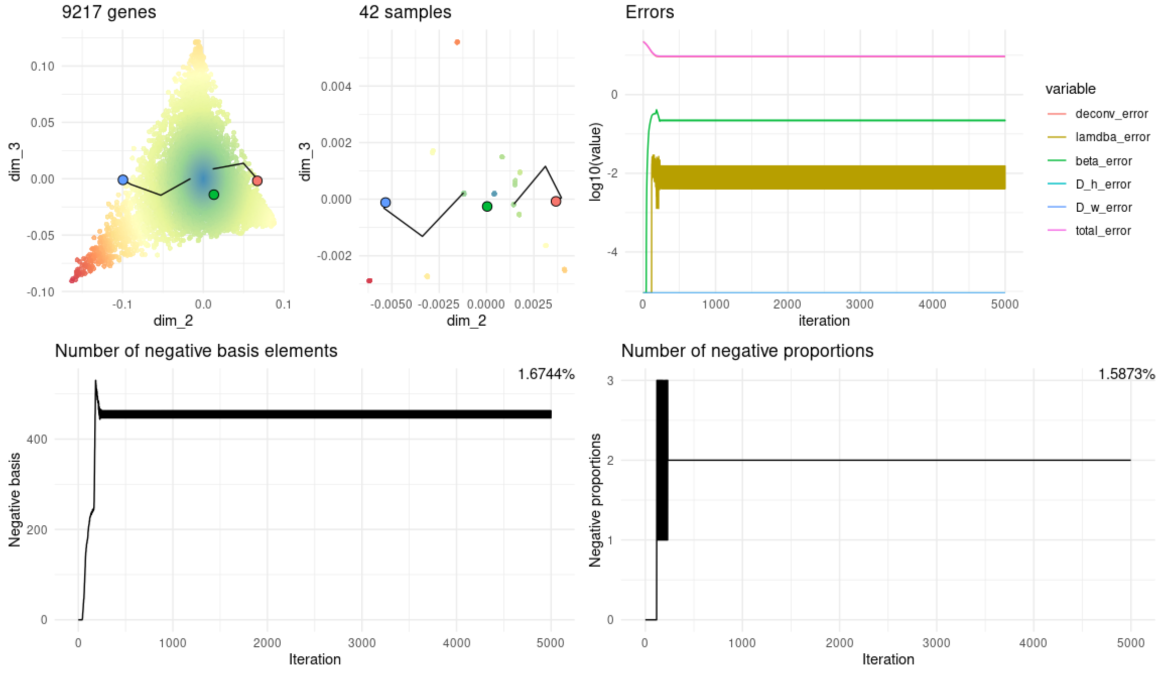


(GSE19830, $\lambda = 50,\ \beta = 50,\ \mu = 0.01,\ \nu = 0.01,\ M = 9217,\ N = 42,\ K = 3$)

**For optimal parameters** deconvolution error will be optimized until it's absolute value will reach magnitude of negativity constraints values (which usually is increasing over training steps for unconstraint case). Then the solution will be pushed towards NMF solution, which will penalize $X$ rows and $\Omega$ columns from being too far from corresponding simplex corners. The meeting points of deconvolution error and both lambda and beta errors are deciding. Starting from this time timestamp the model will force matrices $\overset{\infty}{H}_{ss}$ and $\overset{\infty}{W}_{gs}$ to be positive, penalizing $X$ and $\Omega$ from being far from corners.

The schema of training, we found useful, is to start with situation, similar to unconstrained one. This is done by setting $\lambda$ and $\beta$ parameters as small as possible. After this we gradually increase parameters to penalize model for having negative elements more and more until we reached desired negative percentage.

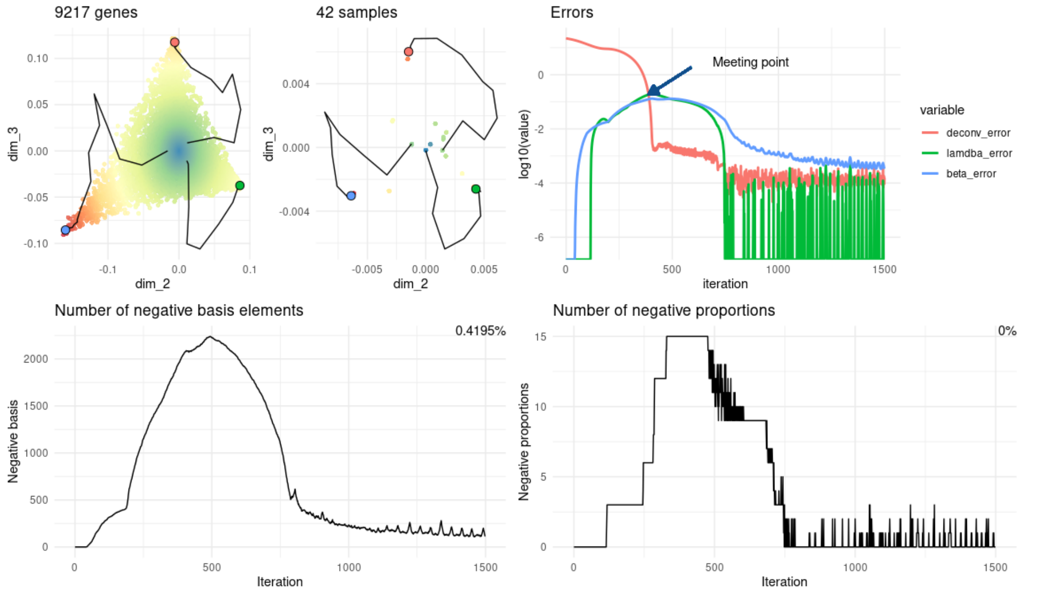


(GSE19830, $\lambda = 1,\ \beta = 1,\ \mu = 0.01,\ \nu = 0.01,\ M = 9217,\ N = 42,\ K = 3$)

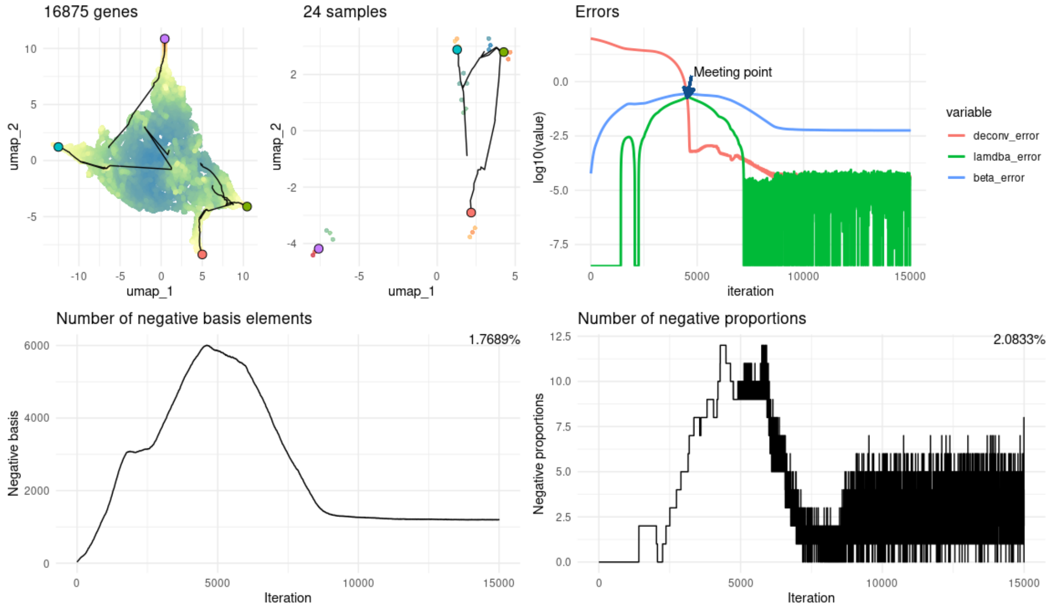


(GSE11058, $\lambda = 1,\ \beta = 1,\ \mu = 0.001,\ \nu = 0.001,\ M = 16875,\ N = 24,\ K = 4$)

### Initialization strategies

The deconvolution problem as formulated in (3) belongs to the non-convex class of functions. That means we are not guaranteed to find a global minimum of function (3), but instead we are looking for some local minimum. Knowing this we need to have different initialization strategies that can produce all possible solutions for the deconvolution problem. In this section, we will describe two approaches to finding initialization points.

By initialization we mean the selection initial values for $X$ and $\Omega$ matrices before training. For each matrix we should select $K$ points from $K$ dimensional projected space of features and samples respectively to represent initial corners of simplexes.

In general case we don't have to select $D$ sine it could be derived from known $X$ and $\Omega$ according to (10):

$$\boxed{\vec{D} = NNLS(Q(\mathrm{X}, \Omega), \mathrm{vec}(S \times \overset{\infty}{V_{ss}} \times R^T))}$$

*Single-side initialization using vertex component analysis (VCA)*

Both simplexes described in this paper are constructed to be related. To maintain this relation, we can select initial points for one simplex (features or samples) and calculate corresponding initial points for the another using deconvolution error formula (2).

$$\boxed{C(X, \Omega, D) = \left\| S \times \overset{\infty}{V_{ss}} \times R^T - \Omega \times D \times X \right\|_F^2}$$

This approach will help us to keep the coordinates in both spaces synchronized, which will give model more correct intuition about the search space.

The proposed solution for single-side initialization is:

1. Initialize matrix for features ($X$) or samples ($\Omega$) simplex with any possible procedure.
2. Calculate the values of $D$ using already known matrix. For this we can utilize the relation between normalizing matrix $D$ and matrices $X$, $\Omega$ (explained in Appendix 9a)

$$\boxed{\vec{D} = \frac{M}{N} X^{-T} \times \vec{A}} \quad (*)$$

Where $\vec{A}$ is column vector obtained from R vectors:

$$\begin{bmatrix} A_1 \\ A_2 \\ \vdots \\ A_K \end{bmatrix} = \begin{bmatrix} \sum_{j=1}^{N} r_{1,j} \\ \sum_{j=1}^{N} r_{2,j} \\ \vdots \\ \sum_{j=1}^{N} r_{K,j} \end{bmatrix}$$

Or

$$\boxed{\vec{D} = \Omega^{-1} \times \vec{B}} \quad (**)$$

Where $\vec{B}$ is column vector obtained from S vectors:

$$\begin{bmatrix} A_1 \\ A_2 \\ \vdots \\ A_K \end{bmatrix} = \begin{bmatrix} \sum_{i=1}^{M} s_{1,i} \\ \sum_{i=1}^{M} s_{2,i} \\ \vdots \\ \sum_{i=1}^{M} s_{K,i} \end{bmatrix}$$

3. Calculate approximation for remaining matrix $\Omega(X)$ which minimizes deconvolution error.

$$\Omega = S \times \overset{\infty}{V}_{ss} \times R^T \times \left(diag\left(\vec{D}\right) \times X\right)^{-1}$$

Or

$$X = \left(\Omega \times diag\left(\vec{D}\right)\right)^{-1} \times S \times \overset{\infty}{V}_{ss} \times R^T$$

An additional benefit of this initialization method is that we can use existing algorithms to select a good estimation of initial points in one space. The one example of such algorithm is vertex component analysis (Nascimento & Dias, 2005) which was used in this paper. This algorithm picks vertices of simplex that correspond to the extreme points of the projection.

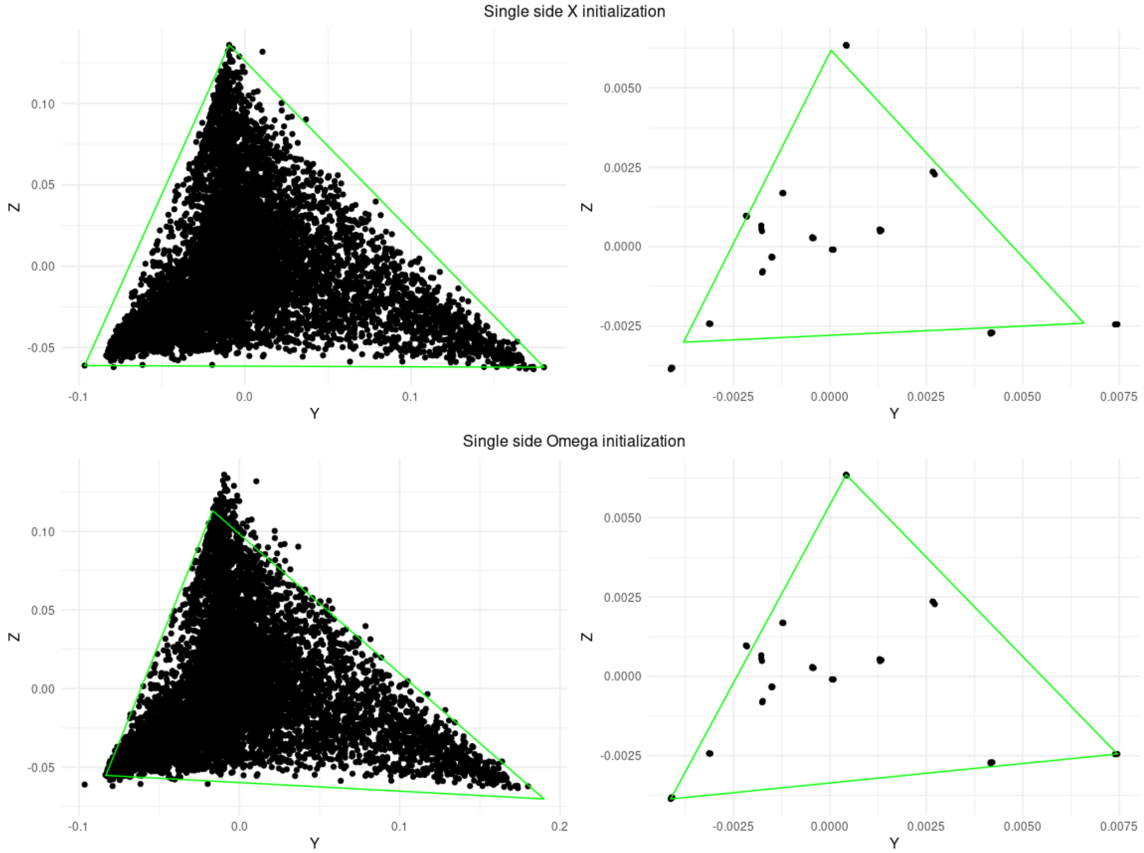


(GSE19830, $M = 10000,\ N = 42,\ K = 3$)

We have a specific pattern of the error's trajectory plots. Because we used deconvolution error to calculate the initial points for second space the value of this error will be minimal. But we cannot guarantee that any of the initial estimations will have a minimal number of negative elements in proportions and expression of pure components. Therefore, lambda and beta errors could be higher at the beginning.

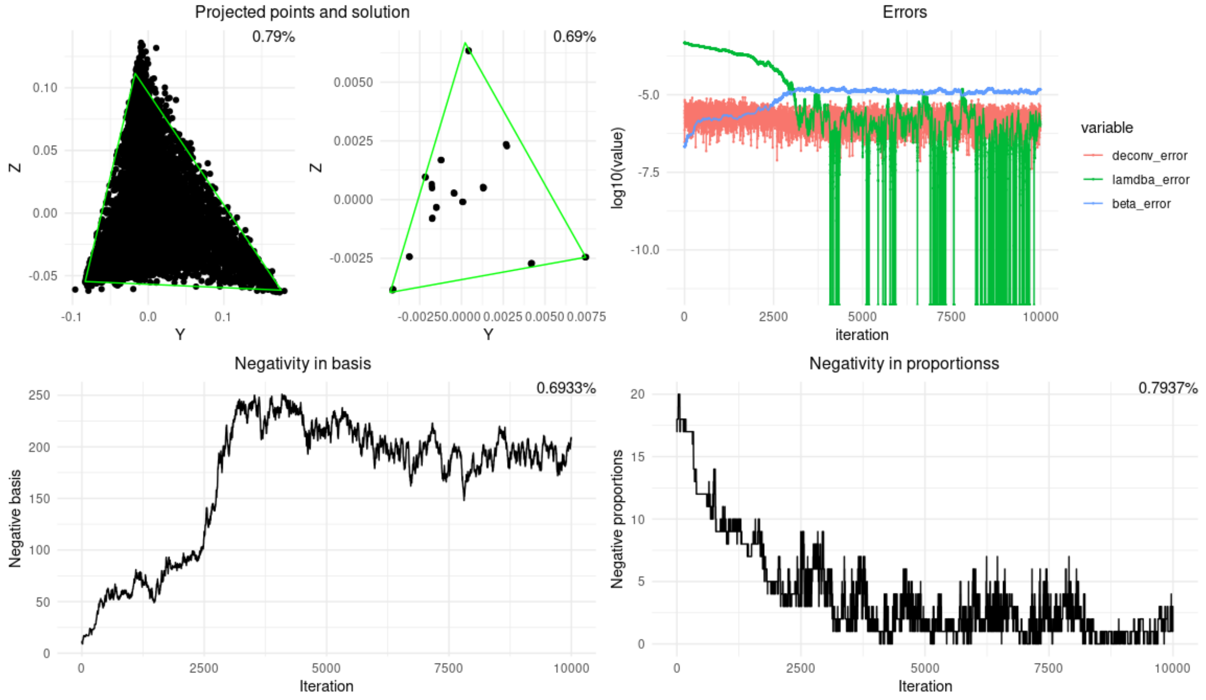


(GSE19830, $\lambda = 0.01,\ \beta = 0.01,\ \mu = 0.001,\ \nu = 0.001,\ M = 10000,\ N = 42,\ K = 3$)

### *Random centered initialization*

Another and the simplest approach is to take $K$ random points from each projected space respectively. The good choice will be just project $K$ random features into new space.

$$\boxed{X_{init} = \begin{bmatrix} \left(\overset{\infty}{V}_{ss}\right)_{i_1,*} \times R^T \\ \vdots \\ \left(\overset{\infty}{V}_{ss}\right)_{i_K,*} \times R^T \end{bmatrix}}$$

Where

$$i_1, \dots, i_K \in [1, M] - random\ row\ indexes\ from\ \overset{\infty}{V}_{ss}\ (random\ genes)$$

Moreover, we know that $\vec{R}_1^{\,T} = \begin{bmatrix} \frac{1}{\sqrt{N}} & \dots & \frac{1}{\sqrt{N}} \end{bmatrix}$ and $\overset{\infty}{H}_{ss}$ is row normalized.

Since

$$X = \overset{\infty}{H}_{ss} \times R^T$$

Then first column of $X$ could be defined directly.

$$X_{*,1} = \begin{bmatrix} \sum_{i=1}^{N} \overset{\infty}{h}_{ss_{1,i}} r_{1,i} \\ \vdots \\ \sum_{i=1}^{N} \overset{\infty}{h}_{ss_{K,i}} r_{1,i} \end{bmatrix} = \begin{bmatrix} \frac{1}{\sqrt{N}} \sum_{i=1}^{N} \overset{\infty}{h}_{ss_{1,i}} \\ \vdots \\ \frac{1}{\sqrt{N}} \sum_{i=1}^{N} \overset{\infty}{h}_{ss_{K,i}} \end{bmatrix} = \begin{bmatrix} \frac{1}{\sqrt{N}} \\ \vdots \\ \frac{1}{\sqrt{N}} \end{bmatrix}$$

For matrix Ω initialization could be done in similar way.

$$\boxed{\Omega_{init} = \begin{bmatrix} S \times \left(\overset{\infty}{V_{gs}}\right)_{*,j_1} \\ \vdots \\ S \times \left(\overset{\infty}{V_{gs}}\right)_{*,j_K} \end{bmatrix}}$$

$$j_1, \dots, j_K \in [1, N] - random\ column\ indexes\ from\ \overset{\infty}{V_{gs}}\ (random\ samples)$$

Moreover, we know that $\vec{S_1}^T = \begin{bmatrix} \frac{1}{\sqrt{M}} & \dots & \frac{1}{\sqrt{M}} \end{bmatrix}$ and $\overset{\infty}{W_{gs}}$ is column normalized.

Since

$$\Omega = S \times \overset{\infty}{W_{gs}}$$

Then first column of Ω will be defined directly.

$$\Omega = \begin{bmatrix} \sum_{j=1}^{M} \overset{\infty}{w_{gs_{j,1}}} s_{1,j} \\ \vdots \\ \sum_{j=1}^{M} \overset{\infty}{w_{gs_{j,K}}} s_{1,j} \end{bmatrix} = \begin{bmatrix} \frac{1}{\sqrt{M}} \sum_{j=1}^{M} \overset{\infty}{w_{gs_{j,1}}} \\ \vdots \\ \frac{1}{\sqrt{M}} \sum_{j=1}^{M} \overset{\infty}{w_{gs_{j,K}}} \end{bmatrix} = \begin{bmatrix} \frac{1}{\sqrt{M}} \\ \vdots \\ \frac{1}{\sqrt{M}} \end{bmatrix}$$

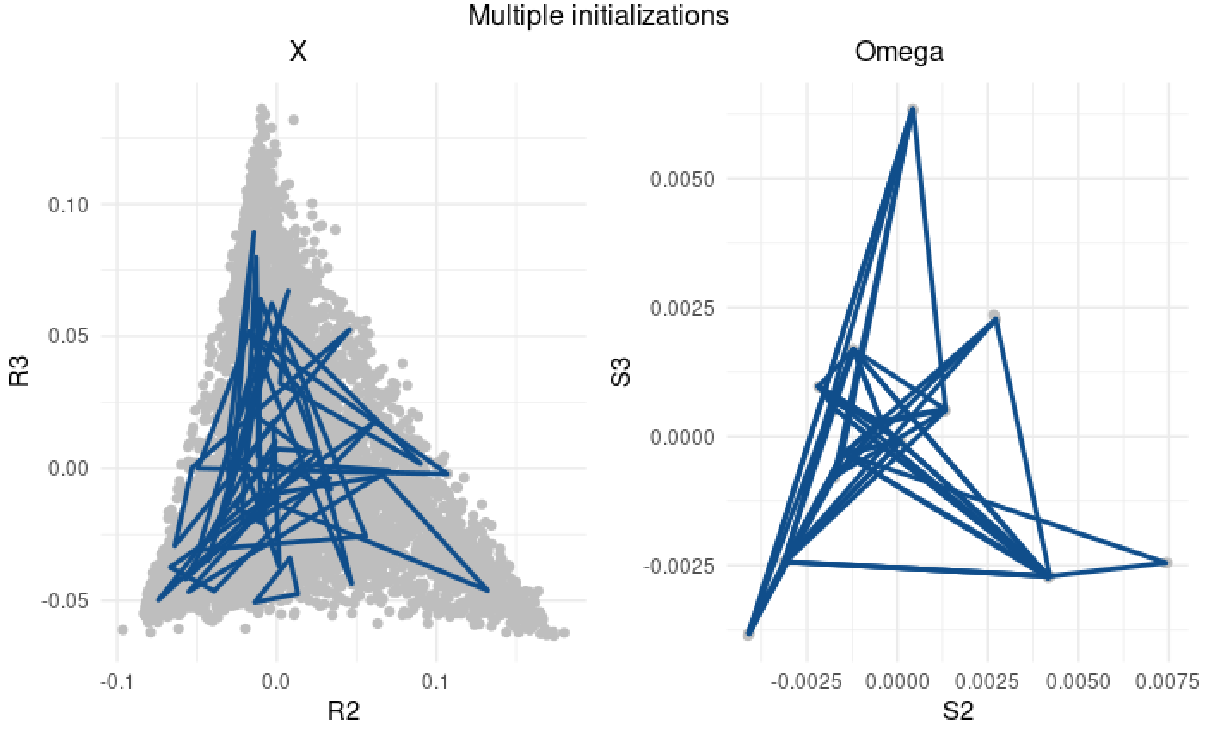


In practice points in the projected space have the center of mass at the beginning of the coordinates system (zero point). This is true for both simplexes. This means that initial simplex guess of the solution should contain this point.

We also observed that initialization which are do not contain zero point will lead to zeros in NNLS problem solution., which makes it impossible for the model to use all projection vectors during optimization.

$$\vec{D} = NNLS(Q(\mathrm{X},\Omega), \mathrm{vec}(S \times \overset{\infty}{V_{ss}} \times R^T))$$

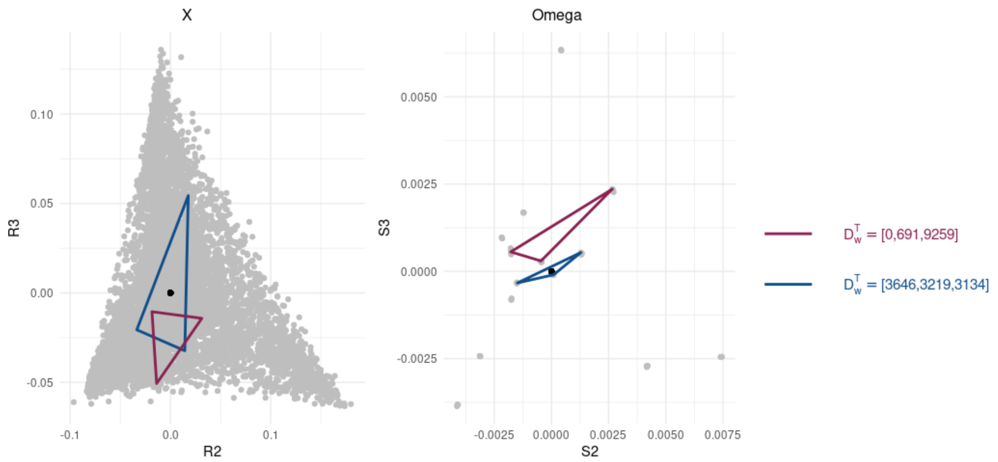


Therefore, we propose another approach to random initialization, which improves initial random guess, making it as centered around zero as possible.

The proposed solution for the most centered random initialization is to perform L (typically around 10000) random pre-initializations for both simplexes in samples and features projected spaces.

After that we select that initial values which will have center of mass for points closest to zero:

$$X_{init} = \underset{X_1, X_2, \ldots, X_L}{argmin} \sqrt{\sum_{i=1}^{K} \left( \frac{\sum_{j=1}^{K} x_{j,i}}{K} \right)^2} \qquad \Omega_{init} = \underset{\Omega_1, \Omega_2, \ldots, \Omega_L}{argmin} \sqrt{\sum_{i=1}^{K} \left( \frac{\sum_{j=1}^{K} \omega_{i,j}}{K} \right)^2}$$

Like the previous initialization method, we can see the specific pattern on the trajectory plot at the first steps of the optimization process. Typically, we will start with a few negative elements in proportions and expression in pure components, which means that initial lambda and beta errors are going to be small. However, initial deconvolution error (2) will have a higher value, due to the fact that our initial points in samples and features spaces are unsynchronized. Therefore, during the first iterations, the objective function (3) will be optimized mostly by decreasing deconvolution error (2).

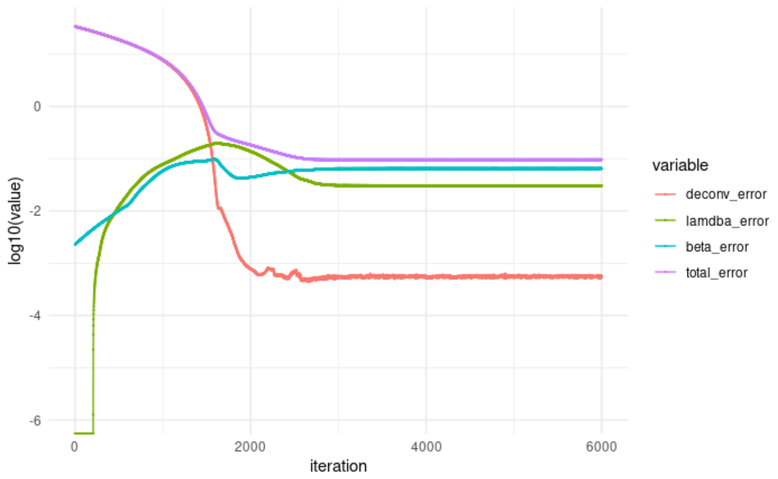


(Simulated data sd=2.5, $\lambda = 1,\ \beta = 10,\ \mu = 0.001,\ \nu = 0.001,\ M = 10000,\ N = 40,\ K = 3$)

### Appendix 10a.Relation relation between normalizing matrix $D$ and matrices $X$, $\Omega$.

*For Sinkhorn transformed matrices we have relation between normalizing matrices $\overset{\infty}{D}_w$ and $\overset{\infty}{D}_h$*

For Sinkhorn transformed matrices we have

$$\underset{col\,norm}{\overset{\overset{i}{\infty}}{H}_{gs}} \rightarrow \underset{row\,norm}{\overset{\overset{i+1}{\infty}}{H}_{ss}} \rightarrow \underset{col\,norm}{\overset{\overset{i+2}{\infty}}{H}_{gs}}$$

$$\overset{\infty}{V}_{gs} = \overset{\infty}{V}_{ss} \times D_V = \underbrace{\overset{\infty}{W}_{ss} \times \overset{\infty}{D}_w}_{\overset{\infty}{W}_{gs}} \times \underbrace{\overset{\infty}{D}_w^{-1} \times \overset{\infty}{H}_{ss} \times D_V}_{\overset{\infty}{H}_{gs}}$$

So

$$\overset{\infty}{H}_{gs} = \overset{\infty}{D}_w^{-1} \times \overset{\infty}{H}_{ss} \times D_V$$

Where

$$D_V = \begin{bmatrix} \frac{N}{M} & \dots & 0 \\ \vdots & \ddots & \vdots \\ 0 & \dots & \frac{N}{M} \end{bmatrix}$$

Note

$$\overset{\infty}{H}_{ss} = X \times R$$

So:

$$\overset{\infty}{H}_{gs} = \overset{\infty}{D}_w^{-1} \times X \times R \times D_V$$

From all above:

$$\overset{\infty}{H}_{gs} = \overset{\infty}{H}_{gs}$$

$$\overset{\infty}{D}_w^{-1} \times X \times R \times D_V = \overset{\infty}{D}_h^{-1} \times \overset{\infty}{H}_{ss}$$

$$\overset{\infty}{D}_w^{-1} \times \overset{\infty}{H}_{ss} \times D_V = \overset{\infty}{D}_h^{-1} \times \overset{\infty}{H}_{ss}$$

$$\overset{\infty}{D}_w^{-1} \times \overset{\infty}{H}_{ss} \times \frac{N}{M} = \overset{\infty}{D}_h^{-1} \times \overset{\infty}{H}_{ss}$$

Which means

$$\frac{N}{M} \times \overset{\infty}{D}_w^{-1} = \overset{\infty}{D}_h^{-1}$$

*$\vec{D}$ could be obtained through the matrix $X$*

We already showed, that for any column normalized initial matrix $H$:

$$\mathbb{I}_K^T \times H = \mathbb{I}_N^T \Rightarrow \forall m \in [1, K] \sum_{j=1}^{N} r_{m,j} = \sum_{k=1}^{K} d_{h_k} x_{k.m}$$

Where $d_{h_i}$ is a sum over row for $H$, $r_{m,j}$ elements of vectors from $R$

The same is true for vectors $R$ obtained for Sinkhorn transformed case, due to column normalization property of matrix $\overset{(2n-2)}{H}$ (if $\overset{(2n-1)}{H}$ is $\overset{\infty}{H_{ss}}$ and $\overset{(2n)}{H}$ is $\overset{\infty}{H_{gs}}$).

Let's introduce elements of $\vec{A}$

$$A_m = \sum_{j=1}^{N} r_{m,j}$$

We can do the same for every $j \in [1, K]$ so in matrix form this equation will be

$$\boxed{\begin{bmatrix} A_1 \\ A_2 \\ \vdots \\ A_K \end{bmatrix} = X^T \times \begin{bmatrix} d_{h_1} \\ d_{h_2} \\ \vdots \\ d_{h_k} \end{bmatrix}}$$

Which in a vector form is

$$X^T \times \vec{D}_h = \vec{A}$$

But

$$\vec{D}_h = diag\left(\overset{\infty}{D}{}_h^{-1}\right) = \frac{N}{M} diag(\overset{\infty}{D}{}_w^{-1}) = \frac{N}{M} \vec{D}_w = \frac{N}{M} \vec{D}$$

$$\boxed{X^T \times \vec{D} * \frac{N}{M} = \vec{A}} \qquad (1a)$$

Similarly, we showed, that for any row normalized initial matrix $\widetilde{W}$

$$\widetilde{W} \times \mathbb{I}_K = \mathbb{I}_M \Rightarrow \sum_{i=1}^{M} s_{m,j} = \sum_{k=1}^{K} d_{w_k} \omega_{m,k}$$

### *$\vec{D}$ could be obtained through the matrix Ω*

The same is true for vectors $S$ obtained for Sinkhorn transformed case, due to row normalization property of $\overset{\infty}{W_{ss}}$.

Let's introduce elements of $\vec{B}$

$$B_m = \sum_{i=1}^{M} s_{m,j}$$

We can do the same for every $j \in [1, K]$ so in matrix form this equation will be

$$\boxed{\begin{bmatrix} B_1 \\ B_2 \\ \vdots \\ B_K \end{bmatrix} = \Omega \times \begin{bmatrix} d_{w_1} \\ d_{w_2} \\ \vdots \\ d_{w_k} \end{bmatrix}}$$

Which in a vector form is

$$\boxed{\Omega \times \vec{D} = \vec{B}} \tag{2a}$$

Then it is possible to use defined equations (1a) and (2a) to obtain equations for a vector $\vec{D}$ ((*) and (**)).

$$\boxed{\vec{D} = \frac{M}{N} X^{-T} \times \vec{A}} \quad \boxed{\vec{D} = \Omega^{-1} \times \vec{B}}$$

### Appendix 10b.Geometrical relation between solution coordinates $X$ and $\Omega$.

#### *Geometrical relation for simplexes $\tilde{\tilde{X}}$ and $\tilde{\tilde{\Omega}}$*

$X$ are simplex vertices in K dimensional space of vectors $R^T$. Then a volume of the entire K dimensional simplex (including 0 point) is proportional to determinant of the $X$

$$V_{\tilde{\tilde{X}}_K} = \frac{1}{K!}\det \tilde{\tilde{X}}^T = \frac{1}{K!}\det \tilde{\tilde{X}}$$

$$V_{\tilde{\tilde{\Omega}}_K} = \frac{1}{K!}\det \tilde{\tilde{\Omega}}^T = \frac{1}{K!}\det \tilde{\tilde{\Omega}}$$

But $\det \tilde{\tilde{X}} = \frac{1}{\det \tilde{\tilde{\Omega}}}$

Then

$$V_{\tilde{\tilde{X}}_K} = \frac{1}{K!}\det \tilde{\tilde{X}} = \frac{1}{K!\det \tilde{\tilde{\Omega}}} = \frac{1}{K!\,K!\,V_{\tilde{\tilde{\Omega}}_K}} \Rightarrow V_{\tilde{\tilde{X}}_K} = \frac{1}{K!^2\,V_{\tilde{\tilde{\Omega}}_K}}$$

#### *Geometrical relation for simplexes $X$ and $\Omega$*

$X$ are simplex vertices in K dimensional space of vectors $R^T$. Then a volume of the entire K dimensional simplex (including 0 point) is proportional to determinant of the $X$

$$V_{X_K} = \frac{1}{K!}\det(X^T) = \frac{1}{K!}\det(X)$$

But we know the relation for the volumes of $n$-dimensional cone if we are going from $K$ to $K-1$

$$V_{X_K} = \frac{hV_{X_{K-1}}}{K}$$

Using this we can obtain equation for determinant of $X$

$$V_{k_X} = \frac{1}{K!}\det(X) = \frac{1}{K}hV_{X_{K-1}} = \frac{1}{K\sqrt{N}}V_{X_{K-1}} \Rightarrow \boxed{\det(\mathrm{X}) = \frac{(K-1)!}{\sqrt{N}}\,V_{X_{K-1}}}$$

For example, case X is 3-dimensional $V_{X_{K-1}}$ is an area of the triangle produced by points $X^T$

Similarly, determinant of $\Omega$ is related to a volume for a $(K-1))$ dimensional simplex $\Omega$

$$\frac{1}{K!}\det(\Omega) = \frac{1}{K\sqrt{M}}V_{\Omega_{K-1}} \Rightarrow \det(\Omega) = \frac{(K-1)!}{\sqrt{M}}\,V_{\Omega_{K-1}}$$

Determinant of $\tilde{\tilde{X}}$ could be calculated using determinant of $X$

$$\det\left(\tilde{\tilde{X}}\right) = \det\left(\sqrt{D}\times X\times\sqrt{\Sigma_{SS}^{-1}}\right) = \det\left(\sqrt{D}\right)\det(X)\det\left(\sqrt{\Sigma_{SS}^{-1}}\right) = \frac{\sqrt{d_1\ldots d_K}}{\sqrt{\sigma_1\ldots\sigma_K}}\det(X)$$

Similarly determinant of $\tilde{\tilde{\Omega}}$ could be calculated using determinant of $\Omega$

$$\det\left(\tilde{\tilde{\Omega}}\right) = \det\left(\sqrt{\Sigma_{SS}^{-1}}\times\Omega\times\sqrt{D}\right) = \det\left(\sqrt{\Sigma_{SS}^{-1}}\right)\det(\Omega)\det\left(\sqrt{D}\right) = \frac{\sqrt{d_1\ldots d_K}}{\sqrt{\sigma_1\ldots\sigma_K}}\det(\Omega)$$

Knowing the fact that $\tilde{\tilde{\Omega}}$ and $\tilde{\tilde{X}}$ are inversed we can define a relation between the volumes of the $(K-1)$ dimensional simplexes.

Determinants are proportional.

$$\det\left(\tilde{\tilde{X}}\right) = \frac{1}{\det\left(\tilde{\tilde{\Omega}}\right)} \Rightarrow$$

$$\frac{\sqrt{d_1 \dots d_3}}{\sqrt{\sigma_1 \dots \sigma_K}} \det(X) = \frac{\sqrt{\sigma_1 \dots \sigma_K}}{\sqrt{d_1 \dots d_K}} \frac{1}{\det(\Omega)} \Rightarrow$$

$$\det(X) = \frac{\sigma_1 \dots \sigma_K}{d_1 \dots d_K} \frac{1}{\det(\Omega)}$$

Let's show that volumes are proportional.

$$\det(X) = \frac{\sigma_1 \dots \sigma_K}{d_1 \dots d_K} \frac{1}{\det(\Omega)} \Rightarrow \frac{(K-1)!}{\sqrt{N}} V_{X_{K-1}} = \frac{\sigma_1 \dots \sigma_K}{d_1 \dots d_K} \frac{\sqrt{M}}{(K-1)!\, V_{\Omega_{K-1}}}$$

Resulting in a proportional relation between volumes

$$\boxed{V_{X_{K-1}} = \frac{\sqrt{M}}{\sqrt{N}} \frac{1}{(K-1)!^2} \frac{\sigma_1 \dots \sigma_K}{d_1 \dots d_K} \frac{1}{V_{\Omega_{K-1}}}}$$

### Supplementary Note 11. Reverse Sinkhorn procedure

#### **What we did so far**

*Defined NMF problem:*

$$V = W \times H$$

$$v_{i,j} \in \mathbb{R}^{M\times N}, \qquad w_{i.j} \in \mathbb{R}^{M\times K}, \qquad h_{i,j} \in \mathbb{R}^{K\times N}$$

$$\begin{aligned} &K - number\ of\ cell\ types \\ &N - number\ of\ samples \\ &M - number\ of\ genes \end{aligned}$$

Where $v_{i,j}$ describes value of $i^{th}$ feature in $j^{th}$ sample, $w_{i.k}$ describes the value of $i^{th}$ feature in $k^{th}$ main component, $h_{k,j} \geq 0$ describes contribution of $k^{th}$ component in $j^{th}$ sample.

*Introduced an iterative Sinkhorn normalization procedure factorizable matrix:*

$$V \overset{\substack{row \\ norm\ 1}}{\Longrightarrow} \underbrace{D_{v_0} \times V}_{\substack{\overset{(1)}{V} \\ simplex\ in \\ samples\ space}} \overset{\substack{column \\ norm\ 1}}{\Longrightarrow} \underbrace{D_{v_0} \times V \times D_{v_1}}_{\substack{\overset{(2)}{V} \\ simplex\ in \\ features\ space}} \overset{\substack{row \\ norm\ 2}}{\Longrightarrow} \ldots$$

$$\ldots \overset{\substack{column \\ norm\ n}}{\Longrightarrow} \underbrace{D_{v_{2n-2}} \times D_{v_{2n-4}} \ldots \times D_{v_0} \times V \times D_{v_2} \times \ldots \times D_{v_{2n-3}} \times D_{v_{2n-1}}}_{\substack{\overset{(2n)}{V} \\ simplex\ in \\ features\ space}}$$

Where odd and even elements could be represented in the following way

$$\begin{aligned} &\overset{(0)}{V} = V \\ \forall i \in [1, 3, \ldots 2n-1] \quad &\overset{(i)}{V} = D_{v_{i-1}} \times \overset{(i-1)}{V} - row\ normalized \\ \forall i \in [2, 4, \ldots 2n] \quad &\overset{(i)}{V} = \overset{(i-1)}{V} \times D_{v_{i-1}} - column\ normalized \end{aligned}$$

At the same time $W$ matrix was transformed to keep normalization properties

$$W \overset{\substack{row \\ norm\ 1}}{\Longrightarrow} \underbrace{D_{v_0} \times W \times D_{h_0}^{-1}}_{\overset{(1)}{W}} \overset{\substack{column \\ norm\ 1}}{\Longrightarrow} \underbrace{D_{v_0} \times W \times D_{h_0}^{-1} \times D_{w_1}}_{\overset{(2)}{W}} \overset{\substack{row \\ norm\ 2}}{\Longrightarrow} \ldots$$

$$\ldots \overset{\substack{column \\ norm\ n}}{\Longrightarrow} \underbrace{D_{v_{2n-2}} \times D_{v_{2n-4}} \times \ldots \times D_{v_2} \times D_{v_0} \times W \times D_{h_0}^{-1} \times D_{w_1} \times D_{h_2}^{-1} \times \ldots \times D_{h_{2n-2}}^{-1} \times D_{w_{2n-1}}}_{\overset{(2n)}{W}}$$

Same was performed for matrix $H$

$$H \overset{\substack{row \\ norm\ 1}}{\Longrightarrow} \underbrace{D_{h_0} \times H}_{\overset{(1)}{H}} \overset{\substack{column \\ norm\ 1}}{\Longrightarrow} \underbrace{D_{w_1}^{-1} \times D_{h_0} \times H \times D_{v_1}}_{\overset{(2)}{H}} \overset{\substack{row \\ norm\ 2}}{\Longrightarrow} \ldots$$

$$\dots \overset{column\ norm\ n}{\Longrightarrow} \underbrace{D^{-1}_{w_{2n-1}} \times D_{h_{2n-2}} \times D^{-1}_{w_{2n-3}} \dots \times D_{h_2} \times D^{-1}_{w_2} \times D_{h_0} \times H \times D_{v_1} \times D_{v_3} \times \dots \times D_{v_{2n-1}}}_{\overset{(2n)}{H}}$$

Where odd and even elements could be represented in the following way

$$\overset{(0)}{H} = H \qquad \overset{(0)}{W} = W$$

$$\forall i \in [1, 3, \dots 2n-1] \quad \overset{(i)}{H} = \boldsymbol{D_{h_{i-1}}} \times \overset{(i-1)}{H} \qquad \overset{(i)}{W} = \boldsymbol{D_{v_i}} \times \overset{(i-1)}{W} \times \boldsymbol{D^{-1}_{h_i}} \quad -row\ normalized$$

$$\forall i \in [2, 4, \dots 2n] \quad \overset{(i)}{H} = \boldsymbol{D^{-1}_{w_{i-1}}} \times \overset{(i-1)}{H} \times \boldsymbol{D_{v_{i-1}}} \qquad \overset{(i)}{W} = \overset{(i-1)}{W} \times \boldsymbol{D_{w_1}} \quad -column\ normalized$$

### We can rewrite this procedure into single algorithm

The whole Sinkhorn procedure for factorized matrix $V = W \times H$ could be written in one procedure.

---

Algorithm 1: Forward Sinkhorn-Knopp procedure

---

Input : $V, W, H$, n

Output : $\overset{(1)}{V}, \overset{(2)}{V}, \dots, \overset{(2n)}{V}$
$\overset{(1)}{H}, \overset{(2)}{H}, \dots, \overset{(2n)}{H}$
$\overset{(1)}{W}, \overset{(2)}{W}, \dots, \overset{(2n)}{W}$
$D_{v_0}, D_{v_1}, \dots, D_{v_{2n-1}}$
$D_{h_0}, D_{h_2}, \dots, D_{h_{2n-2}}$
$D_{w_1}, D_{w_3}, \dots, D_{w_{2n-1}}$

---

$\overset{(0)}{V} = V$

$\overset{(0)}{W} = W$

$\overset{(0)}{H} = H$

For $i$ in $[1,3, \dots 2n-1]$:

$\overset{(i)}{V} = D_{v_{i-1}} \times \overset{(i-1)}{V}$

$\overset{(i)}{H} = D_{h_{i-1}} \times \overset{(i-1)}{H}$

$\overset{(i)}{W} = D_{v_{i-1}} \times \overset{(i-1)}{W} \times D^{-1}_{h_{i-1}}$

$\overset{(i+1)}{V} = \overset{(i)}{V} \times D_{v_i}$

$\overset{(i+1)}{W} = \overset{(i)}{W} \times D_{w_i}$

$\overset{(i+1)}{H} = D^{-1}_{w_i} \times \overset{(i)}{H} \times D_{v_i}$

---

Where:

$$\forall i \in [0, 2n]\ V_i = W_i \times H_i$$

$$D_{v_i} - row\ normalizing\ matrix\ for \overset{(i)}{V}, i \in [0, 2, \dots, 2n-2]$$

$$D_{v_i} - column\ normalizing\ matrix\ for\ \overset{(i)}{V}\ , i \in [1, 3, \dots, 2n-1]$$

$$D_{h_i} - row\ normalizing\ matrix\ for \overset{(i)}{H}, i \in [0, 2, \dots, 2n-2]$$

$$D_{w_i} - column\ normalizing\ matrix\ for\ \overset{(i)}{W}, i \in [1, 3, \dots, 2n-1]$$

### In this paper forward procedure was applied only to matrix $V$

Since $W$ and $H$ are not known initially, in this paper we apply this procedure partially.

So only $V$ and $n$ were passed as input parameters and $(\overset{(1)}{V}, \overset{(2)}{V}, \dots, \overset{(2n)}{V})$ and $(D_{v_0}, D_{v_1}, \dots, D_{v_{2n-1}})$ were returned as an output.

### Introduce reverse Sinkhorn procedure after NMF solution

In this paper the NMF/deconvolution was performed in Sinkhorn transformed space with optimization function problem defined earlier

$$\min_{\substack{X \\ \Omega \\ D}} \left\| S \times \overset{\infty}{V}_{ss} \times R^T - \Omega \times D \times X \right\| \quad \substack{s.t.\ \overset{\infty}{W}_{fs} = S^T \times \Omega > 0 \\ \overset{\infty}{H}_{ss} = X \times R > 0}$$

Where $\overset{\infty}{V}_{ss}, \overset{\infty}{V}_{fs}$ – known row and column normalized matrices produced by Sinkhorn procedure, representing simplexes in new sample and space respectively.

$$\overset{\infty}{V}_{ss} = \overset{\infty}{W}_{ss} \times \overset{\infty}{H}_{ss}$$

$$\overset{\infty}{V}_{fs} = \overset{\infty}{W}_{fs} \times \overset{\infty}{H}_{fs}$$

$$\overset{\infty}{V}_{gs} = \overset{\infty}{V}_{ss} \times D_V = \frac{N}{M} \times \overset{\infty}{V}_{ss} = \underbrace{\overset{\infty}{W}_{ss} \times \overset{\infty}{D}_W}_{\overset{\infty}{W}_{gs}} \times \underbrace{\overset{\infty}{D}{}_W^{-1} \times \overset{\infty}{H}_{ss} \times D_V}_{\overset{\infty}{H}_{gs}}$$

Moreover, since after deconvolution we know matrices $R,\ S,\ X,\ \Omega$

We can get matrices $\overset{\infty}{W}_{fs}, \overset{\infty}{H}_{ss}$

$$\overset{\infty}{W}_{fs} = S^T \times \Omega$$

$$\overset{\infty}{H}_{ss} = X \times \mathrm{R}$$

Note that notation ∞ was used to point out the fact that Sinkhorn procedure was performed enough times to reach the convergence for both subsequences to reach the constant relation property.

$$\overset{\infty}{V}_{fs} = \frac{N}{M} \times \overset{\infty}{V}_{ss}$$

Let's fix the value for $n$ for which this property has already been reached and use this $n$ to represent these matrices in terms of Sinkhorn procedure.

$$\boxed{\overset{(2n)}{W} = \overset{\infty}{W_{fs}} = S^T \times \Omega - column\ normalized\ matrix\ (simplex\ in\ a\ features\ space)}$$

$$\boxed{\overset{(2n-1)}{H} = \overset{\infty}{H_{ss}} = X \times \mathrm{R} - row\ normalized\ matrix\ (simplex\ in\ a\ samples\ space)}$$

And now we will describe how to obtain original $W$ and $H$ from these matrices going from number $n$ to 0.

$$\boxed{\overset{(2n)}{W}, \overset{(2n)}{H} \Longrightarrow \overset{(2n-1)}{W}, \overset{(2n-1)}{H} \Longrightarrow \cdots \Longrightarrow \overset{(1)}{W}, \overset{(1)}{H} \Longrightarrow \overset{(0)}{W}, \overset{(0)}{H}}$$

For that we need to derive normalization matrices $D_{h_i}$ and $D_{w_i}$ for the previous step $\overset{(i)}{H}, \overset{(i)}{W}$ from the next step $\overset{(i+1)}{H}, \overset{(i+1)}{W}$.

### Derive column normalizing matrix for $\overset{(i)}{W}$

Let's note that matrix $D = \overset{\infty}{D_W^{-1}} = D_{w_{2n-1}}^{-1}$ is also known as part of the solution.

But $\forall i \in [1,3,\dots,2n-1]$ it is possible to restore column normalizing $D_{w_i}$ for $\overset{(i)}{W}$ using values of $\overset{(i+1)}{W}$ and row normalization property of $\overset{(i)}{W}$.

$$\forall i \in [1,3,\dots,2n-1]$$

$$D_{w_i} \times D_{w_i}^{-1} = \mathbb{I}_{K\times K}$$

Then

$$\overset{(i+1)}{W} \times D_{w_i}^{-1} = \overset{(i)}{W} \times D_{w_i} \times D_{w_i}^{-1} = \overset{(i)}{W}$$

Since $\overset{(i)}{W}$ is row normalized

$$\overset{(i)}{W} \times \mathbb{I}_K = \mathbb{I}_M$$

$$\overset{(i+1)}{W} \times D_{w_i}^{-1} \times \mathbb{I}_K = \mathbb{I}_M$$

Which means

$$\overset{(i+1)}{W} \times \begin{bmatrix} \frac{1}{(d_{w_i})_1} \\ \frac{1}{(d_{w_i})_2} \\ \vdots \\ \frac{1}{(d_{w_i})_K} \end{bmatrix} = \begin{bmatrix} 1 \\ 1 \\ \vdots \\ 1 \end{bmatrix}$$

Which means that $D_{w_i}^{-1}$ could be found as a solution of NNLS problem

$$\forall i \in [1,3,\ldots,2n-1]$$

$$\vec{d}_{w_i}^{-1} = \underset{x}{argmin} \left\| \begin{bmatrix} 1 \\ 1 \\ \vdots \\ 1 \end{bmatrix} - \overset{(i+1)}{W} \times x \right\|$$

$$D_{w_i}^{-1} = diag(\vec{d}_{w_i}^{-1})$$

### Derive row normalizing matrix for $\overset{(i)}{H}$

Similarly, $\forall i \in [2,4,\ldots,2n-2]$ it is possible to restore row normalizing $D_{h_i}$ for $\overset{(i)}{H}$ using values of $\overset{(i+1)}{H}$ and column normalization property of $\overset{(i)}{H}$.

$$\forall i \in [\,2,4,\ldots,2n-2]$$

$$D_{h_i}^{-1} \times D_{h_i} = D_{h_i} \times D_{h_i}^{-1} = \mathbb{I}_{K\times K}$$

$$D_{h_i}^{-1} \times \overset{(i+1)}{H} = D_{h_i}^{-1} \times D_{h_i} \times \overset{(i)}{H} = \overset{(i)}{H}$$

And in transposed form

$$\overset{(i+1)}{H^T} \times D_{h_i}^{-1} = \overset{(i)}{H^T} \times D_{h_i} \times D_{h_i}^{-1} = \overset{(i)}{H^T}$$

Since $\overset{(i)}{H^T}$ is row normalized, then:

$$\overset{(i)}{H^T} \times \mathbb{I}_K = \mathbb{I}_N$$

$$\overset{(i+1)}{H^T} \times D_{h_i}^{-1} \times \mathbb{I}_K = \mathbb{I}_N$$

Which means

$$\overset{(i+1)}{H^T} \times \begin{bmatrix} \frac{1}{(d_{h_i})_1} \\ \frac{1}{(d_{h_i})_2} \\ \vdots \\ \frac{1}{(d_{h_i})_K} \end{bmatrix} = \begin{bmatrix} 1 \\ 1 \\ \vdots \\ 1 \end{bmatrix}$$

Which means that $D_{h_i}^{-1}$ could be found as a solution of NNLS problem

$$\forall i \in [\,2, 4, \ldots, 2n-2]$$
$$\vec{d}_{h_i}^{-1} = \underset{x}{argmin}\left\|\begin{bmatrix}1\\1\\\vdots\\1\end{bmatrix} - \overset{(i+1)}{H^T} \times x\right\|$$
$$D_{h_i}^{-1} = diag(\vec{d}_{h_i}^{-1})$$

### Algorithmic formulation of reverse procedure

Knowing all these relations we can write a reverse Sinkhorn procedure.

As we already showed $\overset{(2n)}{W}$, $\overset{(2n-1)}{H}$, are already known from deconvolution solution.

We assume that: $D_{v_0}, D_{v_1}, \ldots, D_{v_{2n-1}}$ –already are already known from the forward procedure.

$D_{w_{2n-1}}^{-1} = D$ is also known, but if it is not, we could get it from using defined NNLS problem.

$$D_{w_{2n-1}}^{-1} = diag\left(\underset{x}{argmin}\left\|\begin{bmatrix}1\\1\\\vdots\\1\end{bmatrix} - \overset{(2n)}{W} \times x\right\|\right)$$

Using this we can calculate $\overset{(2n-1)}{W}$ and $\overset{(2n-1)}{H}$ (even if later is not known)

$$\overset{(2n)}{W} = \overset{(2n-1)}{W} \times D_{w_{2n-1}} \Leftrightarrow \overset{(2n-1)}{W} = \overset{(2n)}{W} \times D_{w_{2n-1}}^{-1}$$

$$\overset{(2n)}{H} = D_{w_{2n-1}}^{-1} \times \overset{(2n-1)}{H} \times D_{v_{2n-1}} \Leftrightarrow \overset{(2n-1)}{H} = D_{w_{2n-1}} \times \overset{(2n)}{H} \times D_{v_{2n-1}}^{-1}$$

On the other hand, as we described above

$$D_{h_{2n-2}}^{-1} = diag\left(\underset{x}{argmin}\left\|\begin{bmatrix}1\\1\\\vdots\\1\end{bmatrix} - \overset{(2n-1)}{H^T} \times x\right\|\right)$$

Which gives us all necessary elements to calculate $\overset{(2n-2)}{H}$ and $\overset{(2n-2)}{W}$

$$\overset{(2n-1)}{H} = D_{h_{2n-2}} \times \overset{(2n-2)}{H} \Leftrightarrow \overset{(2n-2)}{H} = D_{h_{2n-2}}^{-1} \times \overset{(2n-1)}{H}$$

$$\overset{(2n-1)}{W} = D_{v_{2n-2}} \times \overset{(2n-2)}{W} \times D_{h_{2n-2}}^{-1} \Leftrightarrow \overset{(2n-2)}{W} = D_{v_{2n-2}}^{-1} \times \overset{(2n-1)}{W} \times D_{h_{2n-2}}$$

We can continue this procedure until $n = 2$, defining the reverse Sinkhorn procedure as an iterative process of getting matrices $\overset{(1)}{W}$ and $\overset{(1)}{H}$ from matrices $\overset{(2n)}{W}$ and $\overset{(2n-1)}{H}$ respectively.

| Algorithm 2: Reverse Sinkhorn-Knopp procedure | |
|---|---|
| Input : | $\overset{(2n)}{V}, \overset{(2n)}{W}, \overset{(2n-1)}{H}$, n<br>$D_{v_0}, D_{v_1}, \dots, D_{v_{2n-1}}$ |
| Output : | $\overset{(1)}{V}, \overset{(1)}{W}, \overset{(1)}{H}$ |

For $i$ in $[2n-1, \dots 2]$:

$$D_{w_i}^{-1} = NNLS(\overset{(i+1)}{W})$$
$$\overset{(i)}{W} = \overset{(i+1)}{W} \times D_{w_i}^{-1}$$
$$if\ i < 2n-1:$$
$$\overset{(i)}{H} = D_{w_i} \times \overset{(i+1)}{H} \times D_{v_i}^{-1}$$
$$D_{h_{i-1}}^{-1} = NNLS(\overset{(i)}{H})$$
$$\overset{(i-1)}{H} = D_{h_{i-1}}^{-1} \times \overset{(i)}{H}$$
$$\overset{(i-1)}{W} = D_{v_i}^{-1} \times \overset{(i)}{W} \times D_{h_i}$$

### Obtain original matrices *W* and *H*

The algorithm defined above can yield matrices $\overset{(1)}{V}, \overset{(1)}{W}, \overset{(1)}{H}$. These matrices are all row-normalized.

We also know that

$$\overset{(1)}{H} = D_{h_0} \times \overset{(0)}{H}$$

$$\boxed{\overset{(0)}{V} = D_{v_0} \times \overset{(0)}{W} \times D_{h_0}^{-1} \times D_{h_0} \times \overset{(0)}{H} = D_{v_0}^{-1} \times \overset{(1)}{W} \times \overset{(1)}{H}}$$

Where $D_{h_0}$ is row-normalizing matrix for initial matrix $\overset{(0)}{H} = H$.

Next we explain how to obtain original matrix $H$ for two possible scenarios: first when we have normalization property for $H$ and then for general NMF problem

#### *Sum-to-one constrained (column normalized) matrix $H$*

Note that for mixture unmixing (deconvolution problem) we have specific property of column normalization for proportional matrix.

$$H^T \times \mathbb{I}_K = \mathbb{I}_N$$

This means that for this problem we can just continue original reverse procedure to obtain matrix $D_{h_0}^{-1}$ using known matrix $\overset{(1)}{H}$ using already described NNLS procedure:

$$D_{h_0}^{-1} = diag\left(\underset{x}{argmin}\left\|\begin{bmatrix}1\\1\\\vdots\\1\end{bmatrix} - \overset{(1)}{H} \times x\right\|\right)$$

Therefore $H$ and $W$ matrices can be obtained using this matrix and known $D_{v_0}$, $\overset{(1)}{H}$ and $\overset{(1)}{W}$:

$$\overset{(1)}{H} = D_{h_0} \times \overset{(0)}{H} \Rightarrow \overset{(0)}{H} = D_{h_0}^{-1} \times \overset{(1)}{H}$$

$$\overset{(1)}{W} = D_{v_0} \times \overset{(0)}{W} \times D_{h_0}^{-1} \Rightarrow \overset{(0)}{W} = D_{v_1}^{-1} \times \overset{(1)}{W} \times D_{h_1}$$

Note that in the case of gene expression deconvolution matrix $\overset{(0)}{H} = H$ is column normalized, which is guaranteed by this procedure.

*General matrix factorization problem*

Note that for general NMF problem we don't have any normalization requirement for the original matrix $H$ which makes it impossible to use this property for obtaining this matrix.

$$\overset{(1)}{H} = D_{h_0} \times \overset{(0)}{H}$$

$$\overset{(1)}{V} = \overset{(1)}{W} \times \overset{(1)}{H}$$

$$\boxed{\overset{(0)}{V} = D_{v_0} \times \overset{(0)}{W} \times D_{h_0}^{-1} \times D_{h_0} \times \overset{(0)}{H} = D_{v_0}^{-1} \times \overset{(1)}{W} \times \overset{(1)}{H}}$$

Without additional constraints for $\overset{(0)}{H}$ or $\overset{(0)}{W}$ the choice of $D_{h_0}$ could be arbitrary. Different values on the main diagonal will lead to different factorizations of the initial expression matrix.

However, all these factorizations are valid solutions for NMF problem.

Therefore, we are choosing identity matrix as $D_{h_0}$ and keeping row-normalized matrix $\overset{(1)}{H}$ defining the rest part of the equation as $\overset{(0)}{W}$, which still yields a valid factorization of the original matrix.

$$\boxed{\overset{(0)}{V} = \underbrace{D_{v_0}^{-1} \times \overset{(1)}{W}}_{\overset{(0)}{W}} \times \overset{(1)}{H}}$$

*Directions of the main components*

Note, that all factorizations will have the same directions of pure expression component vectors $\boldsymbol{W}$.

Since

$$\overset{(1)}{W} = D_{v_1} \times \overset{(0)}{W} \times D_{h_1}^{-1} = \begin{bmatrix} d_{v_{1_1}} & \cdots & 0 \\ \vdots & \ddots & \vdots \\ 0 & \cdots & d_{v_{1_M}} \end{bmatrix} \times \begin{bmatrix} w_{1,1} & \cdots & w_{1,K} \\ \vdots & \ddots & \vdots \\ w_{M,1} & \cdots & w_{M,K} \end{bmatrix} \times \begin{bmatrix} d_{h_{1_1}}^{-1} & \cdots & 0 \\ \vdots & \ddots & \vdots \\ 0 & \cdots & d_{h_{1_K}}^{-1} \end{bmatrix}$$

Then

$$\overset{(1)}{W} = \begin{bmatrix} d_{h_{1_1}}^{-1} d_{v_{1_1}} w_{1,1} & \cdots & d_{h_{1_K}}^{-1} d_{v_{1_1}} w_{1,K} \\ \vdots & \ddots & \vdots \\ d_{h_{1_1}}^{-1} d_{v_{1_M}} w_{M,1} & \cdots & d_{h_{1_K}}^{-1} d_{v_{1_M}} w_{M,K} \end{bmatrix}$$

Where each column could be written as

$$\overset{(1)}{w}_{*,i} = d_{h_{1_i}}^{-1} \begin{bmatrix} d_{v_{1_1}} w_{1,i} \\ \vdots \\ d_{v_{1_M}} w_{M,i} \end{bmatrix}$$

Which means that each column $i$ of matrix $\overset{(1)}{W}$ is a column $i$ of the matrix $D_{v_1} \times \overset{(0)}{W}$, scaled with the value $d_{h_{1_i}}^{-1}$. Here $\overset{(1)}{W}$ and $D_{v_1}$ are known matrices and $\overset{(0)}{W}$ is desired matrix we encoded in the initial problem.

#### *Decomposition of factorization into 3 matrices*

Since directions of main vectors defining the actual NMF solution are important, we can decompose the whole solution equation to have column-normalized matrix $W_{colnorm}$, which specifies directional components of the solution in samples space and row normalized matrix $H_{rownorm}$ which specifies directional components of the solution in features space. Whatever matrix is left in between will specify the relationship between these two spaces.

$$V = W \times H \Leftrightarrow V = W_{colnorm} \times \underbrace{D_w^{-1} \times D_h^{-1}}_{D} \times H_{rownorm}$$

Importantly such a decomposition allows for comparison between different NMF methods, since different algorithms typically return different scales of matrices $W$ and $H$

### Supplementary Note 12. The filtering preprocessing pipeline for gene expression data

Gene expression data matrix typically contain noise due to huge number of factors.

There is no way to make an efficient deconvolution algorithm without the procedure to remove noise in input data. In consequence, it is first required to apply some preprocessing pipeline. In this section, we will describe novel filtering approaches for input data genes and samples in deconvolution problem.

#### Removing non-informative genes.

Before we apply any technical filtering, we use a strict procedure to remove a subset of genes that we are not expecting to be part of any possible cell types. This subset includes.

1. non-coding genes,
2. open reading frames genes (C.orf)
3. genes without known symbols (LOC)

#### Elective filtering for housekeeping genes.

Input gene expression data will usually contain not only cell-type specific genes but also highly expressed genes that will be common for all cell types (housekeeping genes). So, to enhance the simplex structure in the data we recommend removing these genes although it's not critical for algorithm operations.

Since there is no clear definition of housekeeping genes we also removed genes which are similar to filtered genes in terms of KNN distance of expression profile vectors.

#### Filtering out cell cycle genes and ribosomal genes.

Another group of genes that could affect deconvolution results - cell cycle (GO:0007049) and ribosomal genes (RPL/RPS). Those genes are typically highly correlated and introduce additional linear components that are not relevant to cell types. Commonly, those genes should also be removed from input data because of possible overfitting.

#### Filtering out low-expressed and low-variable genes.

To identify low-expressed and low-variable genes we calculate the median absolute deviation (MAD) for each gene. MAD measures dispersion by finding the value where half of the data is closer to the median and half of the data is farther from the median than that value. Therefore, for low-expressed genes MAD value still will be significantly small. The MAD metric is also known to be less sensitive to outliers in the data which allows us to apply this filtering step at the beginning.

The filtering threshold is usually unique for each input dataset, but before it can be found, it is useful to first remove genes that have MAD=0, thereby removing genes that are not expressed or not changing expression in at least 50% of the samples.

To search for a suitable MAD filtering threshold, we must check its distribution. For common expression data, MAD distribution will be bimodal – the first mode will be around 0 value, corresponding to low-expressed and low-variable genes, and the second mode will be around some higher value specific to analyzed data. We put a threshold between two mode values and filter out genes close to the first mode, keeping only genes with high expression and variability, including cell type specific genes.

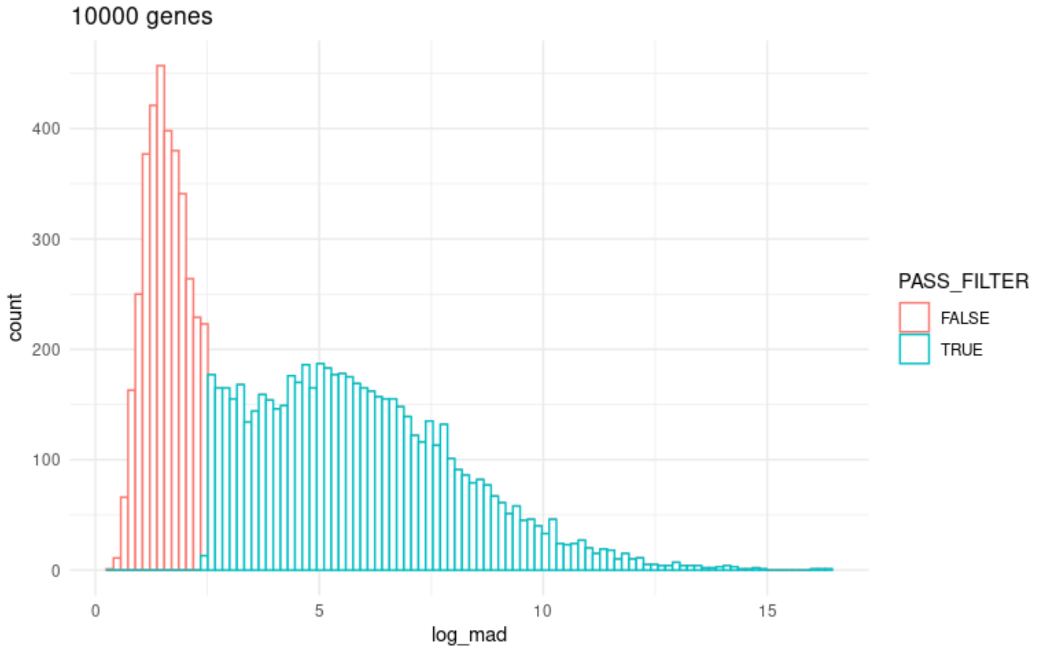

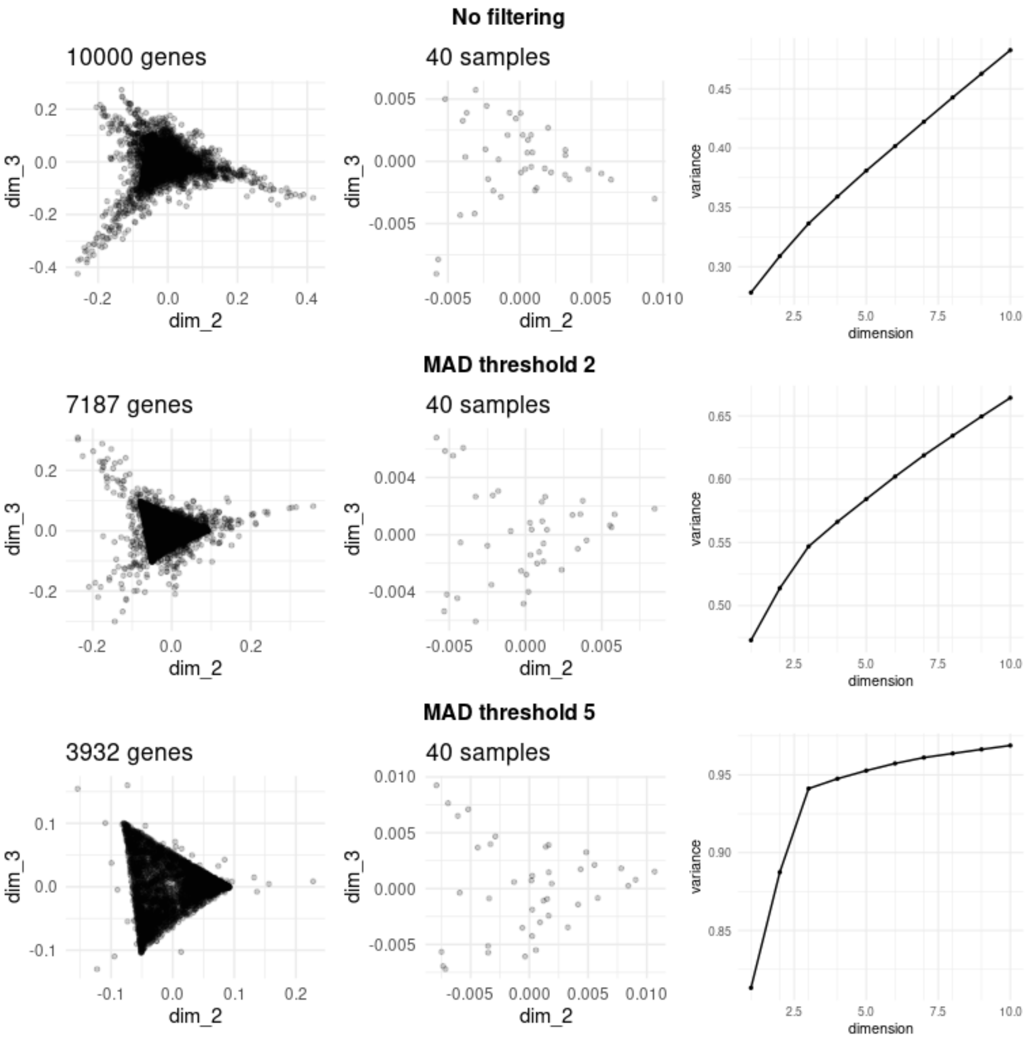


(Simulated data $sd = 4.0,\ M = 10000,\ N = 40,\ K = 3$)

You can always inspect the changes in SVD by looking at elbow plot history

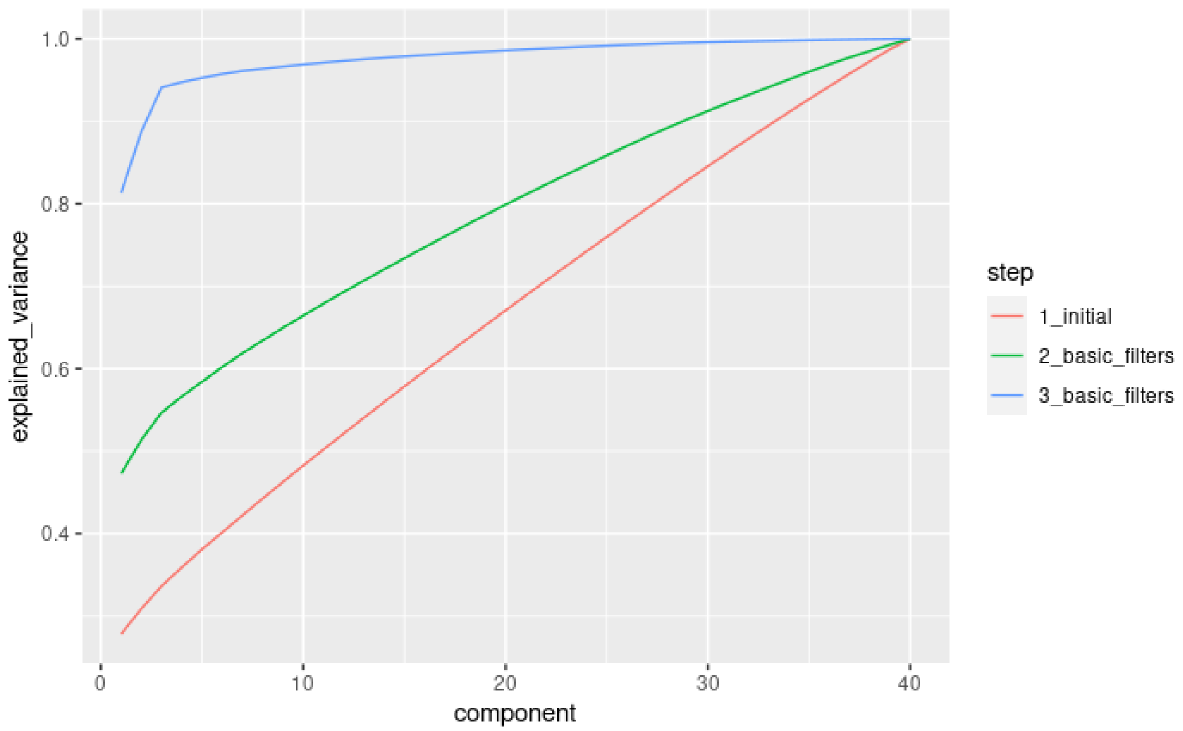


(Simulated data $sd = 4.0$, $M = 10000$, $N = 40$, $K = 3$, MAD=2 and MAD=5 filters applied sequentially)

### Denoising the data.

Another way to improve the projection optimization solution is to denoise the data by removing outlier genes and samples, which are far from the result hyperplane.

If we assume that we have expression data without noise, then we can guarantee that all points in genes' or samples' spaces will lay on some hyperplane in the projection space that will define desired simplex (simplex hyperplane). But it is not common to have no noise in the real experimental data. Moreover, PCA and SVD based approaches are sensitive to noise level. To demonstrate the effects of noise on the existence of hyperplane we modeled data with different noise levels.

One of the advantages of our approach is that it is possible to track the filtering process by examining the elbow plot of singular values. This plot illustrates the variance explained by singular vector components. In general, when the first $K$ components account for a higher proportion of the variance, it signifies that more of the data conforms to the $(K-1)$-dimensional simplex structure.

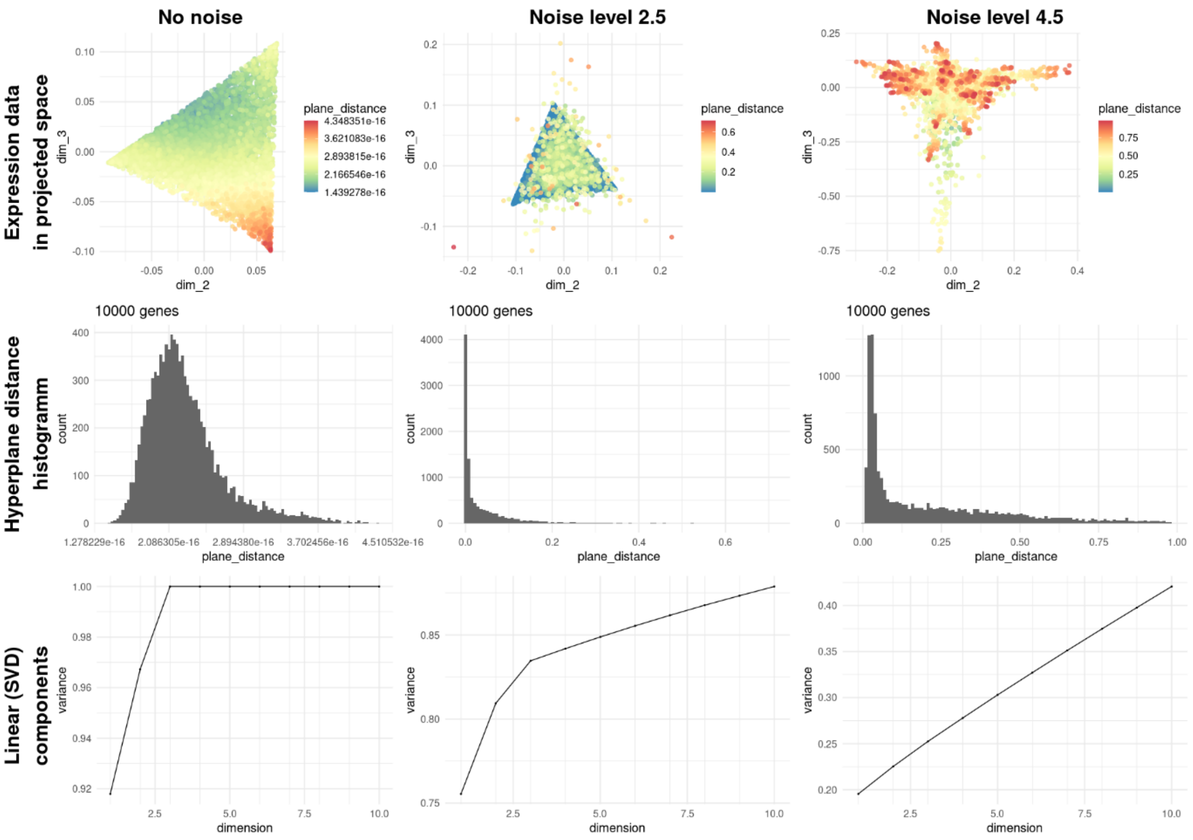


(Simulated data, different levels of noise $M = 10000,\ N = 40,\ K = 3$)

Provided examples show that even in small noise levels some genes are not located on the simplex hyperplane thereby if they will be not filtered, will affect deconvolution error. Also, at higher levels of noise, we can see additional linear components that are not representing real cell types.

Deconvolution method proposed in gives us a systematic way to think about denoising the data.

We can estimate how good a point was projected on a hyperplane by estimating the cost function we used to find projection vectors $R(S)$ for the given point. This cost function represents the distance between original data points and the points which were recovered from projection.

$$\left\| \overset{\infty}{V}{}_{ss}^{T} - R^{T} \times R \times \overset{\infty}{V}{}_{ss}^{T} \right\| \quad \left\| \overset{\infty}{V}_{gs} - S^{T} \times S \times \overset{\infty}{V}_{gs} \right\|$$

If the projection that we found is right, we will end up with some hyperplane that will be close to a simplex hyperplane (and therefore the cost function will have smaller values). But typically, the presence of noise in the input data will negatively affect projection, causing some points to be outliers in terms of recoverability.

For such points we can estimate residual noise of projection using Euclidean distance for individual point.

$$\forall i \in [1, M]\ distance_{gene_i} = \sqrt{\sum_{j=1}^{N} \left( \overset{\infty}{V}{}_{ss\ j,i}^{T} - R^{T} \times R \times \overset{\infty}{V}{}_{ss\ j,i}^{T} \right)^{2}}$$

$$\forall j \in [1, N]\ distance_{sample_j} = \sqrt{\sum_{i=1}^{M} \left( \overset{\infty}{V}_{gs_{i,j}} - S^T \times S \times \overset{\infty}{V}_{ss_{i,j}} \right)^2}$$

By filtering the most distant elements (with the highest distance to hyperplane) we can remove the noise that affects projection accuracy.

In this work we used manual thresholds to remove outlier genes.

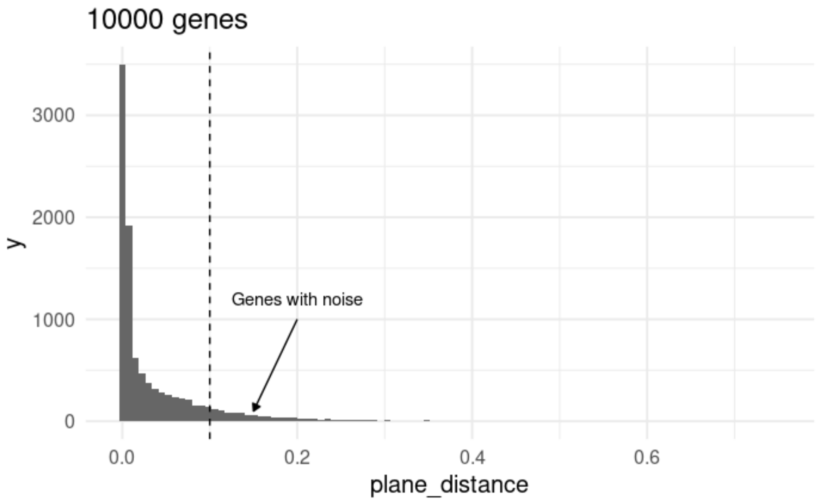


(Simulated data, Distance to plane histogram, $sd = 2.5, M = 10000,\ N = 40,\ K = 3$)

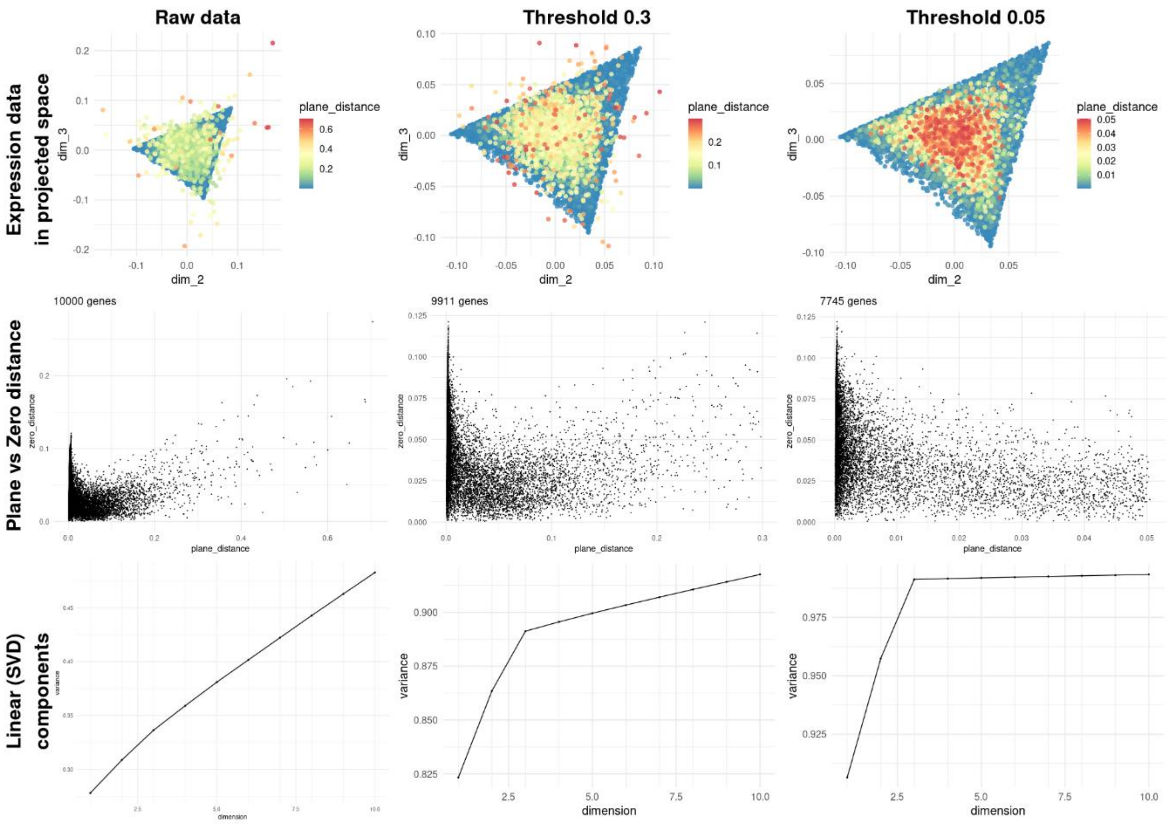


(Simulated data, different noise cutoff levels, $sd = 2.5, M = 10000,\ N = 40,\ K = 3$)
